# Divergent sensory and immune gene evolution in sea turtles with contrasting demographic and life histories

**DOI:** 10.1101/2022.01.10.475373

**Authors:** Blair P. Bentley, Tomás Carrasco-Valenzuela, Elisa K. S. Ramos, Harvinder Pawar, Larissa Souza Arantes, Alana Alexander, Shreya M. Banerjee, Patrick Masterson, Martin Kuhlwilm, Martin Pippel, Jacquelyn Mountcastle, Bettina Haase, Marcela Uliano-Silva, Giulio Formenti, Kerstin Howe, William Chow, Alan Tracey, Ying Sims, Sarah Pelan, Jonathan Wood, Kelsey Yetsko, Justin R. Perrault, Kelly Stewart, Scott R. Benson, Yaniv Levy, Erica V. Todd, H. Bradley Shaffer, Peter Scott, Brian T. Henen, Robert W. Murphy, David W. Mohr, Alan F. Scott, David J. Duffy, Neil J. Gemmell, Alexander Suh, Sylke Winkler, Françoise Thibaud-Nissen, Mariana F. Nery, Tomas Marques-Bonet, Agostinho Antunes, Yaron Tikochinski, Peter H. Dutton, Olivier Fedrigo, Eugene W. Myers, Erich D. Jarvis, Camila J. Mazzoni, Lisa M. Komoroske

## Abstract

Sea turtles represent an ancient lineage of marine vertebrates that evolved from terrestrial ancestors over 100 MYA, yet the genomic basis of the unique physiological and ecological traits enabling these species to thrive in diverse marine habitats remains largely unknown. Additionally, many populations have drastically declined due to anthropogenic activities over the past two centuries, and their recovery is a high global conservation priority. We generated and analyzed high-quality reference genomes for the leatherback *(Dermochelys coriacea)* and green *(Chelonia mydas)* turtles, representing the two extant sea turtle families. These genomes are highly syntenic and homologous, but localized regions of non-collinearity were associated with higher copy numbers of immune, zinc-finger, and olfactory receptor (OR) genes in green turtles, with ORs related to waterborne odorants greatly expanded in green turtles. Our findings suggest that divergent evolution of these key gene families may underlie immunological and sensory adaptations assisting navigation, occupancy of neritic versus pelagic environments, and diet specialization. Reduced collinearity was especially prevalent in microchromosomes, with greater gene content, heterozygosity, and genetic distances between species, supporting their critical role in vertebrate evolutionary adaptation. Finally, diversity and demographic histories starkly contrasted between species, indicating that leatherback turtles have had a low yet stable effective population size, exhibit extremely low diversity compared to other reptiles, and harbor a higher genetic load compared to green turtles, reinforcing concern over their persistence under future climate scenarios. These genomes provide invaluable resources for advancing our understanding of evolution and conservation best practices in an imperiled vertebrate lineage.

**Statement of significance:** Sea turtle populations have undergone recent global declines. We analyzed *de novo* assembled genomes for both extant sea turtle families through the Vertebrate Genomes Project to inform their conservation and evolutionary biology. These highly conserved genomes were differentiated by localized gene-rich regions of divergence, particularly within microchromosomes, suggesting that these genomic elements play key functional roles in the evolution of sea turtles and possibly other vertebrates. We further demonstrate that dissimilar evolutionary histories impact standing genomic diversity and genetic load, and are critical to consider when using these metrics to assess adaptive potential and extinction risk. Our results also demonstrate how reference genome quality impacts inferences of comparative and conservation genomics analyses that need to be considered in their application.

## Introduction

Sea turtles recolonized marine environments over 100 MYA (1, 2) and are now one of the most widely distributed vertebrate groups on the planet (3). Leatherback turtles *(Dermochelys coriacea)* represent the only remaining species of the family Dermochelyidae, which diverged from the Cheloniidae (hard-shelled sea turtles) about 60 MYA (4). Unique morphological (Fig. 1a) and physiological traits allow leatherback turtles to exploit cool, highly productive pelagic habitats (5, 6), while green turtles (*Chelonia mydas*) and other hard-shelled species largely inhabit warmer nearshore habitats following an early pelagic life stage. Most previous research in this group has focused on organismal and ecological adaptations (7), but the genomic basis of traits that differentiate or unite these species is not well understood.

**Fig. 1.**
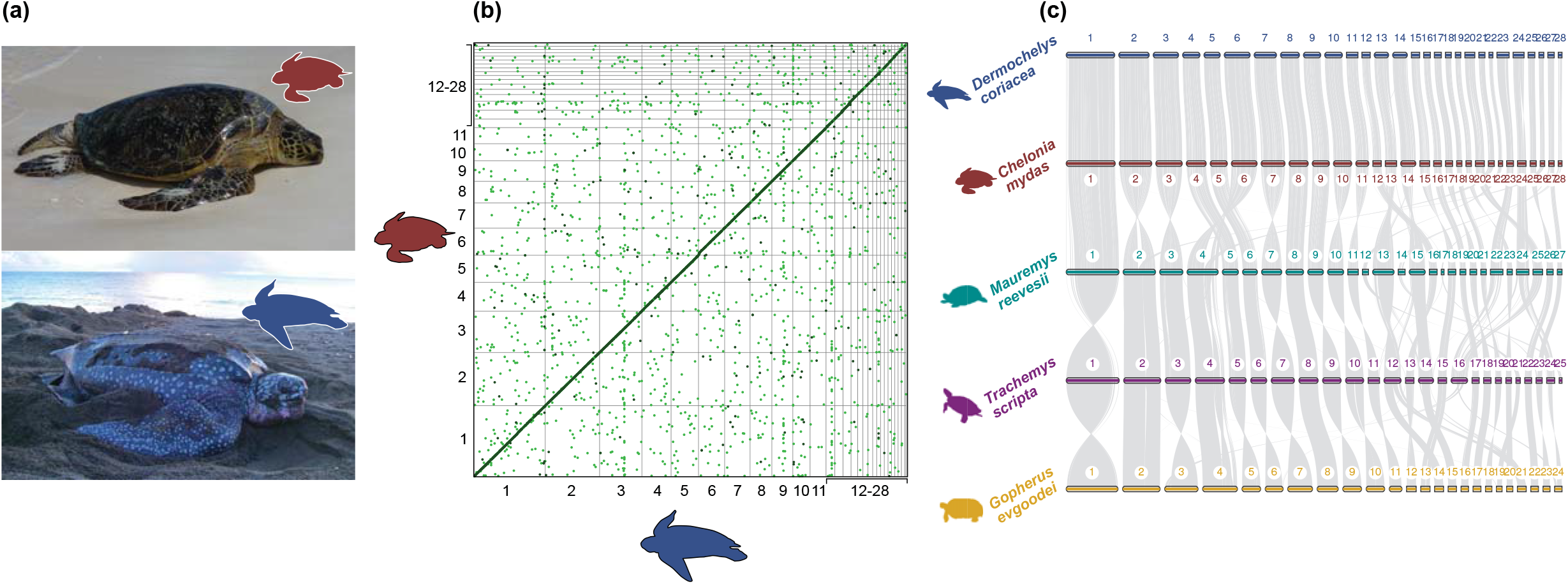
**(a)** Photographs of green turtle *(Chelonia mydas);* photo credit: NOAA NMFS PIFSC under USFWS Permit #TE-72088A-3, and leatherback turtle *(Dermochelys coriacea);* photo credit: Ricardo Tapilatu. **(b)** Dot plot showing regions with an identity greater than 0.5 across the entire genomes of green (red) and leatherback (blue) turtles. **(c)** Gene synteny and collinearity per chromosome among five species of turtles: leatherback turtle (blue), green turtle (red), Chinese pond turtle (*Mauremys reevesii*; green), pond slider turtle (*Trachemys scripta*; purple) and Goode’s thornscrub tortoise *(Gopherus evgoodei;* yellow). Each bar represents chromosomes with respective numbers and gray lines represent homolog gene connections among species.

Anthropogenic pressures have caused substantial population declines in sea turtles, with contemporary populations currently representing mere fractions of their historical abundances (8, 9). Although sea turtles spend most of their life in the ocean, they also exhibit long-distance migrations to natal rookeries for terrestrial reproduction (7, 10, 11). Consequently, they are threatened by human activities in both terrestrial and marine environments, including direct harvest of meat and eggs (12), fisheries bycatch (13), coastal development (14, 15), pollution (16), disease (17), and climate change (18, 19), which is exacerbated by their temperature-dependent mechanism of sex determination (TSD) altering population dynamics (20, 21). The IUCN lists most sea turtle species as vulnerable or endangered, and while decades of conservation efforts have fueled positive trends for some populations (22), others continue to decline (23). In particular, leatherback turtles have undergone extensive declines (>95% in some populations) over the last century (24–27), including the extirpation of the Malaysian nesting population (28). Leatherback turtle recovery is also impeded by relatively low hatching success compared to other sea turtle species (29). In contrast, many green turtle populations have recently increased following conservation actions (22), but their continued recovery remains threatened by anthropogenic activities and high incidence of the neoplastic disease fibropapillomatosis (FP), a likely viral-mediated tumor disease that disproportionately impacts this species (30).

Genomic data have been instrumental in advancing understanding of species’ evolutionary histories and ecological adaptations (31–33), and providing critical information for conservation management (34–37). However, this research has been hampered in taxa where genomic resources remain limited. In particular, the lack of high-quality reference genomes, which are essential for accurate comparative evolutionary analyses (38, 39) and robust estimates of a range of metrics to inform conservation biology such as inbreeding, hybridization, disease susceptibility, genetic load, and adaptation (36, 40, 41), impede this work in threatened species. An early draft genome for the green turtle was assembled almost a decade ago (42), and provided important insights into turtle evolution. However, errors, gaps, mis-assemblies, and fragmentation in draft genomes can lead to spurious inferences, potentially masking signals of interest (38, 43) and impeding effective management strategies (41). Well-annotated, chromosomal-level reference genomes can resolve these issues, improving our understanding of the genomic underpinnings of ecological and evolutionary adaptations (39, 44). For example, high-quality genomes with accurate annotations have enabled examination of gene changes associated with recolonization of the marine environment by terrestrial vertebrates, including the loss of olfactory receptor (OR) gene families (32, 45). Comparative genomic analyses have also demonstrated adaptive diversity in genes underlying reptilian immunity (46), and high-quality genomes have provided key insights into mammalian disease susceptibility (33, 47, 48). Such investigations are critical for sea turtles, with diseases such as FP adversely impacting populations across the globe (30), and information on immune genes is needed for devising effective conservation strategies (49).

We assembled chromosome-level reference genomes for leatherback and green turtles as part of the Vertebrate Genomes Project (VGP), and leveraged these resources to address questions centered around sea turtle evolutionary history and conservation. Specifically, we provide insights into the genomic underpinnings of phenotypic traits that separate and unite these two species by examining genome synteny and regions of divergence. Given the contrasting recent population trends of these two species, we additionally used whole genome re-sequencing (WGR) data of individuals representative of global populations to compare key conservation-relevant metrics, including patterns of diversity and deleterious variants across the genomes, and reconstructed demographic histories to inform assessments of future vulnerability. These genomes represent two of the most contiguous reptilian genomes assembled to date, and our results provide a foundation for further hypothesis-driven investigations into the evolutionary adaptation and conservation of this imperiled vertebrate lineage.

## Results

### Genome quality

Reference genomes for the leatherback and green turtles were generated using four genomic technologies following the VGP pipeline v1.6 (39), with minor modifications (see Methods). A total of 100% of the leatherback and 99.8% of the green turtle assembled sequences were placeable within chromosomes. The assembled genomes were near full-length (~2.1 GB), with annotations of all 28 known chromosomes for both species, composed of 11 macrochromosomes (>50 Mb) and 17 microchromosomes (<50 Mb) (Tables 1 & S1, Fig. S1). These genomes are among the highest quality genomes assembled for non-avian reptiles to date in terms of both contiguity and completeness (Table S3), with the leatherback turtle assembly representing the first reptile genome where all scaffolds were assigned to chromosomes. Scaffold N50s were high for both genomes (Table 1). We annotated 18,775 protein-coding genes in the leatherback and 19,752 in the green turtle genomes (see below for analysis of these gene differences). For the leatherback and green turtles, 96.9% and 97.5% of these genes were supported at >95% of their length from experimental evidence and/or high-quality protein models from related species (see Methods). The numbers of protein-coding genes are within the range of other reptiles (Table S3) and include 97.7% and 98.2% complete BUSCO copies for leatherback and green turtles based on Sauropsida models (50), which are similar or higher than all other assembled reptilian genomes to date (Fig. S2).

**Table 1.** Quality statistics for the genome assemblies and annotations for leatherback *(Dermochelys coriacea)* and green *(Chelonia mydas)* turtles.

|  | Leatherback turtle<br>(*Dermochelys coriacea*) | Green turtle<br>(*Chelonia mydas*) |
| --- | --- | --- |
| Genome ID | rDerCor1 | rCheMyd1 |
| Assembly accession | GCA_009764565.3 | GCA_015237465.1 |
| Assembly level | Chromosome | Chromosome |
| Total genome length (bp) | 2,164,762,090 | 2,134,358,617 |
| Contig N50 (bp) | 7,029,801 | 39,415,510 |
| Scaffold N50 (bp) | 137,568,771 | 134,428,053 |
| Number of scaffolds | 40 | 92 |
| Number of chromosomes | 28 | 28 |
| Quality Value (QV) | 38.9 | 47.7 |
| Annotated protein-coding genes | 18,775 | 19,752 |

| BUSCO category | Assembly | Annotation | Assembly | Annotation |
| --- | --- | --- | --- | --- |
| Complete genes | 91.6% | 97.2 | 94.2% | 97.9% |
| Complete + fragmented | 95.4% | 97.7 | 96.7% | 98.2% |
| Missing | 4.10% | 1.3% | 2.8% | 0.7% |
| Duplicated | 0.5% | 1.0% | 0.5% | 1.1% |

### Genome architecture

Despite diverging over 60 MYA (4), leatherback and green turtles show extremely high genome synteny and collinearity (Figs. 1b,c, S6, S7), with Progressive CACTUS revealing 95% sequence identity across the length of the genomes (Table S5). After multiple rounds of manual curation to correct artifacts of mis-assemblies, few large structural rearrangements between the two species remained, including inversions of up to 7 Mb on chromosomes 12, 13, 24 and 28 (Fig. S6). The high collinearity between species included near-complete end-to-end contiguous synteny for nine of 28 chromosomes (Fig. S6). The remaining 19 chromosomes exhibited at least one small region of reduced collinearity (RRC) between the species, with RRCs representing a total of ~83.4 Mb (~3.9%) and ~110.5 Mb (~5.2%) of the leatherback and green turtle genome lengths, respectively. Eight chromosomes exhibited small RRCs (0.1–3 Mb), and 11 contained RRCs that were between 3–18 Mb in length (Figs. 2a-d & Table S6). Analyses of coding regions revealed a similar pattern of strong collinearity between the two species (Figs. 1c, S6), particularly within the macrochromosomes, which contain more than 80% of the total length of the genomes. The two genomes also displayed similar percentages of repetitive elements (REs), which were almost exclusively transposable elements (TEs) and unclassified repeats (Fig. S8). The landscape of TE superfamily composition over evolutionary time was also similar between the two species, with the exception of REs with low Kimura values (<5%) which appeared at higher frequency in the leatherback turtle genome (see Supplementary Appendix 1 for full analyses).

**Fig. 2.**
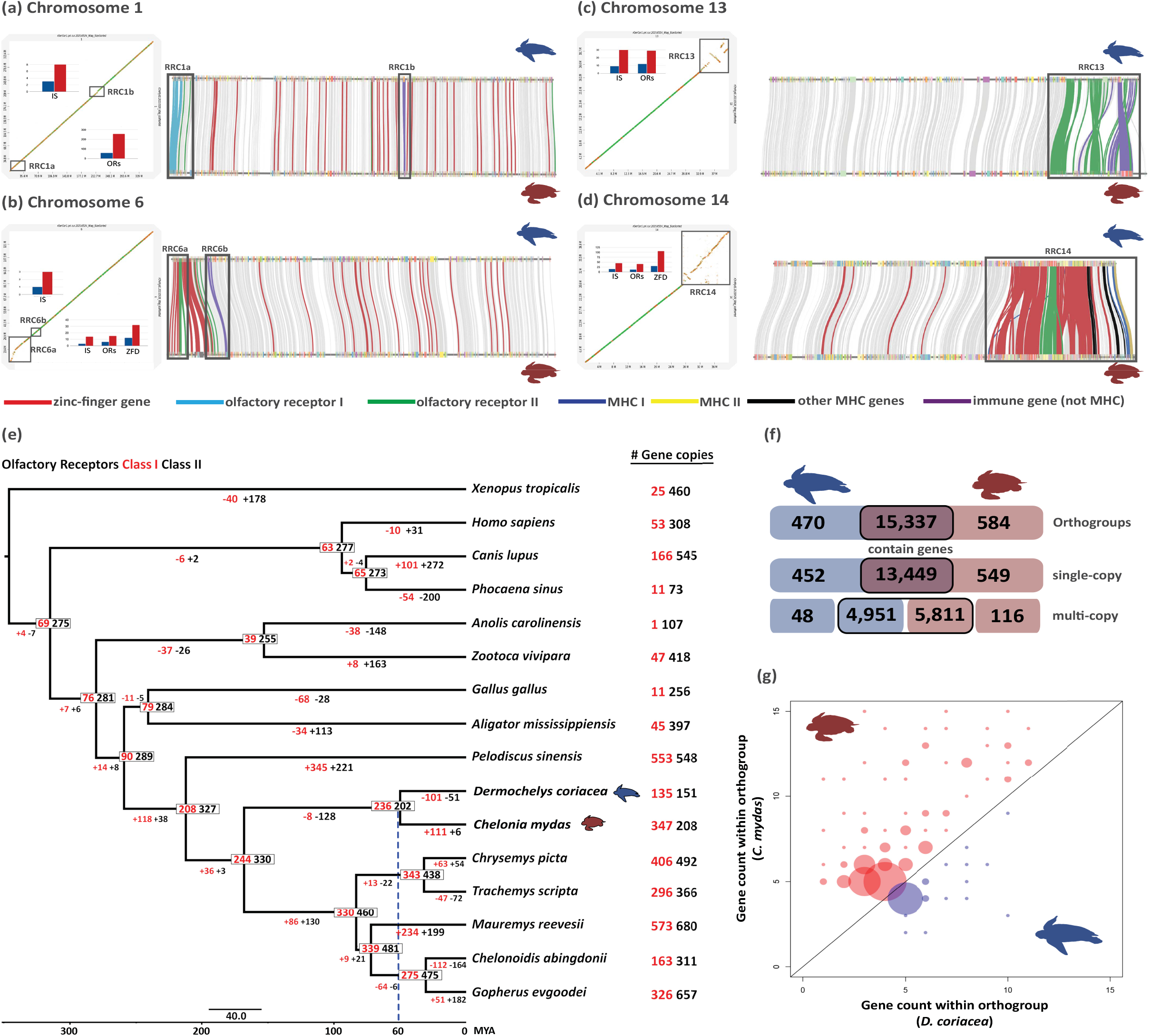
**(a-d)** Dotplots (identity values as color; dark green=1-0.75, green=0.75-0.5, orange=0.5-0.25 and yellow=0.25-0) depicting four of the regions with reduced collinearity (RRC) identified within chromosomes and associated with higher copy numbers of immune system (IS), olfactory receptor (ORs), or zinc finger domain genes (ZFD) in the green turtle *(Chelonia mydas)* relative to leatherback *(Dermochelys coriacea)* turtle (see also Fig. S3, Tables S3 & S5 for full details of all RRCs). Positions of each RRC are marked with gray squares on the dot plots (left; with *D. coriacea* on the X-axes and *C. mydas* on the Y-axes) and gene collinearity maps (right) for each chromosome highlighting the connections among specific gene families in different colors. **(e)** Gene family evolution of olfactory receptors Class I (red) and Class II (black) for amniote phylogeny. Gene numbers are presented on the nodes and gain/loss along each branch are presented below branches. Small scale bar represents substitutions/site and big scale bar represents divergence times (MA). The blue dashed line shows the estimated divergence between the two sea turtle families. **(f)** Number of unique and shared orthogroups and single and multi-copy genes between the two sea turtles (coding genes including genes with rearrangement). The boxes outlined in black denote shared orthogroups, with the higher multi-copy in the green turtle due to greater gene copies within orthogroups. **(g)** Comparison of gene counts between both species per multigenic orthogroup, depicting only those orthogroups where both species have different numbers of genes and a minimum number of five genes for one of the species. Bubbles above the diagonal represent higher counts for the green turtle and below for the leatherback turtle. The size of the bubbles represents the number of orthogroups with the same gene count combination.

### Gene families and gene functional analysis

Gene function analysis of localized RRCs revealed that most contained genes with higher copy numbers in the green turtle compared to the leatherback (Fig. 2a-d, Table S6). Nineteen chromosomes had RRCs with higher gene copy numbers in the green turtle, and of these, ten contained genes associated with immune system, olfactory reception and/or zinc-finger protein coding genes. Many of the same gene families were also detected as high-diversity exonic regions *via* separate, independent analyses (see Supplemental Appendix 1), reinforcing their importance in the divergent evolution of these species. In addition to localized RRCs, higher gene copy numbers in the green turtle occurred in many gene orthologous groups (orthogroups) across the entire genome, and generally in variable multicopy genes (Fig. 2f, g). Copy number variation accounted for most of the nearly one thousand more genes annotated in the green turtle genome relative to the leatherback (Fig. 2f, g; Table 1). We detected no evidence of collapsed multicopy genes in the leatherback turtle assembly across multiple analyses (see Methods; Table S7), supporting this as a biological signal rather than technical artifact of the assemblies.

Olfactory receptors (ORs) represented the largest orthogroups in both genomes, and differences in copy numbers were connected to many of the identified RRCs. All OR Class I genes were clustered at the beginning of chromosome 1, and the green turtle had higher copy numbers in this region (Fig. 2a-d). This area also contained a cluster of OR Class I genes in at least three additional testudinid species (Fig. S10), and is the only divergent region across the very large chromosome 1 in the turtles analyzed. In contrast, OR Class II genes were spread across several chromosomes in both sea turtle species, with higher copy numbers again in the green turtle found within RRCs (Fig. 2b-d). The instability and rapid evolution of OR gene numbers in turtles is further illustrated in the expansion-contraction analysis of orthogroups (Fig. 2e, Table S10a-d), which showed that OR Class I genes underwent a modest contraction in the ancestral sea turtle lineage, followed by an expansion in the green turtle but a further contraction in the leatherback turtle. Similar trends were detected for OR Class II genes, but with a greater magnitude of contraction in the ancestral sea turtle lineage followed by a further contraction for the leatherback turtle and only a small expansion for the green turtle (Fig. 2e).

Another important RRC (RRC14) encompassed the major histocompatibility complex (MHC), which plays a critical role in vertebrate immunity and is particularly relevant to sea turtle conservation due to the threat of FP and other diseases (32). In addition to the MHC region, this RRC includes several copies of OR Class II genes, zinc-finger protein coding genes and other genes involved with immunity, such as butyrophilin subfamily members and killer cell lectin-like receptors (Fig. 2d, Table S6). Invariably, the green turtle carried higher numbers of all the multicopy genes present in RRC14. RRCs on other chromosomes similarly showed increased levels of zinc-finger protein genes in the green turtle, including the RRCs labeled 6A, 11A, 14A, and 28 (Table S6). In particular, zinc-finger protein genes were highly prevalent on chromosomes 14 and 28 in both sea turtles, representing more than 50% of all the protein domains present on these chromosomes (Fig. S11). Finally, all but three genes with known roles in TSD in reptiles (Table S11) were located as single-copy genes within both sea turtle genomes, with homologous copies located in the same region of the chromosomes in both species (see Supplementary Appendix I for full analyses).

### Macro and microchromosomes

Microchromosomes contained significantly higher proportions of genes than macrochromosomes (Fig. 3a,b; green turtle: F_(2,25)_=16.46, p < 0.01; leatherback turtle: F_(2,25)_=16.35, p < 0.01), and gene content was strongly positively correlated with GC content (Fig. S13; green turtle R^2^ = 0.81, p < 0.01; leatherback turtle R^2^ = 0.87, p < 0.01). These patterns were particularly apparent in small (<20 Mb) microchromosomes, where GC content reached 50%, compared to the 43-44% genome-wide averages. Within chromosome groups, larger proportions of multicopy genes were generally associated with higher total gene counts (green turtle: R^2^=0.84, p < 0.01; leatherback turtle: R^2^=0.92, p < 0.01), and chromosomes with the highest multicopy genes numbers have increased proportions of RRCs (Fig. 3a,b; green turtle: R^2^=0.69, p < 0.01; leatherback turtle: R^2^=0.81, p < 0.01).

**Fig. 3.**
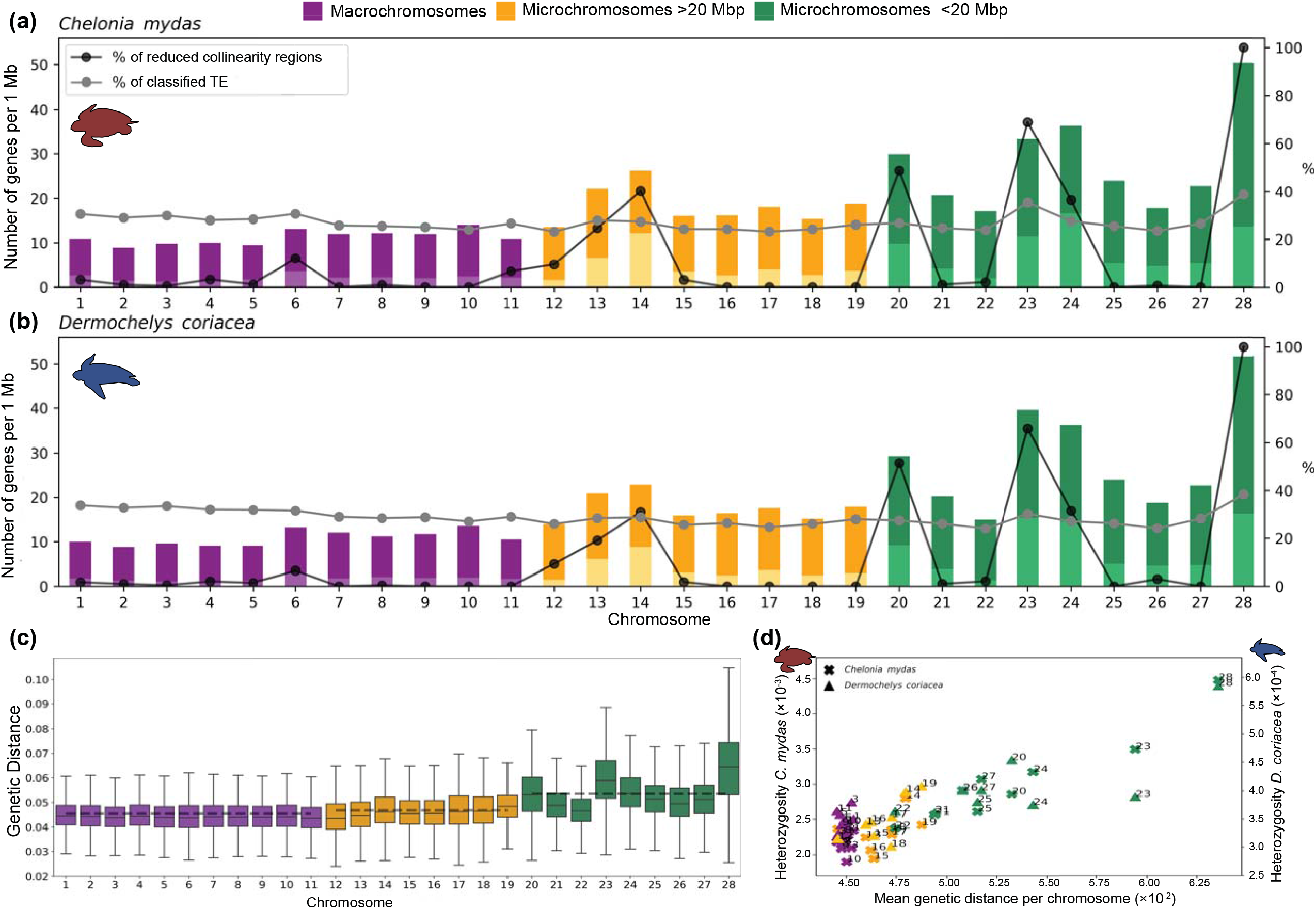
Number of genes, genetic distance between species and heterozygosity within species in macrochromosomes, small (<20 Mb) and intermediate (>20 Mb) microchromosomes. **(a)** Relation between the number of genes, percentage of reduced collinearity regions (RRCs), and classified TE per chromosome for the green *(Chelonia mydas)* and **(b)** leatherback *(Dermochelys coriacea)* turtles. Dark colors indicate the total number of genes and light colors indicate the number of multicopy genes. **(c)** Average genetic distance between green and leatherback turtles per chromosome. **(d)** Relation between genetic distance and heterozygosity per chromosome for each species.

Mean genetic distances for single-copy regions between the two sea turtles were also higher in small microchromosomes (0.053) compared to both intermediate (>20 Mb) microchromosomes (0.047), and macrochromosomes (0.045) (Fig. 3c; F_(2,25)_=21.98, p < 0.01). However, examination of intermediate microchromosome and macrochromosome RRCs revealed elevated genetic distances in these regions that approached the values observed in small microchromosomes (Table S12). Genetic distances were also significantly positively correlated with heterozygosity (green turtle: R^2^=0.97, p < 0.01; leatherback turtle R^2^=0.97, p < 0 .01), which was significantly higher in small microchromosomes for both species (Fig. 3d; green turtle: F(_2_,_25_)=15.72, p < 0.01; leatherback turtle: F_(2,25)_=5.09, p < 0.05).

### Genome diversity

Genome-wide nucleotide diversity was almost a magnitude of order lower in leatherback compared to green turtles (mean repeat masked π = 2.86×10^-4^ and 2.46×10^-3^, respectively; t_(5.52)_ = 36.9, p < 0.001; Figs. 4a, S15-17, Table S14). Despite having largely similar gene content identified in the annotation, this strong pattern was also observed in coding regions (Fig 4a.; t_(5.52)_ = 37.7, p < 0.001), such that leatherback turtles possess much less standing functional variation, which may impact their adaptive capacity to future novel environmental conditions. The strikingly low genomic diversity of leatherback turtles is also less than almost all other reptile species examined (Fig S19; but see (51)), including *Chelonoidis abingdonii*, where low diversity has been considered a contributing factor to their extinction (52). In contrast, the genomic diversity of the green turtle fell in the mid-range for reptiles, as well as for mammals examined using similar methods (53, 54). Finally, within both species, heterozygosity was lower in coding regions (mean π = 2.77×10^-4^ and 2.18×10^-3^ for leatherback and green turtles; Fig. 4a) relative to non-coding regions (mean π = 3.18×10^-4^ and 2.64×10^-3^; leatherbacks: [t_(4)_ = −8.9, p < 0.01] and greens: [t_(5)_ = −30.9, p < 0.01]), as expected from selection pressures driving higher sequence conservation in these functional genomic regions.

**Fig. 4.**
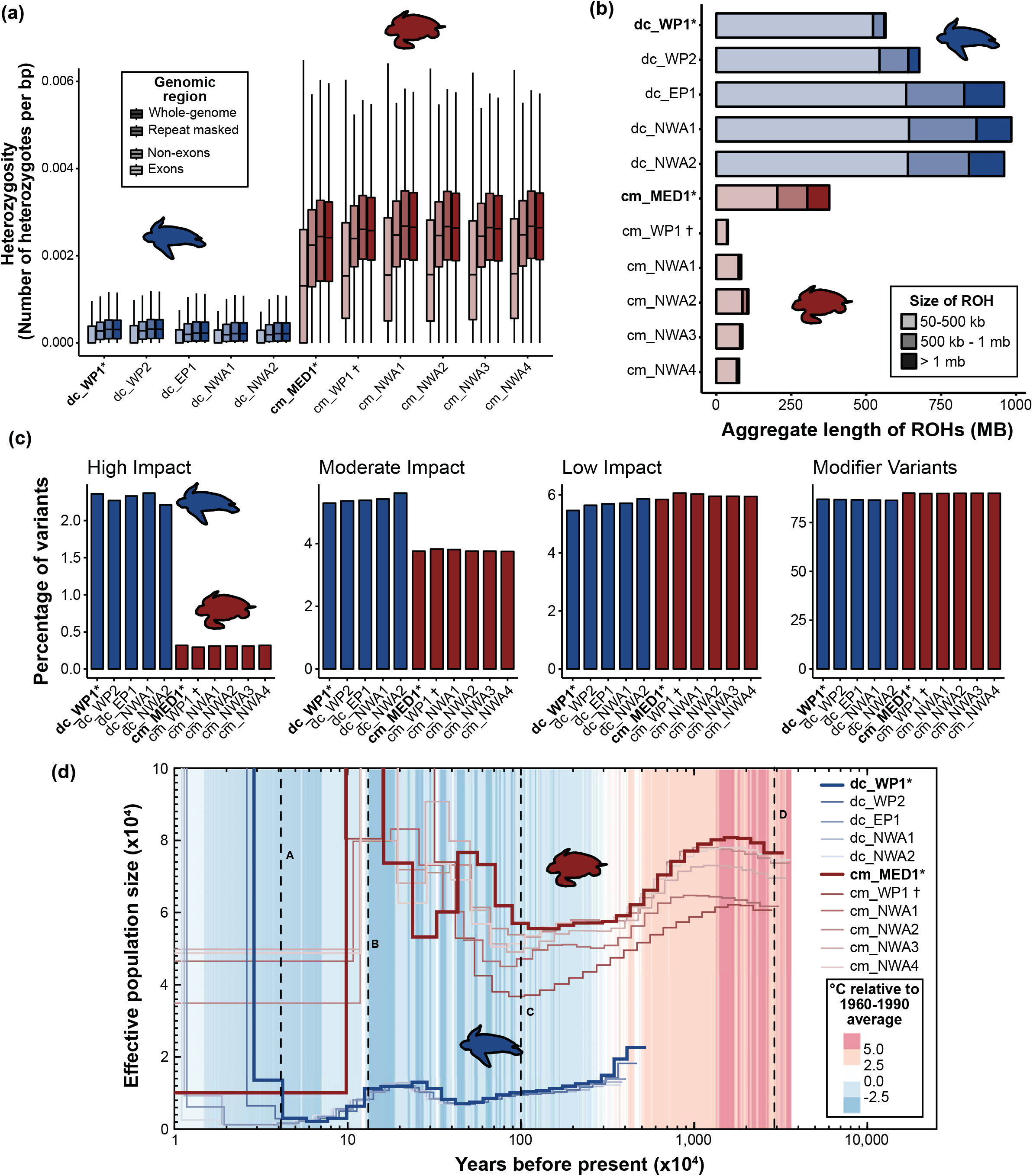
Data is presented for the leatherback *(Dermochelys coriacea;* blue) and green *(Chelonia mydas,* red) turtle genomes, including reference individuals for both species (*), and the individual used to generate the draft genome (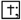 Wang et al. 2013). (a) estimates of heterozygosity across the genome calculated with 100 Kb non-overlapping windows for the entire genome, repeat-masked genome, exons and non-exon regions, with outliers removed. (b) accumulated lengths of runs of homozygosity (ROH). (c) predicted impacts of variants from within coding regions. (d) Pairwise sequential Markovian coalescent plot (PSMC) of demographic history of both species using a mutation rate of 1.2×10^-8^ and generation time of 30 years, overlayed with temperature. Letters indicating portions of the PSMC curves (A-D) are geological events referred to in the main text and Supplementary Appendix I.

### Runs of homozygosity (ROH)

In addition to lower genome-wide heterozygosity, leatherbacks had a greater total number of ROHs (>50 Kb) than green turtles (mean N_ROH_ = 4,510 and 829, respectively), as well as a greater total aggregate length of the genome in ROH (range =26.1 −45.5% in leatherback turtles; 1.8 – 17.7% in green turtles). The mean length of ROHs was also significantly higher in leatherback (L_ROH_ = 183.9 Kb) compared to green turtles (L_ROH_ = 154.9 Kb) (t_(7429.4)_ = −8.85, p < 0.01). Length distribution breakdown showed that leatherbacks have a higher aggregate length of all categories of ROHs relative to the green turtles (Figs. 4b, S22). Short ROHs (50-500 Kb) had the highest total aggregate length in leatherbacks, with a mean aggregate length of 597 Mb (Fig. 4b), suggesting long-term low population sizes in the leatherback turtle.

Within species, overall ROH distributions were generally similar between samples representative of different populations for leatherback turtles, although individuals from the Northwest Atlantic and East Pacific populations displayed slightly higher total aggregate lengths of ROHs than those from the West Pacific population, primarily due to greater aggregate lengths of medium and long ROHs (Fig. 4b). Among green turtles, the aggregate length of ROHs in all categories were generally small and similar across individuals, with the clear exception of the genome reference sample that originated from the Mediterranean population. This individual displayed higher numbers and lengths of long ROHs (>1 Mb) compared to all other green turtles (n = 50 compared to <5, and aggregate length = 74 Mb compared to <4 Mb), suggesting higher levels of recent inbreeding relative to the other green turtle populations represented in our dataset. Comparative analyses mapping this individual to the two previous green turtle assemblies failed to detect these long ROHs (Fig. S23), demonstrating the importance of highly contiguous reference genomes for detecting biologically important patterns using this conservation-relevant metric.

### Genetic load

Coding region variants with predicted high (e.g., stop-codon gain or loss) or moderate impacts were significantly more common in leatherback compared to green turtles (Fig. 4c; high impact variants: t_(4.18)_ = −65.7, p < 0.001; moderate impact variants: t_(4.51)_ = −29.5, p < 0.001). Conversely, low impact and modifier (i.e. variants predicted to cause negligible impacts) variants were significantly more common in green turtles (Fig. 4c; low impact variants: t_(5.88)_ = 4.0, p < 0.01; modifier variants: t_(5.33)_ = 31.8, p < 0.001). The missense to silent mutation ratio was also higher in leatherbacks than green turtles (t_(7.19)_ = – 72.3, p < 0.001; mean = 0.99 and 0.70), further suggesting that genetic load is higher in the leatherback turtles. Within species, there was limited variation between individuals for all variant categories (Fig. 4c).

### Demographic history

Pairwise Sequential Markovian Coalescence (PSMC) analyses indicated different historical effective population sizes (*N_e_*) between the two sea turtle species (Fig. 4d). *N_e_* for all leatherback turtle populations represented in our dataset have been relatively small and sustained over time, ranging in size from approximately 2,000 to 21,000 over the last 10 million years, up until the Last Glacial Maximum (LGM) and at the lower end of this range for most of the last 5 million years. This pattern is consistent between all individuals examined, with similar timings and magnitudes of *N_e_* fluctuations until recent history (Fig. 4d). In contrast, green turtles have experienced wider variation and a higher overall *N_e_* in general, fluctuating between approximately 50,000 and 125,000, until the late Pleistocene, with estimates varying by population (Figs. 4d, S24). While *N_e_* for leatherback turtles is relatively low, it modestly increased prior to the Eemian warm period (Fig. 4d [B]), followed by a subsequent decrease during this period until the LGM (Fig. 4d [A]) when all populations exhibit sharp spikes in *N_e_* possibly due to interocean gene flow following warming after the LGM. In contrast, green turtles generally displayed three distinct peaks in *N_e_* (Fig. 4d), associated with ocean connectivity changes following the closure of the Tethys Sea [D], during the Pleistocene period [C], and prior to the Eemian warming period (Fig. 4d [B]). While the patterns of *N_e_* are broadly similar within green turtles, the timing and magnitude of these fluctuations varied between populations (Fig. S24).

## Discussion

### Divergence in localized RRCs and microchromosomes amidst high global genome synteny

The ancestral lineage leading to leatherback and green turtles diverged over 60 MYA (4), giving rise to species that are adapted to dissimilar habitats, diets, and modes of life. Despite high overall levels of genome synteny between the sea turtle families in both macro- and microchromosomes, RRCs and small microchromosomes were associated with higher concentrations of multicopy gene families, as well as heightened nucleotide diversity and genetic distances between species, suggesting that these genomic elements may be important sources of variation underlying phenotypic differentiation. Higher heterozygosity despite richer gene content in the microchromosomes suggests that these regions are prone to variation accumulation and therefore may have high adaptation value. Though our results here do not demonstrate direct causality, we have identified candidate regions and gene families that can be targeted in further studies quantifying evidence for positive selection and their roles in sea turtle adaptation and speciation.

The high global stability of macro- and microchromosomes between sea turtle families also aligns with recent work showing similar patterns across reptiles, including birds, emphasizing the important roles of microchromosomes in vertebrate evolution (55). Higher evolutionary rates for microchromosomes relative to macrochromosomes has been documented in intraspecific (56) and interspecific (57) studies with chicken and turkey genomes, respectively, so it is possible that the characteristics of microchromosomes and RRCs we observed are not unique to sea turtles, but rather, are prevalent in many vertebrates, which will become clearer as more high-quality assemblies are produced. The mechanisms driving these patterns are not well-understood, but could be related to higher recombination rates in micro-compared to macrochromosomes (58) that result in higher nucleotide diversity and lower haplotype sharing. Once generated, balancing selection may also play a role in maintaining variation in these gene dense regions, but more work is needed across taxa to determine the broad support for these hypotheses. Our detailed analyses of RRCs, microchromosomes, and their associated genes were only possible due to the high-quality of the assembled sea turtle genomes because these analyses can be sensitive to genome fragmentation and mis-assemblies (39). For example, the RRCs and many microchromosomes could not be detected using the draft green turtle genome due to fragmentation and sequence gaps (Figs. S3-4). The prevalence of localized genomic differentiation and underlying mechanisms among other closely or more distantly related vertebrate groups has yet to be widely evaluated due to a lack of equivalent quality genomic resources, but this is rapidly changing. As chromosomal-level genomes across all vertebrate lineages become available, our work provides a roadmap for identifying genomic regions harboring contrasting expansion/contractions of gene families and diversity levels. For taxa with highly conserved genomes like sea turtles, analyses of RRCs and microchromosomes are likely important to understand their divergent evolutionary histories and the phenotypic connections of the genes within them.

### Contrasting sensory and immune gene evolution between sea turtle families

Sea turtles have complex sensory systems and can detect both volatile and water-soluble odorants, which are imperative for migration, reproduction and identification of prey, conspecifics, and predators (59–63). However, leatherback and green turtles occupy dissimilar ecological niches and depend on different sensory cues. While leatherback turtles almost exclusively inhabit the pelagic environment post-hatching, performing large horizontal and vertical migrations to seek out patches of gelatinous prey (64), green turtles recruit to neritic coastal and estuarine habitats as juveniles, and can have highly variable diets (65, 66). Sea turtle nasal cavity morphology also differs between species, with leatherback turtle cavities relatively shorter, wider, and more voluminous than chelonids (67–69), suggesting reduced requirements for olfactory reception. OR genes encode proteins used to detect olfactory chemical cues, with the number of OR genes strongly correlated with the number of detectable odorants (70), and linked to the chemical complexity of the inhabited environment (71). The two major groups of ORs in amniote vertebrates are separated by their affinities with hydrophilic molecules (Class I) or hydrophobic molecules (Class II) (72). Class I OR genes may be particularly important in aquatic adaptation (32), and expansions of Class I ORs in testudines, including green turtles, have been previously reported. However, the accuracy of these estimates for complex gene families using short-read assemblies has been uncertain because they may be prone to mis-assembly (32, 42, 73). We detected an additional 93 Class I OR genes in our green turtle genome compared to those reported in the draft green turtle genome (42), suggesting they can be erroneously collapsed in short-read assemblies. Our reconstruction of both Class I and Class II OR gene evolution throughout the sea turtle lineage revealed that after ancestral contractions, gene copy evolution diverged in opposite directions between the sea turtle families. The greater loss of Class II compared to Class I OR genes in the ancestral sea turtle lineage likely reflects relaxed selection for detection of airborne odorants, as has been observed in other lineages that recolonized marine environments, including marine mammals (74). However, as sea turtles continue to use terrestrial habitats for reproduction, they may need to retain some of these capabilities, which could explain why the observed contraction was weaker than those in exclusively marine species (e.g., the vaquita *Phocaena sinus;* Fig. 2e).

The strong Class I OR expansion in the green turtle may be related to its distribution in complex neritic habitats and variable diet, requiring detection of a high diversity of waterborne odorants, while the continued loss of ORs in the leatherback turtle could be a consequence of its more specialized diet and the lower chemosensory-complexity of pelagic habitats. Although leatherback turtles can detect chemical cues from their prey, sensory experiments have indicated that visual cues are more important for food recognition in this species (75, 76). Additionally, while the precise mechanisms underpinning philopatry in sea turtles remain unclear, green turtles are thought to use olfactory cues to reach specific natal nesting beaches following long-distance navigation guided by magnetoreception (61, 63). In contrast, leatherback turtles exhibit more ‘straying’ from natal rookeries than other species, and such relaxed philopatry may be related to reduced capability to detect olfactory cues to hone in on specific beaches.

Diversity within the highly-complex MHC region is a key component in the vertebrate immune response to pathogens, with greater gene copy numbers and heterozygosity linked to lower disease susceptibility (77). While both sea turtle species contained most of the core MHC-related genes, the green turtle had more copies of genes involved in adaptive and innate immunity. Pathogen prevalence and persistence is often greater in neritic habitats than open ocean habitats (78), so green turtles may be exposed to higher pathogen loads and diversity than leatherback turtles (79). However, reptilian immune systems are understudied compared to other vertebrates, and very few studies of MHC genes have been conducted in turtles (80). Thus, it is not yet understood how immune gene diversity translates into disease susceptibility or ecological adaptation in sea turtles, which is particularly critical for their conservation as FP continues to threaten the recovery of populations around the globe (30). Although this likely viral-mediated tumor disease occurs in all sea turtle species, disease prevalence and recovery greatly varies between and within species, making it plausible that harboring certain genes, copy numbers, or specific alleles may play important roles in disease dynamics. Despite decades of research on this disease (30) only one study on the immunogenomic factors governing FP susceptibility or resilience has been conducted (81), in part due to difficulty in accurately quantifying hypervariable and complex MHC loci with short-read sequencing technologies (82). Our reference genomes now enable studies to accurately interrogate MHC and other immune genes to close this critical research gap and advance our fundamental understanding of immune gene evolution in testudines.

### Differential genomic diversity and demographic histories

Genomic diversity is a critical metric for evaluating extinction risk and adaptive potential to environmental perturbation (83–85), with heterozygosity positively correlated with individual fitness (see reviews by (86, 87). Understanding the causes and consequences of genomic diversity is imperative for sea turtles, and for leatherback turtles in particular, where contemporary populations have experienced recent sharp declines due to human activities (25). Leatherback turtles exhibited exceptionally low genomic diversity relative to the green turtles and other reptiles and mammals, broadly aligning with previous estimates (88, 89). However, factors influencing genomic diversity can vary among species (90), and our PSMC and ROH results indicate that low diversity in the leatherback turtle is likely a consequence of long-term low effective population sizes and historical bottleneck events, rather than a loss during recent population declines. This is consistent with mitochondrial analyses suggesting that contemporary populations radiated from a small number of matriarchal lineages within a single refugium following the Pleistocene (89). The low, yet relatively evenly spread heterozygosity, is also congruent with sustained low population sizes similar to that observed in several mammal species (91, 92). In contrast, the higher heterozygosity, limited ROHs (though see discussion below), and estimated larger, more variable historical *N_e_* in green turtles likely reflects their radiation from many refugia and frequent admixing of populations (93).

Regardless of the causes of current genomic diversity levels in sea turtles, the amount of standing variation can have important implications for species’ future persistence (94), especially given the adaptive capacity likely required to keep pace with rapid anthropogenic global change. Although informative genome-wide diversity estimates can be made without high-quality reference genomes, these enable deeper examination of diversity patterns that are relevant for conservation. For example, the use of our reference genomes demonstrated that diversity is very low within coding regions of the leatherback turtle genomes, indicating limited standing functional variation that may have implications for the adaptive potential of this species to novel conditions. Additionally, leatherback turtles exhibited higher genetic load compared to green turtles, and this signal was consistent across all samples regardless of population source. Leatherback turtles have substantially lower hatching success compared to other sea turtle species (29), potentially related to the heightened genetic load and low heterozygosity (95, 96), and may combine with other factors to slow population recoveries despite conservation measures. However, recent studies have documented low genome diversity in a number of species with wide geographic distributions and relatively large census population sizes, including some long-lived marine vertebrates (91, 97–100). Additionally, other species with low diversity have rebounded following population declines and/or appear to have purged deleterious alleles through long-term low population sizes (98, 101, 102), thereby limiting the impacts on viability (54, 98, 103). Although our results of greater genetic load despite long-term low *N_e_* suggest this is not the scenario for leatherback turtles, further assessments including more individuals over greater spatial and temporal scales are needed. Studies enabled by the reference genomes presented here quantifying diversity and genetic load within and among global populations will clarify these relationships for leatherback and other sea turtle species to guide conservation recommendations.

While diversity and genetic load patterns were consistent within species, ROH analyses revealed variation providing possible insight into different recent population histories, though these results must be interpreted with caution given our limited sample sizes. Although all leatherback turtles displayed a high number and aggregate length of short ROHs, consistent with a historical bottleneck generating long ROHs that are subsequently broken by mutations and recombination events (104), individuals from the West Pacific population show limited evidence of recent inbreeding (i.e., few long ROHs). In contrast, individuals from the Northwest Atlantic and Eastern Pacific harbor higher aggregate lengths of medium and long ROHs, suggesting more recent breeding of related individuals. These populations differ substantially in their recent census sizes and trends (105) that are generally concordant with these patterns; for example, the Western Pacific meta-population is relatively larger but declining, while some Northwest Atlantic populations have undergone rapid increases and others remain small and isolated. However, there is limited knowledge of abundances for these populations prior to the last several decades, and the long generation times of sea turtles makes it likely that impacts of very recent demographic changes may not yet be fully reflected in the genomes. Thus, as conservation efforts continue to mitigate the ongoing major anthropogenic threats to the survival of this species, genomic monitoring over longer temporal scales is needed to discern if populations are likely to encounter complications arising through inbreeding depression during recovery. In green turtles, long ROHs were absent or in very low numbers in all individuals, with the striking exception of the reference individual from the Mediterranean. This isolated population that has undergone severe decline over the last century due to human exploitation (106), and our results indicate that consequent inbreeding is likely occurring, which may impact its recovery. The specific individual was from the Israel green turtle rookery that is estimated to have only 10-20 nesting females in the last decade (107, 108). However, it is currently unclear if Israel is demographically isolated from other rookeries in the region (108, 109), so further research is needed to understand if inbreeding is a concern only for this nesting aggregation, or the Mediterranean population more broadly. Finally, these findings highlight the utility of ROH even in animals with long generation times, and the importance of using highly contiguous genomes for accurate ROH assessment to inform conservation.

The lower, long-term *N_e_* of leatherback turtles detected in our demographic reconstructions may be associated with this species’ greater mass and trophic position, as was found in recent study assessing relationships between key life-history traits and genomic variation in avian species (110). While it is widely documented that environmental changes can strongly impact species’ abundances and distributions (111–113), following an initial decrease associated with declining temperatures, *N_e_* of leatherback turtles remained relatively constant throughout the substantial temperature fluctuations of the Pleistocene. As ectotherms, reptiles are generally sensitive to climatic thermal fluctuations, however, leatherback turtles exhibit unique physiological adaptations that produce regional endothermy and facilitate exploitation of cold-water habitats (6) that potentially led them to being less susceptible to periods of cooler temperatures. In contrast, wide fluctuations for green turtles appear correlated with climatic events, beginning with the closure of the Tethys Sea, which altered ocean connectivity and represented a period of increasing temperatures that may have opened more suitable habitat. As temperatures subsequently decreased, *N_e_* also decreased, however temperature fluctuations during the Pleistocene were associated with an additional increases in *N_e_*. While warmer temperatures presumably allowed for larger population sizes of green turtles, large spikes in *N_e_* around the Eemian warming, particularly for the Mediterranean individual, are very likely associated with mixing of previously isolated populations due to warm-water corridors allowing movement between populations and ocean basins (114). While our overall estimates and trends for both species were broadly concordant with previous studies (89, 115, 116), a recent study using Multiple Sequentially Markovian Coalescent (MSMC2) analyses found steep declines in *N_e_* for green turtles >100,000 years before present (116), which was not detected in our PSMC analyses. Since this decline was also not detected in a prior study using PSMC on the draft green turtle genome (115), and demographic inferences are generally robust to genome quality (117, 118), this is likely a consequence of the different methods, with MSMC analyses inferring large *N_e_* for more ancient time scales (117).

### Enabling future research and conservation applications

In addition to the insights reported here, the reference genomes for both extant sea turtle families provide invaluable resources to enable a wide breadth of previously unattainable fundamental and applied research. Combined with other forthcoming chromosomal-level vertebrate genomes (39), in-depth comparative genomics analyses can further investigate ecological adaptation related to immune and sensory gene evolution, as well as the genomic basis for traits of interest such as adaptation to saltwater, diving capacity, and long-distance natal homing. Studies leveraging these reference genomes alongside whole-genome sequencing of archival sample collections can assess how genomic erosion, inbreeding and mutational load are linked to population size, trajectories, and conservation measures in global sea turtle populations. For instance, the fact that leatherback turtles have persisted with low diversity and *N_e_* for extended periods offers hope for their recovery, but given that some populations have now been reduced to only a few hundred individuals (105), research quantifying purging of deleterious alleles, inbreeding depression, and adaptive capacity within populations is urgently needed (119). We emphasize that high-quality reference genomes are not required for all research goals, and combined with other recent studies (117, 118, 120), our findings provide clear guidance on when they may, or may not, be necessary in order to generate accurate results to inform conservation. For example, genome-wide diversity estimates are typically robust to assembly quality, but the ability to detect long ROHs can be strongly affected. As ROH metrics are increasingly being used to guide species management plans (121–123), it is important for researchers to understand how genome quality may impact their analyses and inferences. Additionally, many conservation applications that may not explicitly require whole-genome data can also directly benefit from the utility of these reference genomes, including the development of amplicon panels and molecular assays to investigate TSD mechanisms and adaptive capacity under climate change, and assessing linkages between immune genes and disease risk. Finally, with global distributions and long-distance migratory connectivity, sea turtle conservation requires international collaboration that has been previously hampered by difficulty comparing datasets between laboratories. Existing anonymous markers (e.g. microsatellites and restriction-site based SNP markers) can now be anchored to these genomes, and new ones can be optimized for conservation-focused questions and shared across the global research community, facilitating large-scale syntheses and equitable capacity building for genomics research. While ongoing anthropogenic impacts continue to threaten the viability of sea turtles to persist, combined with the important work of reducing major threats such as fisheries bycatch and habitat loss, these genomes will enable research that make critical contributions to recovering imperiled populations.

## Methods

### Reference sample collections, genome assembly and annotation

Blood was collected from leatherback and green turtles using minimally invasive techniques for isolation of ultra-high molecular weight DNA, and tissue samples of internal organs for RNA were collected opportunistically from recently deceased or euthanized animals. Full details of sample collection, storage, and laboratory processing prior to sequencing can be found in Supplementary Appendix I. Resulting raw data were deposited into the VGP Genome Ark and NCBI Short-Read Archive (SRA) (see Data Accessibility Statement). We assembled both genomes using four genomic technologies following the VGP pipeline v1.6 (39) with a few modifications detailed in Supplementary Appendix I. Briefly, PacBio Continuous Long Reads were assembled into haplotype phased contigs, with contigs scaffolded into chromosome-level super scaffolds using a combination of 10X Genomics linked reads, Bionano Genomics optical maps, and Arima Genomics Hi-C 3D chromosomal interaction linked reads. Base call errors were corrected to achieve high quality (>Q40). The assemblies were manually curated, with structural errors corrected according to the Hi-C maps (Fig. S1), and the 28 super scaffolds (hereinafter referred to as chromosomes) numbered in both species according to sequence lengths in the leatherback turtle assembly, and synteny between the two species. A manual inspection comparing the sequence collinearity between the first curated versions of the genomes revealed a small number of artefactual sequence rearrangements that were corrected in a second round of manual curation (see Supplementary Appendix I).

To enable accurate, species-specific annotations for each genome, both short and long-read transcriptome data (RNA-Seq and Iso-Seq) were generated from tissues known for their high transcript diversity in each species. These data, plus homology-based mapping from other species, were used to annotate the genomes using the standardized NCBI pipeline (124). We performed annotation as previously described (39, 125), using the same RNA-Seq, Iso-Seq, and protein input evidence for the prediction of genes in the leatherback and green turtles. We aligned 3.5 billion RNA-Seq reads from eight green turtle tissues (blood, brain, gonads, heart, kidney, lung, spleen and thymus) and 427 million reads from four leatherback turtle tissues (blood, brain, lung and ovary) to both genomes, in addition to 144,000 leatherback turtle and 1.9 million green turtle PacBio IsoSeq reads, and all Sauropsida and *Xenopus* GenBank proteins, known RefSeq Sauropsida, *Xenopus,* and human RefSeq proteins, and RefSeq model proteins for *Gopherus evgoodei* and *Mauremys reevesii*.

### Genome quality analysis

We used the pipeline assembly-stats from https://github.com/sanger-pathogens/assembly-stats to estimate scaffold N50, size distributions and assembly size. BUSCO analysis (115) and QV value estimations (116) were conducted to assess the overall completion, duplication, and relative quality of the assemblies. We used D-GENIES (118) with default parameters to conduct dot plot mapping of the entire genomes and each individual chromosome to evaluate the synteny between leatherback and green turtle genomes, and Haibao Tang JCVI utility libraries following the MCScan pipeline (119) to verify the contiguity of the genomes. Incongruences in gene synteny blocks were manually investigated using Artemis Comparative Tool (120), identifying possible regions of inversion that could be caused by artifacts during assembly. These regions were then identified and corrected in the latest version of the assembly for both species. Only a few structural rearrangements between the two species remained after two rounds of manual curation with support of sequencing data. The final curated assemblies were analyzed using the Genome Evaluation Pipeline (https://git.imp.fu-berlin.de/cmazzoni/GEP) to obtain all final QC plots and summary statistics.

### Identification and analysis of RRCs and REs

Leatherback and green turtle genomes were mapped to each other using Minimap2 with a dot plot of the mapping generated using D-GENIES (126). Using windows of 20 Mb, the dot plot was screened visually with regions larger than 1 Mb showing reduced collinearity (i.e., one or more breaks in the diagonal indicating homology), as well as smaller regions with obvious signals of genomic rearrangements (e.g., inversions), cataloged as regions of reduced collinearity (RRCs; Fig S5). Several genomic features (e.g. GC content, repeat elements) were examined within these regions and compared to regions of the same length directly up- and down-stream of the RRCs, which are most likely under relatively similar cellular and molecular influences to the RRCs (Table S9). We identified the functions of the genes present in RRCs using genome annotations and identified protein domains using Interproscan (127). The proportion of GO terms in each chromosome was estimated for each species using PANTHER (128); Fig. S25). To examine if RRCs presented differential patterns of sequence and/or gene duplication between the species, we aligned the genomes of the sea turtles against each other using Progressive Cactus (129, 130), and all homologous genes that presented more than one copy for one of the two species were isolated using an inhouse script (*IdentifyDupsReciprocalBlast.sh*) to retrieve duplicated genes (see Supplementary Appendix I for further details on Cactus alignments). Repetitive elements (REs) were identified by creating a *de novo* database of transposable elements using RepeatModeller2 (131), followed by running RepeatMasker (132, 133) to calculate Kimura values for all REs (see full analysis details in Supplementary Appendix I).

### Gene families and gene functional analysis

To estimate the timing of gene family evolution for the OR gene families on sea turtles we used Computational Analysis of gene Family Evolution v5 (134). Briefly, CAFE uses phylogenomics and gene family sizes to identify gene family expansions and contractions. We used a dataset containing 8 species of turtle, 4 non-turtle reptiles, 3 mammals and 1 amphibian using OrthoFinder (135, 136). OR orthogroups were grouped based on subfamily (Class I and Class II; see (73)), and an ultrametric phylogeny was generated by gathering 1:1 orthologs. We then aligned amino acid sequences for each orthogroup and generated a phylogenetic tree (see Supplementary Appendix I for details).

To identify genes related to immunity, and the MHC in particular, we searched the genome for the list of core MHC genes following Gemmell et al. (2020) (46). We conducted initial searches of gene identifications, followed by a search of protein identifiers. As genes associated with the MHC are diverse, and vary substantially among species, we did not use a BLAST search for these genes. Locations of the genes were then compared between species to determine which genes were annotated, and where the core MHC region is located within the genomes. We conducted a search following similar methods for genes with known function in TSD in other reptiles (see Supplementary Appendix I for details).

### Genetic distance, genome diversity, and runs of homozygosity

To estimate the genetic distance between the leatherback and green turtle genomes, we used the halSnps pipeline (137) to compute interspecific single variants based on genome alignments obtained with Progressive Cactus (129, 130) using the leatherback turtle genome as the reference. Genetic distances were calculated for windows across the genome where each window included exactly 10,000 positions presenting single alignments against the green turtle genome in the Cactus output. Positions with zero, or more than one alignment were ignored, and if this occurred over more than 50% of a given window, it was skipped entirely (i.e., each window analyzed covered between 10 and 20 Kb of the genome). Interspecific distances per bp were calculated by dividing the number of variants found within a window by 10,000. Differences in genetic distance, gene content, GC content, and heterozygosity between macro-, intermediate micro-, and small microchromosomes were tested using one-way ANOVAs for each species. Regression analyses were used to test for correlations between these measures across chromosomes.

For genome diversity, ROH, demographic history, and genetic load analyses, we also included whole-genome resequencing (WGR) data for additional individuals representing multiple global populations in each species (Table S13 and Supplemental Appendix I Methods for sample details). We calculated genome-wide heterozygosity using a method adapted from Robinson et al. (2019) (92), which used the Genome Analysis Toolkit (GATK) (138) to call genotypes at every site across the genome from reads mapped to our reference genomes with BWA-mem (139). To avoid any biases arising from differences in processing between samples, 10X linked-reads from the reference individuals were initially processed using the proc10xG pipeline (https://github.com/ucdavis-bioinformatics/proc10xG), and then treated identically to Illumina short-read data from resequenced individuals. Heterozygosity was calculated within 100 Kb non-overlapping windows, with only sites that had a depth of between ⅓× and 2× mean coverage retained for genotype scoring. Heterozygosity was calculated for (1) the entire genome, (2) the genome with repeat-regions masked, (3) exonic regions, (4) and for the non-coding regions. Statistical comparisons between species were made using T-tests, with paired T-tests used when comparing between regions within species. We subsequently applied the heterozygosity pipeline to generate genome-wide heterozygosity for a number of additional reptilian species with sequences sourced from the NCBI SRA where species-specific reference genomes were available (see details in Supplementary Appendix I).

ROHs were identified by initially generating a SNP-list using the Analysis of Next Generation Sequencing Data (ANGSD; (140) pipeline. ANGSD was parameterized to output files that were configured for use as input for the ROH analysis incorporated in PLINK (141). ROHs were then further characterized as ‘short’ (50-500 Kb), ‘medium’ (500Kb-1 Mb), or ‘long’ (>1 Mb), with size class categories loosely based on (104).

### Genetic load

Estimates of deleterious allele accumulation were conducted using the snpEff variant annotation software (142). We estimated the impacts of variants (SNPs and INDELs) from coding regions using the species-specific genome annotations generated for both species, with a total of 18,775 genes for the leatherback turtle genome, and 19,752 genes for the green turtle genome used in the analysis. gVCFs were generated for each individual followed by joint-genotyping using GATK (138), allowing the reference individuals to include homozygous alleles found in other individuals. Combined VCFs were then separated for each individual and filtered using based on depth of coverage (⅓× – 2× mean coverage for each individual). The snpEff program predicts variant impacts and bins them into ‘high’, ‘moderate’, or ‘low’ impact categories, and outputs a list of genes that have predicted variant effects. We ran the snpEff analysis on all individuals for both species, and compared the percentages of each variant between species using T-tests.

### Historical demography

Pairwise Sequential Markovian Coalescence (PSMC; (143)) analyses of demographic history were employed for all individuals for both species. We used SAMtools (144) and BCFtools (145) to call genotypes with base and mapping quality filters of >Q30, before filtering for insert size (50-5,000bp) and allele balance (AB) and retaining only biallelic sites with an AB of <0.25 and >0.75. We then ran PSMC analysis using the first 10 scaffolds, which constituted over 84% of the total length of the genome. We scaled our outputs using a generation time of 30 years (mid-way between reported generation times for both species; see Supplementary Appendix I), and a mutation rate of 1.2 × 10^-8^ (115).

## Abbreviations

TE: transposable element
RE: repetitive element
RRC: region of reduced collinearity
FP: Fibropapillomatosis
ROH: runs of homozygosity

## Acknowledgments

We are grateful for the assistance with the (1) leatherback turtle sample collection from the St. Croix Sea Turtle Program and the US Fish and Wildlife Service, the NOAA-SWFSC California in-water leatherback research team, and the New England Aquarium; (2) green turtle sample collection from the Israel National Sea Turtle Rescue Centre, the NOAA PIFSC-MTBAP team, and Thierry Work (USGS). We also thank Estefany Argueta and Jamie Adkins Stoll for assistance with literature searches for TSD and immune genes and library preparations for leatherback resequenced individuals, and Phillip Morin, Andrew Foote, Anna Brüniche-Olsen, Annabel Beichman, Morgan McCarthy, David L. Adelson, and Yuanyuan Cheng for their invaluable discussions surrounding analysis approaches and comments on the manuscript. Sequencing of the green turtle has been performed by the Long Read Team of the DRESDEN-concept Genome Center, DFG NGS Competence Center, part of the Center for Molecular and Cellular Bioengineering (CMCB), Technische Universität Dresden and MPI-CBG. For green turtle resequenced samples, we also thank Jessica Farrell, Whitney Crowder, Brooke Burkhalter, Nancy Condron, and the veterinary and rehabilitation staff and volunteers of the University of Florida’s (UF) Sea Turtle Hospital at Whitney Laboratories, and the staff of the South Carolina Aquarium for sampling support. Thanks also are due to UF’s Interdisciplinary Center for Biotechnology Research Core for sequencing services, and to the Florida Fish and Wildlife Conservation Commission and South Carolina Department of Natural Resources for valuable assistance with permitting. Leatherback DNA for additional WGR came from samples archived in NOAA-SWFSC Marine Mammal and Sea Turtle Research Collection that were originally collected as part of previous population genetics studies, and we thank Erin LaCasella for her help processing samples from the SWFSC.

## Data Accessibility Statement

Assemblies for both species have been deposited on NCBI GenBank. The NCBI GenBank accession numbers for the leatherback turtle genome assembly (rDerCor1) are GCF_009764565.3 and GCA_009762595.2 for the annotated primary and original alternate haplotypes in BioProject PRJNA561993, and for the green turtle assembly (rCheMyd1) are GCF_015237465.2 and GCA_015220195.2 for primary and alternate haplotypes respectively in BioProject PRJNA561941. The raw data used for assemblies are available on the Vertebrate Genome Ark (https://vgp.github.io/genomeark/). The leatherback turtle RNA-Seq data generated for the purpose of assembly annotation was deposited in the SRA under accession numbers SRX8787564-SRX8787566 (RNA-Seq) and SRX6360706-SRX6360708 (ISO-Seq). Green turtle RNA-Seq data generated for annotation were deposited in SRA under accessions SRX10863130-SRX10863133 (RNA-Seq) and as SRX11164043-SRX11164046 (ISO-Seq). The NovaSeq 6000 DNA-Seq data for the green turtle resequencing, including raw reads, are deposited in NCBI (https://www.ncbi.nlm.nih.gov/) under BioProject ID: PRJNA449022. All scripts used for downstream analyses following genome assembly and annotation have been deposited on GitHub under repository https://github.com/bpbentley/sea_turtle_genomes.

