## Supplementary Appendix for "Divergent sensory and immune gene evolution in sea turtles with contrasting demographic and life histories"

**Supplemental Information Appendix**

### I. Extended Materials & Methods

All scripts associated with these analyses have been deposited under GitHub repository https://github.com/bpbentley/sea_turtle_genomes.

*Sample collection & data generation*

The conservation status of leatherback (*Dermochelys coriacea*) and green (*Chelonia mydas*) turtles precludes the sacrifice of individuals to obtain tissue samples, so for reference genome assembly, blood was collected using minimally invasive techniques for isolation of ultra-high molecular weight DNA from a male leatherback turtle off the coast of Monterey, California (NMFS ESA10a1A permit #21260 and USFWS Recovery Permit #TE-72088A-3) and a captive male green turtle in Israel National Sea Turtle Rescue Centre (INPA Permit worker 02457/2021 given to YL). While the green turtle used for reference genome assembly was housed in captivity, this individual was collected from the wild for the purposes of population augmentation in the year 2000, and genetically represents the wild Mediterranean population. Blood samples were flash frozen following collection and stored at -80°C until processing. Frozen subsamples of whole blood were placed in 1ml of 95-100% ethanol and processed using a modified version of the Bionano blood DNA isolation protocol optimized for frozen whole nucleated blood stored in ethanol (https://bionanogenomics.com/wp-content/uploads/2017/03/30215-Bionano-Prep-Frozen-Blood-Protocol.pdf). DNA quality was assessed using pulse field gel electrophoresis (PFGE) (Pippin Pulse, SAGE Science, Beverly, MA) or the Femto Pulse instrument (Agilent Technologies, Santa Clara, CA). DNA was then further prepared for the different library types (PacBio, 10X Chromium and Bionano optical map imaging) as described in Rhie et al. (1). Hi-C of the green turtle was performed on flash-frozen blood following the Arima Hi-C protocol (Arima Hi-C user guide for Animal tissues, v01, Material Part Number: A510008).

Tissue samples of internal organs for RNA were collected opportunistically from recently deceased or euthanized animals in the US Virgin Islands, New England Aquarium, and the National Marine Fisheries Service Pacific Island Fisheries Science Center (NMFS permit #15685), flash frozen and stored at -80°C until processing. Total RNA was extracted placing 20-30mg of frozen tissue on dry ice and cut into 2mm pieces before being disrupted and homogenized with the Qiagen TissueRuptor II (Cat No./ID: 9002755), followed by extraction using Qiagen kits (leatherback turtle: gonad, lung and brain tissues using QIAGEN RNeasy kit, Cat. No. 74104; green turtle: brain, gonads, thymus, and spleen using QIAGEN RNeasy Protect kit, Cat. No. 74124). The quality and quantity of RNA were measured with a Qubit 3 Fluorometer (Qubit RNA BR Assay Kit, Cat no. Q33216; ThermoFisher Scientific, Waltham, MA) and a Fragment Analyzer (Agilent Technologies); RINs were within 7.5-9.5. Libraries were then prepared for short-read Illumina sequencing (RNA-Seq) and long-read PacBio sequencing (Iso-Seq). For RNA-Seq, aliquots of total RNA from each tissue and both species were sent to Psomagen (Rockville, MD) for library preparation (TruSeq stranded mRNA kits, Illumina) and sequencing. For the leatherback turtle, PacBio Iso-seq libraries were prepared according to the 'Procedure & Checklist - Iso-SeqTM Template Preparation for Sequel® Systems' (PN 101-070-200 version 05) without Blue Pippin size selection. Briefly, cDNA was reversely transcribed using the SMRTer PCR cDNA synthesis kit from 1 µg total RNA and amplified in a large-scale PCR. Two fractions of amplified cDNA were isolated using either 1x AMPure beads or 0,4x AMPure beads. Both fractions were pooled equimolar and went into the Pacbio SMRTbell template preparation v1.0 protocol following the manufacturer's instruction. For the green turtle, PacBio Iso-seq libraries were prepared according to the ‘Procedure & Checklist – Iso-Seq™

Express Template Preparation for Sequel® and Sequel II Systems' (PN 101-763-800 Version 01). Briefly, cDNA was reverse transcribed using the NEBNext® Single Cell/Low Input cDNA Synthesis & Amplification Module (New England BioLabs, cat. no. E6421S) and Iso-Seq Express Oligo Kit (PacBio PN 10 1-737-500) from 300ng total RNA. Forward and reverse barcoded primers were used during cDNA amplification. PacBio Iso-seq libraries were sequenced on one PacBio 8M SMRT Cell (PN: 101-389-001) on the Sequel II instrument with Sequencing Kit 2.0 (PN: 101-820-200) and Binding Kit 2.1 (PN: 101-843-000) and 24 hours movie with 2 hours pre-extension. Resulting raw data was deposited into the NCBI Short-Read Archive (SRA) for genome annotation (see Data Accessibility Statement).

For green turtle whole genome resequenced individuals, sequence data was obtained from rehabilitating animals in Florida (University of Florida's Sea Turtle Hospital at the Whitney Laboratory for Marine Bioscience) and South Carolina (South Carolina Aquarium) as previously described (2). Briefly, green turtle tissue (non-tumor skin, lung and kidney) was obtained from juvenile rehabilitation patients by 4 mm punch biopsies during the tumor removal surgery, or during necropsies conducted immediately after euthanasia. Whole blood was obtained during routine blood draws during veterinary examinations. DNA was extracted using a DNeasy Blood & Tissue Kit (Qiagen, Cat No. 69504). Sequencing of these samples was conducted at the University of Florida's Interdisciplinary Center for Biotechnology Research Core Facilities, using an Illumina NovaSeq 6000, paired-end reads (2x150bp) at a depth of approximately 80x *C. mydas* genome coverage. Sequencing libraries were constructed using the NEBNext UltraTMII DNA Library Prep Kit for Illumina (Cat# E7645S), PCR was done for only 6-7 cycles of amplification in order to minimize duplicate reads. Barcoding was done using the indexing reagents provided in the NEBNext Unique Dual Index Oligos kit (Cat# E6440S). Green turtle sampling was carried out under permit numbers MTP-17-236 and MTP-2019-0005 from the Florida Fish and Wildlife Conservation Commission and the South Carolina Department of Natural Resources, respectively, with ethical approval from the University of Florida's Institutional Animal Care and Use Committee (IACUC). We additionally downloaded Illumina short-reads of the West Pacific green turtle that was used for the assembly of the draft genome (3) from the SRA (SAMN02981410). For leatherback whole genome resequenced individuals, we used DNA previously extracted from samples in the NOAA Marine Mammal and Sea Turtle Research Tissue Collection (La Jolla, CA). DNA was assessed for quality and quantity with a Fragment Analyzer (model number: 5200, Agilent Technologies, Santa Clara, CA) using a high-sensitivity sensitivity large fragment kit (Agilent Technologies, Santa Clara, CA, DNF-462). We prepared individually barcoded whole genome libraries following the protocol of (4), with minor modifications to use $1/10^{th}$ reactions. Libraries were assessed on the Fragment Analyzer as previously described, pooled in equal molarity and sequenced on an Illumina NovaSeq System at Novogene Corporation (Sacramento, CA). Note that these samples were run with additional samples for a companionate project, so their total coverage is not equivalent to that of a full NovaSeq lane.

*Genome assembly & curation*

Both genomes were assembled following the VGP pipeline v1.6 (1) with a few modifications. Initially, all genomic data from each species were screened for low quality and contamination with Mash (5) as described by Rhie et al. (2021). A preliminary analysis was performed using the 10X Illumina data (with 24bp-barcodes trimmed-off) and GenomeScope 2.0 (6) to estimate the haploid genome length, repeat content, and heterozygosity and with a k-mer size of 21bp (Fig. S1). The predicted genome length was used to help select the amount of PacBio reads covering 50× of the genome. The selected PacBio reads were first corrected and subsequently assembled into partially phased contigs using FALCON and

FALCON-unzip (7). The primary assembly was further purged of false haplotype duplications using purge_dups (8) and all removed regions were assumed to represent haplotype retention and added to the alternative assembly (Fig. S1). Scaffolding of the primary assembly was performed in three major steps. First, the 10XG linked reads were aligned to the primary contigs, and two scaffolding rounds were performed using scaff10x v2.2 (https://github.com/wtsi-hpag/Scaff10X). Subsequently, Bionano cmaps were generated using the Bionano Pipeline in non-haplotype assembly mode and used to further scaffold the assembly with Bionano Solve v3.2.1. We used the DLE-1 one enzyme non-nicking approach, and scaffold gaps were sized according to the software estimate. Finally, Hi-C reads were aligned to the Bionano cmaps scaffolded assembly using the Arima Genomics mapping pipeline (https://github.com/ArimaGenomics/mapping_pipeline), as described in Rhie et al. (2021). The restriction enzymes used to generate each library were specified using parameters *-e GATC*, *GANTC* for Arima reads. The processed Hi-C alignments were then used for scaffolding with Salsa2 (9) using the parameters *-m yes -i 5 -p yes*. In parallel, the mitochondrial genome was assembled by the mitoVGP pipeline (10) using the corrected PacBio reads and 10XG reads as input.

Following the scaffolding steps, primary, alternative and mitochondrial assemblies were concatenated for two rounds of nucleotide polishing. As described in Rhie et al. (2021), a first round of polishing was performed with Arrow (11) using the PacBio CLR reads, followed by two rounds of polishing using the 10XG Illumina short-reads. For the latter, reads were first aligned to the assembly with Longranger align 2.2.2 (12) and variants were called with FreeBayes v1.2.0 (12) using default options. Consensus were called with bcftools consensus (13). To minimize the impact of the remaining algorithmic shortcomings, both assemblies were subjected to multiple rounds of rigorous manual curation (14). All data generated for both of the resulting assemblies; rDerCor1 and rCheMyd1 were collated, aligned to the primary assembly and analyzed in gEVAL (15); (https://vgp-geval.sanger.ac.uk/index.html), visualizing discordances in a feature browser and issue lists. In parallel, each species' Hi-C data were mapped to the primary assembly and visualized using Juicebox (16, 17) and HiGlass (18). Based on identified mis-joins, missed joins and other anomalies from genome curators, the primary assembly was corrected accordingly. A second round of curation was performed after the synteny analysis between both genomes revealed a small number of remaining anomalies. Table S2 shows genome assembly statistics generated using gfastats (19) for both genomes before and after curation.

*Genome annotation*

Annotation was performed as previously described (1, 20), using the same RNA-Seq, IsoSeq and proteins input evidence for the prediction of genes in the leatherback and green turtle. A total of 3.5 billion RNA-Seq reads from eight green turtle tissues (blood, brain, gonads, heart, kidney, lung, spleen and thymus) and 427 million reads from four leatherback turtle tissues (blood, brain, lung and ovary) were aligned to both genomes, in addition to 144,000 leatherback and 1.9 million green turtle PacBio IsoSeq reads, all *Sauropsida* and *Xenopus* GenBank proteins, all known RefSeq *Sauropsida, Xenopus*, and human RefSeq proteins, and RefSeq model proteins for *Gopherus evgoodei* and *Mauremys reevesii*. Prediction of the function of gene models was done by calculating their orthologs to human proteins and annotated on reference GRCh38.p12 using a combination of protein sequence similarity and local synteny information. Turtle genes for which a human ortholog could be determined inherited the symbol (e.g. BRCA1) and description from the human gene. The remaining genes were assigned LOCXX identifiers. Their putative function was obtained from their best BLASTP hit to the UniProtKB/SwissProt database.

*Transposable element analysis*

Transposable elements (TEs) from the genomes of the leatherback and green turtles were identified by creating a denovo database of transposable elements using RepeatModeller2 (21) using the module -LTRStruct for each genome. Using this database, RepeatMasker (22, 23) was run with the additional parameters of -a -s -gccalc to calculate kimura values for all the transposable elements identified using the script *calcDivergenceFromAlign.pl* with the parameters -s and -a. An inhouse script was also used, *align_with_divHandeler.py*, to isolate the TEs flagged as Unknowns from which each representative sequence of all TE families of Unknowns was isolated. Once isolated, the distribution of size and number of transposable elements was analysed for both genomes for the complete scaffolds and for the low synteny regions using the inhouse script StatsTeRegion.py (Table S8); CheckNesting.py, Size_nesting.py (Table S8); Calculate_masking_size.sh and *createRepeatLandscape.pl* with the same parameters used in the first iteration, to create the TE landscape presented in Fig. S8.

*Genome alignment*

The genomes of the sea turtles were aligned against each other using two outgroups. For this, genome assemblies of four turtle species (leatherback turtle, green turtle, *Gopherus evgoodei* [GCA_007399415.1] and *Mauremys reevesii* [GCA_016161935.1]) were first soft-masked with RepeatMasker to reduce the total number of potential genomic anchors formed by the many matches that occur among regions of repetitive DNA. Progressive Cactus, a reference-free whole genome aligner, was used (24, 25) to align all other genomes applying the parameter --realTimeLogging. The guide tree and divergence time used as input for Cactus were retrieved from (26), with branch lengths reflecting neutral substitutions per site. To obtain an alignment only for the two sea turtles the parameter --root was used, setting as root the ancestral of the two sea turtles. For the alignment among all four turtles no root was set.

*Analysis of regions of low synteny*

Leatherback and green turtle genomes were mapped to each other using Minimap2 and a dot plot with the mappings was generated using D-GENIES (27) to evaluate genome synteny and identify regions that presented low identity or structural rearrangements. Specifically, windows of 20 Mb were screened by eye in the dotplot, and every region bigger than 1 Mb presenting one or more breaks in the synteny was cataloged (Table S6; Fig. S5). Some regions smaller than 1 Mb but larger than 100,000bp that contained obvious signals of genomic rearrangements were also cataloged for future analysis. To identify if these low syntenic regions present differences in content or nucleotide composition, they were compared to two sections of the same length immediately upstream and downstream in the chromosome. In cases where the low syntenic region was located at one of the chromosome extremities, either two upstream or downstream sections were used for comparison in order to maintain the total number of sites used for comparison (Table S9). The function of the genes present on those regions were extracted using the annotation results as well as the identification of protein domains using Interproscan (28). To verify if the low synteny regions present a pattern of higher sequence duplication, the Cactus alignment was analyzed. First, the tool hal2maf from HalTools (29) was used to convert the output of cactus to the .maf format selecting (1) green turtle as reference and (2) leatherback turtle as reference. Also, using the coordinates for the low synteny regions, coding sequences (CDS) were isolated from the genomes fasta files based on the coordinates provided by the annotation file (.gff) using GFFreads tool (30). A reciprocal blast (31) was performed between the two species and, for each low synteny region, all homologous genes that presented more than one copy for one of the two species were isolated to retrieve duplicated genes using an inhouse script.

To determine if olfactory receptor (OR) genes were more numerous in one of the species throughout the genome in addition to the differences found within RRCs, we searched the annotation for the term “olfactory”. Grep searches were performed on annotation files (gff) for both sea turtle species, *M. reevesii, G. evgoodei* and *T. scripta* in order to identify and compare gene numbers between these species. ORs were considered as Class I if numbered 51-56, while the remaining ORs were considered as Class II genes. After preliminary findings showing consistent higher gene copy numbers in the green turtle, we performed multiple analyses in order to rule out the possibility of collapsed multicopy genes in the leatherback turtle assembly. Specifically, we checked gene connections based on similarity for each set of gene copies manually, and estimated the predicted number of multicopy genes based on short read (Illumina 10X data) coverage for each RRC (Table S7). Neither analysis showed evidence of gene collapse in the leatherback turtle, indicating that observations were biological rather than technical artifacts.

*Gene families and gene functional analysis*

To estimate the timing of gene family evolution for the olfactory receptor gene families on sea turtles we used Computational Analysis of gene Family Evolution v5 (32) https://github.com/hahnlab/CAFE5). CAFE5 uses phylogenomics and gene family sizes to identify gene families with rapid expansions and/or contractions for all branches in a phylogeny. First, we generated a dataset containing the numbers of OR genes for a dataset containing 8 species of turtles, 4 non-turtle reptiles, 3 mammals and 1 anura species using Orthofinder v 2.5.4 (33, 34). OR orthogroups were grouped based on OR class I and class II subfamilies as described previously (35) and identified from the human genome (36). We generated an ultrametric phylogeny by gathering all 1:1 orthologues identified by Orthofinder. We aligned amino acid sequences from each ortholog group with MAFFT v6.864b (37) using default parameters and trimmed with Trimal v1.4 (38) using the “automated1” algorithm. Then we concatenated the trimmed alignments in a supermatrix using geneSticher.py (https://github.com/ballesterus/Utensils/blob/master/geneStitcher.py) and generated a tree with IqTree v2.1.4 (39, 40), considering each orthogroup as a partition and with 1000 bootstrap. We then calibrated the tree using r8s (41) with the same known evolutionary divergences based on fossil records used by (3).

We additionally searched the genomes for known TSD-related genes. We initially searched the annotation files (gff) using gene identification strings and protein names from our compiled reference list of 217 genes (see Table S11 for details) using a ‘grep’ search. Given that some genes have many aliases depending on the lineages they were discovered in, and their function, we additionally applied a BLAST (42) search using orthologous protein sequences pulled from the NCBI protein database. We used ‘tblastn’ (e-value =$1e^{-3}$; max_target_sequences=5; and max_hsps=10) to query the protein sequences against the genome, and where possible, pulled down sequences from the species where the gene had been previously implicated in TSD. The majority of the gene sequences were sourced from *Trachemys scripta scripta*, *Chrysemys picta belli*, and *Alligator mississippiensis* (but see Table S11). Matches were then filtered downstream such that only sequences with ≥90% identity matches were retained, and positions of matches were checked against the annotation file. Results from grep and BLAST searches were then examined and compiled to create a comprehensive list of TSD genes for each of the two genomes. To compare the position of the genes within the genome, the positions of each gene were plotted on a Circos plot using CIRCA (http://omgenomics.com/circa).

*Genome-wide heterozygosity*

We used the 10X Genomics paired-end reads generated for the leatherback and green turtle reference individuals and aligned them back to their respective primary assembly to conduct analyses of genome-wide diversity and historical demography. To apply standard mapping and genotype calling pipelines to the data, we first removed the 10X linked barcodes from the raw reads using the script 'process_10xReads.py' (43) followed by quality trimming using Trimmomatic v 0.39 (44). Reads were aligned to the respective reference genomes with BWA-MEM v0.7.17 (45) using default parameters. PCR duplicates were then removed and read group headers were added with Picard-Tools v2.23.2 using the MarkDuplicates and AddOrReplaceReadGroups functions, respectively (http://broadinstitute.github.io/picard). The resulting alignment files for each species were used for all downstream analyses described below. As described above, we also included four and five additional individuals for the leatherback and green turtles respectively (see Table S13). To allow for direct comparisons, the data from these resequenced individuals were treated as described for the reference individuals, with the exception of the 10X pre-processing, as these data were generated using Illumina short-reads.

Genome-wide heterozygosity was calculated using an approach adapted from methods described in (46), and using the Genome Analysis Toolkit (GATK; v4.1.8.1 (47)). HaplotypeCaller was applied to identify and call loci in the emit reference confidence mode with base-pair resolution (-ERC BP_RESOLUTION), with the output GVCF file containing both variant and non-variant sites. Genotypes at each site were then generated from this output using GenotypeGVCFs, including at the non-variant sites. We removed unused alternate alleles from the genotypes using SelectVariants, and then filtered the VCF file based on depth of coverage (⅓× - 2× mean coverage) and genotype quality scores (MinQ = 20) at each site using an inhouse python script. We used the resulting filtered VCF file to visualize heterozygosity ($\pi$) in 100 Kb non-overlapping windows across the genome. To ensure the number of callable sites didn't influence our results, we calculated heterozygosity as the number of heterozygous sites divided by all sites that passed filtering steps, and only retained windows that contained a minimum of 80 Kb callable sites for each window. Heterozygosity estimates for regions without a known location in the genome (i.e. unplaced scaffolds) were not included in calculations. Overall heterozygosity was calculated as the total number of heterozygotes divided by the total number of callable sites across all windows. We also estimated heterozygosity for regions of the genome using the same methods as above, using an input BED file to specify the regions of interest. Specifically, we targeted regions that: (1) were not identified as containing repeat or low-complexity sequences (i.e. the 'masked genome', see *Transposable element analysis* section above), (2) were identified as exon regions through the annotation and (3) non-exon regions (i.e., regions not identified as exons, identified by inverting the exon region BED file using BedTools v2.29.2 (48). For the windows containing exons, we examined the genes associated with regions of high diversity by extracting the annotation information for windows that had a proportion of heterozygosity that was higher than 3× SD above the mean. Gene lists were then run through PANTHER (49) to investigate gene ontology (GO) terms.

To directly compare heterozygosity between the two sea turtle species, we also mapped the 10X barcode removed reads to the reference genome for *Mauremys reevesii* (50) using the same methodology as described above for alignment, duplicate removal and genotype calling as described above, using scaffolds that were at least 10 Mb in length (*N*=43, ~98% of the genome), and estimated diversity for whole-genome and exons. We then compared heterozygosity in corresponding exon windows for both species, and identified regions of high heterozygosity, as described above (i.e. windows that contained heterozygosity estimates greater than 3x the mean for each species were flagged as 'high'). These windows were subsequently subset depending on whether heterozygosity was high in both species, or only one species, such that they were sorted into either (1) substantially higher heterozygosity in one species than the other; or (2) exceptionally higher heterozygosity in both species. Following this identification, annotations of genes present in these windows were extracted and explored to determine differences between the two species.

To examine the context of the genomic diversity found in the two sea turtle species, we also directly estimated the genome-wide heterozygosity for a number of other reptile species (N=13). As the software and parameters used for genotyping can directly influence the heterozygosity estimates (see (51)), we downloaded raw reads associated with reference genome assemblies from the EBI-ENA database and employed a standardized mapping and genotyping pipeline to generate comparable heterozygosity estimates. The heterozygosity pipeline is similar to that described above for the two focal species with slight alterations: if data was generated with 10X Chromium linked-reads, the first 22bp of the R1 read were trimmed using Trimmomatic v0.39 (44). Following this, paired and trimmed reads were used as input for quality-trimming with Trimmomatic using default parameters, before being aligned to the reference genome with BWA-mem, having duplicate reads removed and read group headers added with Picard-Tools. The resulting alignment files were then used with the GATK pipeline described above, using 100 Kb windows, and only retaining scaffolds that were at least 100 Kb in length. Windows were discarded from downstream calculations if they contained fewer site calls than one standard deviation from the mean number of calls.

To determine the impact of genotype calling method on heterozygosity, we also generated genome-wide heterozygosity using the Analysis of Next Generation Sequencing Data software (ANGSD; v0.933(52)). To achieve comparable results to the GATK heterozygosity pipeline, we initially re-aligned the consensus genome around insertion-deletion (indel) sites using the RealignerTargetCreator and IndelRealigner functions included in GATK (v3.5), as this step is automatically included in the GATK analysis software (> v4.0). The indel realigned bam file was used as input for ANGSD, with site allelic frequencies calculated (-doSaf) using SamTools v1.9 (13) genotype-likelihoods (-GL1), and the same depth and quality filters as those applied in the GATK pipeline applied. Site allelic frequency files were then parsed through the realSFS function in ANGSD to calculate the site frequency spectra (SFS), with the outputs used to calculate heterozygosity within 100kb windows which were generated through bedtools MakeWindows function.

*Runs of homozygosity*

To detect autozygosity within the genome, we used the PLINK v 1.90b6.9 SNP-based runs of homozygosity (ROH) analysis (53). Briefly, we used ANGSD (52) to generate a SNP-list from the indel-realigned BAM files, with outputs in the form of a PLINK file, generated with a posterior probability cutoff of 0.95 and a SNP p-value of $1e^{-6}$. The ANGSD-generated SNP-list containing all WGR samples and the genome sample was then run through PLINK (--homozyg) to determine the length and distribution of ROHs across the genome for each individual using a minimum ROH length of 50 Kb (--homozyg-kb 50), a minimum of 20 SNPs (--homozyg-snp 20), an allowed missingness of 5 sites (--homozyg-window-missing 5), and a maximum of 1 heterozygous site allowed per window, to account for sequencing errors (--homozyg-window-het 1). The PLINK outputs were then exported and analyzed using the R environment (54). Given that limited literature exists on ROHs in reptiles (but see (55)), ROHs were segregated into length classes approximately based on those in (56), with 'small' ROHs classified as those between 50-500 Kb in length, ‘medium’ ROHs were 500 Kb-1 Mb in length, and ‘long’ ROHs >1 Mb in length. Total aggregate lengths were then calculated for each length class for each individual. Mean ROH length was compared between species using a T-test.

*Demographic history*

The demographic histories of leatherback and green turtles were inferred using the pairwise sequential Markovian coalescent (PSMC) (Li & Durbin 2011). PSMC analyses were run for all individuals for each species. For the reference individuals, 10X barcode trimmed reads were used as input, while for the resequenced individuals, reads were trimmed following the methods described previously, and all were aligned to their respective reference genomes using the BWA-mem pipeline described above. To process the data for PSMC we used samtools v1.11 (13) and bcftools v1.6 (57) to call variants, requiring base and mapping qualities of 30. We performed additional filtering by insert size retaining reads between 50-5000 bp, to remove potentially spurious short alignments. To mitigate the possibility of spurious heterozygotes we filtered by allele balance (AB), removing biallelic heterozygotes with AB<0.25 or AB>0.75 and filtered by repeat-masked positions. We retained the first 10 ‘SUPER’ scaffolds, which do not include any sex-linked chromosomes as sex-determining genes are not localized to discrete sex chromosomes in sea turtles. Following protocol (58), we retained sites between a third of the average read depth (-d) and twice the average read depth (-D). We applied PSMC using the parameters -N25 -t15 -r5 -p "4+25*2+4+6", and scaled the output using a mu of 1.2*10-8 (59) and a generation time of 30 years (which is the midpoint between literature estimates for the two species). We additionally plotted the PSMC outputs using species-specific generation times for each species, with values of 14 and 42.8 for leatherback and green turtles respectively. This scaling factor produced negligible impacts on the curves for $N_e$, with the 30-year generation time used for all downstream tests.

*Genetic load*

In order to examine deleterious allele accumulation and genetic load, we extracted variants from coding regions for all individuals of both species. We initially generated gVCFs for exonic regions individually using ‘HaplotypeCaller’ in GATK with a base-pair resolution and a minimum base quality of 20. Subsequently, gVCFs were imported into genomic databases using ‘GenomicsDBImport,’ before being joint-genotyped using ‘GenotypeGVCFs’ in order to obtain homozygous alleles for the reference individuals at additional SNPs identified in the resequenced individuals. In order to use these variants for the purpose of the genetic load analysis, the generated VCFs were split by sample using a simple bash script. Given that the VCFs contained all potential variants regardless of base quality and depth coverage, these were then filtered using VCFtools to retain only variants that were between 1/3 and 2x the mean coverage of each sample. These variants were then annotated using snpEff (61) with databases built for each species using the reference genomes and species-specific annotations, and where each variant was designated as producing either ‘modifier’, ‘low’, ‘moderate’, or ‘high’ impact. Proportions of each type of variant were then compared between and within species. SnpEff also calculated the silent to missense ratio of variants, with higher ratios showing a higher proportion of variants that are expected to have an effect on amino acid sequences.

### II. Extended Results and Discussion

*Analysis of regions of low synteny*

Here we provide in-depth descriptions of gene function and copy number comparisons between the two sea turtle species found in each region of low synteny. See Tables S6 and S8 for complete details. Two regions of low identity were identified on chromosome 1 from 1 Mb to 8 Mb for the green turtle and 1 Mb to 6 Mb for the leatherback turtle for region A, and from 210.8 Mbp to 214.4 Mbp for the green turtle and 215.7 Mb to 216.85 Mb for the leatherback turtle for region B. Inside region B, an unusual string of Ns was observed for the green turtle (51.2% of the total region length). The 3.5 Mb region was analyzed together with the same length section upstream and downstream for both green and leatherback turtles. The cactus alignment detected that both species exhibited more than 4 times duplications in this region, and the duplications are at least double in base-pair lengths, compared to surrounding regions (Table S6). We further selected only duplications larger than 21, 100, and 500bp for examination, and in all the cases the pattern remained the same for the region of low identity.

Additionally, there was a small increase in the amount of TEs for this region in the leatherback turtle (35;**46**;30 number of TEs in up to downstream order), but no difference in the green turtle (39;**35**;34 number of TEs in up to downstream order), possibly as a result of the high proportion of Ns in the green turtle for this region (Fig S9). Region A presented 59 genes with functions associated with Olfactory Receptors (OR) in the leatherback turtle, while the corresponding region for the green turtle presented a total of 256 OR gene copies (Table S8). The region B of chromosome 1 also presented multiple copies of three genes related to the Immune System (antigen WC1.1-like, TAPASIN and one gene containing Scavenger receptor cysteine-rich domain) for the green turtle compared to the leatherback turtle. We additionally checked for a possible association between the RRCs and TEs by comparing the RRCs with regions up- and down-stream, and found that the number of TEs was similar between these regions (Table S8). However, all large RRCs (> 1 Mb) in the green turtle that were associated with gene copy number differences had larger average TEs, potentially indicating an association of differential activity of TEs and structural differences in associations with gene copy number variations between species.

Two regions of low synteny were found on chromosome 2, region 2A (0 - 2.2 Mbp green turtle and 0 - 2.4 Mbp on the leatherback turtle) were associated with the presence of a duplication of one gene related to sphingomyelin phosphodiesterase 5 for the green turtle. The beginning of chromosome 4 also encompassed a region of low synteny (0 - 4.5 Mbp green turtle and 0 - 3.03 Mbp leatherback turtle) where multiple copies of genes related to the immune system (erythroid membrane-associated protein/butyrophilin and major histocompatibility complex class I) and one gene containing maestro-related heat domain were found for the green turtle. In chromosome 6, two low identity regions were identified at the beginning of the chromosome sequence. The first one (6A- and 0 - 15.47 Mbp green turtle and 0 - 7.67 Mbp leatherback turtle) contained potential gene duplication for genes related to olfactory receptors, the immune system and zinc-fingers for the green turtle compared to the leatherback turtle (see details in Table S6), while the second (6B) contained one gene of the immune system (NACHT 2C LRR and PYD domains-containing protein 3) with three copies on the green turtle compared to one on the leatherback turtle. The low synteny region on chromosome 8 (8A - 61.7 - 2.7 Mbp green turtle and 63.53 - 64 Mbp leatherback turtle) included the immune system gene complement factor H with 3 copies in the green turtle and 1 in the leatherback turtle. On chromosome 11, one region of low identity (11A - 74. 2 - 79.5 Mbp green turtle and 80.0 - 80.022 Mbp leatherback turtle) had multiple copies of zinc-finger genes for the green turtle compared to the leatherback turtle. Chromosome 12 presented a large inversion in the beginning of the chromosome; however, no signs of gene duplication were found for this region (3.004 - 7.090 Mbp green turtle and 3.296 - 7.396 Mbp leatherback turtle). As was found for chromosomes 1 and 6, multiple copies of genes related to the immune system and OR were found on a region of low synteny on chromosome 13 (13A - 32.3 - 42.95 Mbp green turtle and 33.3 - 41.16 Mbp leatherback turtle), and chromosome 14 (14A - 26.5 - 44.3 Mbp green turtle and 27.6 - 40.02 Mbp leatherback turtle). While the first region of low synteny identified on chromosome 15 did not present signs of gene duplication, the second region (15B - 13.7 - 14.3 Mbp green turtle and 13.3 - 13.6 Mbp leatherback turtle) had eight copies of one gene related to immunoglobulin lambda constant 1 for the green turtle compared with one copy for the leatherback turtle. Chromosome 20 presented duplication signs for genes related to Keratin type II head, adhesion G protein-coupled receptor E1 in the low synteny region 20A (4.9 - 14.1 Mbp green turtle and 4.8 - 14.7 Mbp leatherback turtle). The low synteny region found on chromosome 21 did not present signs of gene duplication. Chromosome 23 presented one of the larger regions of low synteny (6.0 - 19.3 Mbp green turtle and 5.9 - 17.23 Mbp leatherback turtle) with multiple copies of genes from immune system, reproductive system and iron homeostasis for the green turtle compared to the leatherback turtle. Additionally, chromosome 24 displayed rearrangements that were confirmed using 10X data as biologically real (Fig. S6; 24A - 12.2 - 19.2 Mbp green turtle and 11.6 - 16.95 Mbp leatherback turtle) containing multiple copies of genes from the immune system and maintenance of the mucosal structure (IGGFC-binding protein) again for the green turtle relative to the leatherback turtle. Finally, chromosome 28 was one of the largest low synteny regions, corresponding to the entire chromosome and included the presence of multiple copies of zinc-finger genes in the green turtle. All the genes present in multiple copies for the green turtle are shown in Table S6. The low synteny regions present on chromosome 2 (2B), 3 (3A), 5 (5A and 5B), 12, 15 (15A), 21, and 26 did not contain genes or signs of gene duplication. Other functions of genes with higher copies for the green turtle within RRCs included lipid metabolism (region 20A and 24A), cornification (region 20A), response to hypoxia (region 23A), and mucus production (region 24A).

*Repetitive elements*

Overall, the two genomes displayed similar percentages of repetitive elements (REs; 45.8% and 44.4%, respectively; Fig. S8 & Table S8), which consisted almost exclusively of transposable elements (TEs; 30.5% and 27.4%) and unclassified repeats (14.6% and 16.5%, respectively). While both genomes carried similar proportions of REs, the leatherback exhibited relatively longer TEs across all but two chromosomes (23 and 28), when compared to the green turtle (Fig. S9a). The landscape of TE superfamily composition over evolutionary time was also generally similar between the two species (Fig. S8), and consistent with other reptiles (51, 52). One striking difference between the sea turtle species, however, was seen in the REs with low Kimura values (<5%), which appeared at much higher frequency in the leatherback turtle (Fig. S8), representing either relatively recent insertions or reflecting a lower mutation rate in this species relative to the green turtle.

*Genome diversity*

Genome-wide nucleotide diversity was almost a magnitude of order lower in leatherback turtles compared to green turtles ($t(5.52) = 36.9$, $p < 0.01$; mean repeat masked $\pi = 2.86\times10\text{-}4$ and $2.46\times10\text{-}3$, respectively; Fig. 4a). Across regions, heterozygosity was lower in coding regions (mean $\pi = 2.77\times10\text{-}4$ and $2.18\times10\text{-}3$ for leatherback and green turtles, respectively; Fig. 4a) when compared to non-exonic regions (mean $\pi = 3.18\times10\text{-}4$ and $2.64\times10\text{-}3$; leatherbacks: [$t(4) = -8.9$, $p < 0.01$] and greens: [$t(5) = -30.9$, $p < 0.01$]), with the mean difference between coding and non-coding regions greater in the green turtles (19% compared to 13% in the leatherback turtle), likely due to the lower baseline heterozygosity in the leatherback turtle (Fig. 4a, main text). Further, the number of 100 Kb windows containing zero heterozygous sites was much lower in green turtles (mean = 540 windows) than leatherback turtles (mean = 2,363 windows; $t_{(8.21)} = -3.7$, $p < 0.01$); however the green turtle reference sample was a strong outlier within this species (n = 2,072 windows; Fig. S20), suggesting longer stretches of homozygous sites in this individual (see ROH results).

To identify genes with high diversity relative to baseline genome variation, we extracted exon-containing 100 Kb windows that had higher proportions of heterozygous sites than the mean for each species (see Methods) and identified 1,945 and 3,987 exons for the leatherback turtle and the green turtle, respectively (Table S15). Windows containing tRNA genes showed high heterozygosity for both species; however, the only specific genes observed in both species were *EPHA3* and *CHID1*, which encode an ephrin receptor and a response protein to excess calcium, respectively. Though a large proportion of the unique genes these exons comprise were with unannotated gene identifiers in both species (171 out of 302 for the leatherback turtle; 439 out of 506 for the green turtle), analysis of the annotated unique genes with PANTHER showed that the genes were involved with biological processes including development, locomotion, growth, response to stimulus and signaling (Fig. S21). The leatherback turtle also showed high diversity in genes associated with reproductive processes (Fig. S21). Examination of the annotated molecular functions from these exons revealed many with diversity in the leatherback turtle were related to cell adhesion, transport, and binding, while in the green turtle, they were associated with olfactory reception, immunity, tumorigenesis, and zinc finger proteins (Table S15).

When aligned to a common reference (*M. reevesii*) as opposed to themselves, we found similar results, with the diversity of the green turtle generally higher than the leatherback turtle (Fig. S18), albeit with a dampened difference between species (Table S15). In regions where diversity was high for both species (see Methods), many olfactory receptors were once again present, as were T-cell receptors, other immune-related genes (e.g. MHC related genes), maestro heat-like repeat-containing family members, and zinc finger proteins (Table S15). These were found for both species, but were especially prevalent for the green turtle which showed a greater number of high-diversity windows. For regions that were only indicated to have high diversity in the leatherback turtle, the genes within these regions were linked to some olfactory receptor genes, zinc finger proteins, and genes involved with signaling. Olfactory receptor genes were present in a higher number in the regions of high diversity in the green turtle, as were many immune-related genes, including genes linked to the MHC. Given the striking similarity to the RRC analysis results, these findings independently reinforced the importance of these gene families in the divergent evolution of these species. When compared to estimates from other non-avian reptiles generated using a standardized heterozygosity pipeline, we show that the leatherback turtle possesses very low genomic diversity (Fig. 4b), with estimates lower than even that of the well documented extinct *Chelonoidis abingdonii (62)*. The green turtle diversity falls midway between the other species, with estimates close to that of *Gopherus evegoodei* (1). Diversity did not correlate with the conservation status for the species examined.

*Searches for genes related to the core region of the MHC*

We further investigated immune genes associated with the core MHC region and found substantial differences between the leatherback turtle and the green turtle (Table S16). Out of the core set of MHC genes (63), 46 were present in the leatherback turtle and 39 in the green turtle, similar in number to those found in *Chrysemys picta bellii* and *Alligator mississippiensis* using the same gene set (63). Several genes were missing in both species, suggesting that either these genes have been lost in sea turtles, are too variable to be effectively annotated, or that this region still contains gap-rich regions.

Eleven genes present in the leatherback turtle genome were absent from the green turtle, including *BAG6*, *DDX39B*, *RNF5*, and *STK19*, but only four genes that were present in the green turtle versus the leatherback turtle (*KIFC1*, *LTA*, *TAP1,* and *TAP2)*. Excluding the MHC Class I and II genes, all core MHC-related genes were found on chromosome 14, except for *C4*, which was found on chromosome 1 in both species. In the green turtle, the *ATFB6*, *NOTCH4*, and *PRRT1* genes were additionally located on an unplaced scaffold (NW_025111287.1), while these were found on chromosome 14 in the leatherback turtle. This suggests that the assembled MHC region in the green turtle genome may be partly fragmented. Examination of MHC Class I genes suggested that multiple copies were present on chromosome 14 in both species (Fig. 2d), with seven copies found in the region for the leatherback turtle and six copies found for the green turtle, with an additional copy located on another unplaced scaffold (NW_025111276.1). There were two additional copies of the MHC Class I α gene in both species that were not located within the core MHC region on chromosome 14, with a single copy located on chromosomes 4 and 5.

*Conservation of reproductive genes and repetitive elements.*

In contrast to olfactory and immune genes, almost all genes with a priori linkages to TSD pathways (80–82) occurred as single copy orthologs with highly conserved chromosomal locations between the two species. Almost all 216 genes previously implicated in male- or female-producing pathways in reptilian species with TSD were single-copy genes in both sea turtle species (Table S11; 210 genes per species). Only three genes (*MAP3K3*, *EP300*, and *HSPA8*) were duplicated in both genomes, with the copies located on different chromosomes in all cases. Moreover, homologous genes were generally located in the same region of the genomes for both species (Fig. S12), and missing genes were typically absent in both species, with only four genes found in one species but not the other (Table S11).

This is likely indicative of strong selection for conservation of this reproductive pathway, but our understanding of the specific roles these genes play in sea turtle TSD remains limited. Resolving whether inter- (83) and intra-specific (84) variations in thermal thresholds are due to the few genes that diverged from the general pattern we observed, functional sequence variation between orthologs, or other factors (e.g., epigenetic processes) is of high conservation concern for sea turtles (85), as climate warming is expected to skew sex ratios and alter population demographics (86) in the absence of substantial plasticity or adaptation. Our results serve as the foundation for these much-needed studies to quantify genomic mechanisms of TSD in sea turtles and determine their adaptive capacity to persist under climate change.

While REs in turtles have been investigated for over 30 years (87, 88), few studies have directly addressed the distribution and diversity of REs within testudine genomes (89). Both sea turtle genomes have substantially larger RE compositions (>40%) than previous estimates for other turtle species (41, 89, 90), including the draft genome of the green turtle (10% of the genome (41)). Interestingly, more recent reptile genome assemblies show higher proportions of REs (90, 91), with results similar to our estimates. The benefits of whole-genome approaches are further highlighted in the tuatara, where initial RE estimates suggested <10% of the genome was composed of REs (92), yet a subsequent whole-genome assembly increased this estimate to 64% (45). Collectively, these results support the notion that RE patterns could be more conserved across non-avian reptiles than previously believed, and the continued application of recent advances in genome sequencing, assembly methods, and analyses are needed to better understand the RE patterns and the processes that generate them (39, 43).

### III. Supplemental Tables

**Table S1.** Summary assembly and gene statistics per scaffold. *See corresponding file*.

**Table S2.** Comparison of assembly statistics for rDerCor1 and rCheMyd1 before and after genome curation.

| | rCheMyd1 pre-curation | rCheMyd1 post-curation | rDerCor1 pre-curation | rDerCor1 pre-curation |
|---|---|---|---|---|
| Number of scaffolds | 129 | 92 | 96 | 40 |
| Total scaffold length | 2,160,271,820 | 2,134,358,617 | 2,166,457,950 | 2,164,762,090 |
| Average scaffold length | 16,746,293.18 | 23,199,550.18 | 22,567,270.31 | 54,119,052.25 |
| Scaffold N50 | 129,896,487 | 134,428,053 | 86,730,914 | 137,568,771 |
| Scaffold auN | 158,811,992.50 | 163,989,398.44 | 92,337,897.63 | 169,520,390.80 |
| Scaffold L50 | 6 | 5 | 9 | 5 |
| Largest scaffold | 343,335,227 | 348,265,484 | 149,920,125 | 354,452,888 |
| # contigs | 394 | 390 | 719 | 708 |
| Total contig length | 2,125,975,334 | 2,122,398,441 | 2,159,928,848 | 2,159,167,478 |
| Average contig length | 5,395,876.48 | 5,442,047.28 | 3,004,073.50 | 3,049,671.58 |
| Contig N50 | 45,429,622 | 39,415,510 | 7,029,801 | 7,029,801 |
| Contig auN | 52,823,488.45 | 46,322,642.98 | 9,107,684.00 | 8,982,433.65 |
| Contig L50 | 15 | 17 | 88 | 89 |
| Largest contig | 120,531,461 | 107,684,605 | 27,993,706 | 27,993,706 |
| # gaps | 265 | 298 | 623 | 668 |
| Total gap length | 34,296,486 | 11,960,176 | 6,529,102 | 5,594,612 |
| Average gap length | 129,420.70 | 40,134.82 | 10,480.10 | 8,375.17 |
| Gap N50 | 1,686,848 | 873,136 | 88,375 | 87,240 |
| Gap auN | 2,496,873.07 | 895,276.10 | 101,131.14 | 87,449.98 |
| Gap L50 | 6 | 5 | 27 | 25 |
| Largest gap | 6,200,714 | 1,638,132 | 349,742 | 153,563 |
| Base composition (A:C:G:T) | 594,891,663:467,921,984:467,881,242:595,280,445 | 594,093,851:466,992,810:467,080,695:594,231,085 | 611,710,726:468,217,715:468,140,090:611,860,317 | 611,482,842:467,997,451:468,033,727:611,653,458 |
| GC content | 44.02 | 44.01 | 43.35 | 43.35 |
| # soft-masked bases | 58,645,083 | 0 | 0 | 0 |
| # paths | 129 | 92 | 96 | 40 |

**Table S3.** Comparison of quality statistics for the genomes that exist for reptile species. *See corresponding file.*

**Table S4.** Comparison of assembly quality metrics for the draft green turtle genome (CheMyd1.0, Wang et al. 2013), the DNAZoo re-scaffolded assembly, and our new assembly (rCheMyd1).

| | **CheMyd1.0** | **CheMyd (DNAzoo)** | **rCheMyd1** |
|---|---|---|---|
| **Number of scaffolds** | 140,022 | 139,044 | 93 |
| **Number of contigs** | 274,366 | 275,152 | 391 |
| **Number of gaps** | 134,344 | 136,108 | 298 |
| **Scaffold N50** | 3,864,108 | 127,535,543 | 134,428,053 |
| **Contig N50** | 29,240 | 29,183 | 39,415,510 |
| **QV score** | 35.3307 | 35.3307 | 47.6069 |

**Table S5.** Nucleotide similarity levels based on whole genome alignment with the leatherback turtle (*Dermochelys coriacea*) genome used as the reference for Progressive CACTUS comparisons.

| **Genome** | **Nucleotide identity** | **Number of identical sites** | **Total number of aligned sites** |
|---|---|---|---|
| *Dermochelys coriacea* | 100 | 2,112,010,776 | 2,112,010,776 |
| Sea turtle ancestor lineage | 96.93 | 1,807,293,241 | 1,864,508,166 |
| *Chelonia mydas* | 95.35 | 1,712,413,459 | 1,795,899,214 |

**Table S6.** Interproscan functional analysis results of proteins present in multiple copies for the green turtle (*Chelonia mydas*) in the low synteny regions. *See corresponding file.*

**Table S7.** Gene collapse estimation for *Dermochelys coriacea* in the RRC regions.

| Region | Number of genes in assembly | Sum of average 10X coverages | Estimated number of genes from 10X coverage |
|---|---|---|---|
| R1A | 59 | 5,758.86 | 65.95 |
| R1B | 3 | 271.88 | 3.11 |
| R2A | 1 | 72.65 | 0.83 |

| | | | |
|---|---|---|---|
| R4 | 3 | 260.04 | 2.98 |
| R6A | 31 | 2,878.16 | 32.96 |
| R6B | 1 | 86.09 | 0.99 |
| R8A | 2 | 172.21 | 1.97 |
| R11A | 4 | 330.59 | 3.79 |
| R13A | 26 | 2,536.35 | 29.05 |
| R14A | 63 | 5,894.86 | 67.51 |
| R15 | 1 | 82 | 0.94 |
| R20 | 10 | 1,216.00 | 13.93 |
| R23A | 18 | 1,706.25 | 19.54 |
| R24A | 19 | 1,761.62 | 20.17 |
| R28A | 18 | 2,632.52 | 30.15 |

**Table S8.** Proportions of families and subfamilies of transposable elements in the green turtle (*Chelonia mydas*) and the leatherback turtle (*Dermochelys coriacea*) genomes. *See corresponding file.*

**Table S9.** Regions of reduced collinearity (RRCs) and their associated transposable elements (TEs) in comparison with adjacent high collinearity regions. *See corresponding file.*

**Table S10**. Olfactory receptor (OR) orthologs predicted by CAFE analysis. *See corresponding file.*

**Table S11.** Results of searches for genes associated with temperature-dependent sex determination (TSD) in both *Dermochelys coriacea* and *Chelonia mydas* genomes. *See corresponding file.*

**Table S12.** Genetic distance between the green (*Chelonia mydas*) and the leatherback (*Dermochelys coriacea*) turtles in reduced collinearity (RRC) and high synteny regions in macrochromosomes, small (<20 Mb) and intermediate (>20Mb) microchromosomes.

| | **Green turtle (*Chelonia mydas*)** | | **Leatherback turtle *(Dermochelys coriacea)*** | |
|---|---|---|---|---|
| | **RRC** | **High synteny** | **RRC** | **High synteny** |
| **Macrochromosomes** | 0.0501 | 0.0447 | 0.0509 | 0.0449 |
| **Microchromosomes >20** | 0.0526 | 0.0452 | 0.0545 | 0.0452 |

| | | | | |
|---|---|---|---|---|
| **Microchromosomes <20** | 0.0542 | 0.0524 | 0.0562 | 0.0524 |

**Table S13.** Information for additional whole-genome resequenced samples used in heterozygosity, runs of homozygosity (ROH), genetic load and demographic history analyses. Depth estimates are calculated after duplicates have been removed.

| **Sample ID** | **Original collection ID** | **Location** | **Tissue type** | **Sequencing technology** | **Alignment rate to reference** | **Mean depth (X)** |
|---|---|---|---|---|---|---|
| *Dermochelys coriacea* | | | | | | |
| Dc_WP1 (Reference) | rDerCor1 | California, USA | Whole blood | 10X linked reads | 98.1% | 76.59 |
| Dc_EP1 | dc_11171 | Mexiquillo, Mexico | Skin | Illumina | 99.8% | 17.35 |
| Dc_WP2 | dc_20292 | Lababia, Papua New Guinea | Skin | Illumina | 99.8% | 14.74 |
| Dc_NWA2 | dc_33126 | Gandoca, Costa Rica | Skin | Illumina | 99.6% | 14.39 |
| Dc_NWA1 | dc_ 101533 | St. Croix, US Virgin Islands | Skin | Illumina | 99.8% | 13.88 |
| *Chelonia mydas* | | | | | | |
| Cm_MED1 | rCheMyd1 (reference individual) | Israel | Whole blood | 10X linked reads | 98.3% | 48.33 |
| Cm_NWA1 | Flower | South Carolina, USA | Whole blood | Illumina | 99.9% | 10.48 |
| Cm_NWA2 | Yucca | Florida, USA | Kidney | Illumina | 99.8% | 48.16 |

| | | | | | | |
|---|---|---|---|---|---|---|
| Cm_NWA3 | Poppy | South Carolina, USA | Whole blood | Illumina | 99.9% | 13.96 |
| Cm_NWA4 | 27-2017-Cm | Florida, USA | Lung | Illumina | 99.9% | 53.52 |
| Cm_WP1 | CheMyd draft individual | Hong Kong, China | Unspecified | Illumina | 99.4% | 71.23 |

**Table S14**. Heterozygosity estimates for the leatherback (*Dermochelys coriacea*) and the green (*Chelonia mydas*) turtle genomes. Mean estimates were calculated from the individuals presented in Table S13 using the GATK SNP calling pipeline.

| Heterozygosity (heterozygous sites/bp) | *D. coriacea* (N=5) | *C. mydas* (N=6) | Ratio[#] |
|---|---|---|---|
| Genome-wide | $3.17\times10^{-4}$ | $2.61\times10^{-3}$ | 8.2 |
| Repeat and low-complexity masked | $2.86\times10^{-4}$ | $2.18\times10^{-3}$ | 7.6 |
| Exons | $2.77\times10^{-4}$ | $2.13\times10^{-3}$ | 7.7 |
| Non-exons | $3.18\times10^{-4}$ | $2.64\times10^{-3}$ | 8.3 |

#: Ratio of *C. mydas/Dermochelys coriacea*

**Table S15.** Genes associated with windows of relatively high diversity (mean(diversity) + 3*sd(diversity)) within the reference individuals of each species , and for regions within the *Maureyms reevesii* genome when *Dermochelys coriacea* and *Chelonia mydas* reference 10X reads were mapped to a common reference genome.

**Table S16**. List of immune-related genes of interest following Gemmell et al. (2020) and examined within annotations of both sea turtle species, with their corresponding positions in the genomes. *See corresponding file.*

**Table S17.** Variant rates, absolute values and percentages of variant groups, and missense to silent mutation ratios in coding regions of the leatherback and green turtles.

| ID | Number of variants | Number of effects | Variant rate | High impact | Moderate impact | Low impact | Modifier variants | High percentage | Moderate percentage | Low percentage | Modifier percentage | Missense to Silent ratio |
|---|---|---|---|---|---|---|---|---|---|---|---|---|
| dc_EP1 | 142,211 | 561,927 | 15,218 | 13,081 | 30,238 | 31,973 | 486,630 | 2.33% | 5.38% | 5.69% | 86.60% | 0.9816 |
| dc_WP2 | 143,294 | 566,813 | 15,103 | 12,843 | 30,384 | 31,988 | 491,592 | 2.27% | 5.36% | 5.64% | 86.73% | 0.984 |
| dc_NWA2 | 126,823 | 494,250 | 17,065 | 10,898 | 27,713 | 28,941 | 426,692 | 2.21% | 5.61% | 5.86% | 86.33% | 0.9868 |
| dc_NWA1 | 142,552 | 564,383 | 15,182 | 13,381 | 30,603 | 32,224 | 488,169 | 2.37% | 5.42% | 5.71% | 86.50% | 0.9811 |
| dc_WP1 | 157,348 | 620,286 | 13,754 | 14,644 | 32,788 | 33,846 | 539,001 | 2.36% | 5.29% | 5.46% | 86.90% | 0.9991 |
| cm_WP1 | 1,071,568 | 3,528,926 | 1,971 | 10,274 | 134,129 | 212,057 | 3,145,912 | 0.29% | 3.83% | 6.06% | 89.82% | 0.6959 |
| cm_MED1 | 1,149,778 | 3,785,501 | 1,837 | 12,044 | 141,048 | 219,192 | 3,380,517 | 0.32% | 3.76% | 5.84% | 90.08% | 0.7104 |
| cm_NWA1 | 1,053,798 | 3,474,840 | 2,005 | 10,665 | 131,325 | 207,974 | 3,097,630 | 0.31% | 3.81% | 6.03% | 89.85% | 0.6972 |
| cm_NWA2 | 1,122,555 | 3,709,995 | 1,882 | 11,478 | 138,486 | 219,056 | 3,311,363 | 0.31% | 3.76% | 5.95% | 89.97% | 0.6986 |
| cm_NWA3 | 1,105,243 | 3,652,545 | 1,911 | 11,222 | 136,054 | 215,544 | 3,260,538 | 0.31% | 3.76% | 5.95% | 89.99% | 0.6985 |
| cm_NWA4 | 1,124,679 | 3,716,742 | 1,878 | 11,638 | 138,139 | 219,138 | 3,317,605 | 0.32% | 3.75% | 5.94% | 89.99% | 0.6971 |

**Table S18.** Total number of homozygous and heterozygous variants that were predicted by snpEff. Note for the reference individuals, all variants will be heterozygotes except at SNPs/INDELs that were found in at least one other individual. (*) denotes reference individuals, (†) shows individual used to assemble draft genome (Wang et al. 2013).

| *Dermochelys coriacea* | | | *Chelonia mydas* | | |
|---|---|---|---|---|---|
| Individual | Homozygotes | Heterozygotes | Individual | Homozygotes | Heterozygotes |
| **dc_WP1*** | 19,081 | 64,001 | **cm_MED1*** | 2,651 | 299,284 |
| dc_WP2 | 34,580 | 38,194 | cm_NWA1 | 145,646 | 269,814 |
| dc_EP1 | 38,189 | 32,936 | cm_NWA2 | 157,708 | 319,904 |
| dc_NWA1 | 40,390 | 31,751 | cm_NWA3 | 151,319 | 298,870 |
| dc_NWA2 | 35,485 | 28,928 | cm_NWA4 | 156,464 | 324,725 |
| | | | cm_WP1† | 181,585 | 315,967 |

### IV. Supplemental Figures

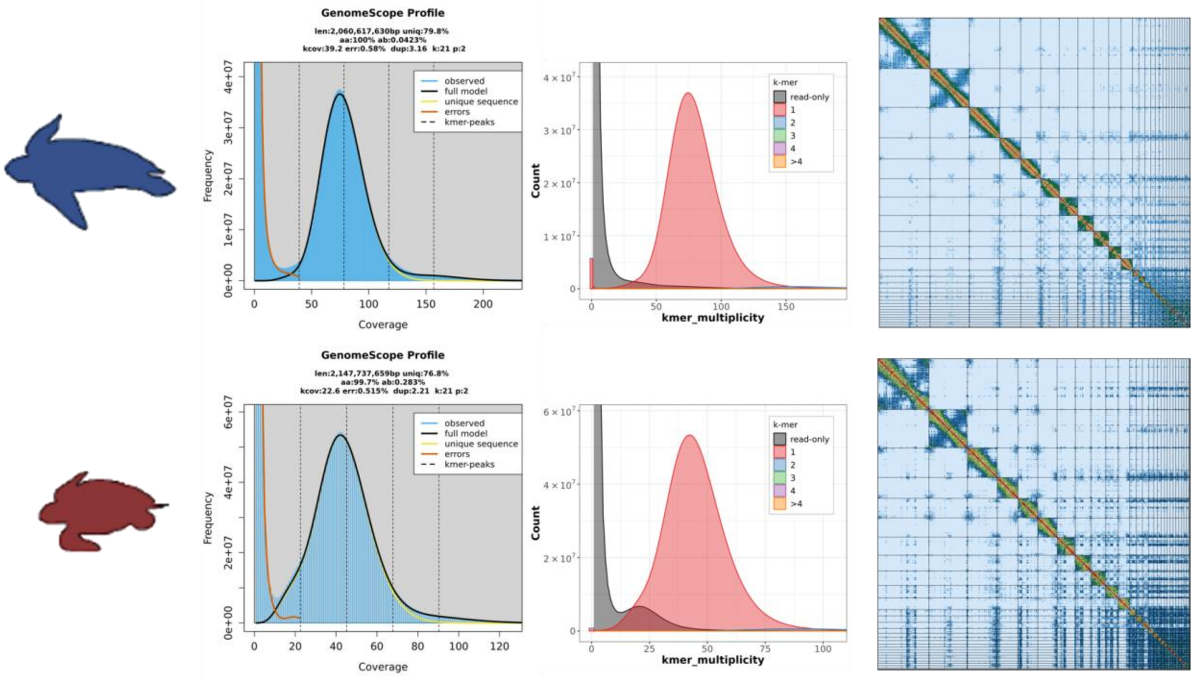


**Fig. S1 |** Quality control plots for the genome assemblies of *Dermochelys coriacea* (upper) and *Chelonia mydas* (lower) turtles. Plots from left to right; Genoscope profile for 21-mers collected from 10X linked reads using Meryl (https://github.com/marbl/meryl).; K-mer spectra plots for both genomes assemblies produced using KAT, showing the frequency of *k-mers* in the assembly versus the frequency of *k-mers* in the raw 10X linked reads. ; Hi-C maps contact map (Pretext https://github.com/wtsi-hpag/PretextView) for the complete assembly. Plots from left to right represent the kmer distribution profile from short reads (GenomeScope 2.0); the kmer multiplicity of reads coloured by the number of times each kmer appears in the assembly; and the contact map based on Hi-C short-read data produced using PreText.

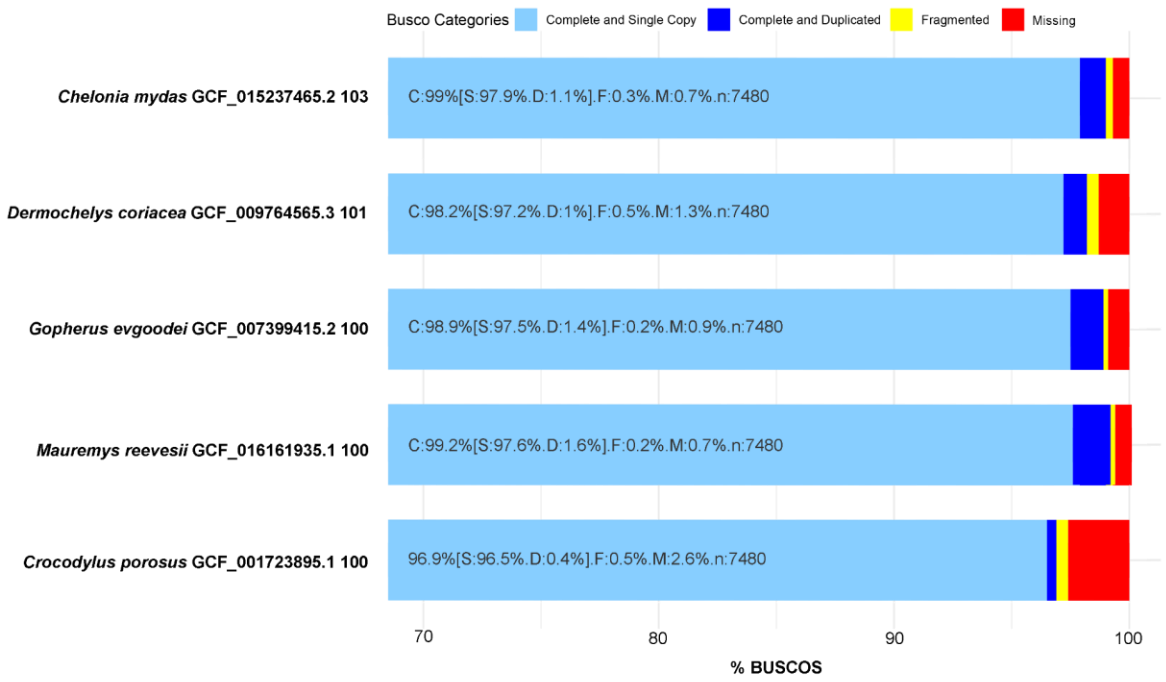


**Fig. S2 |** Comparison of the completeness of gene annotations, as a percentage of sauropsida_odb10 from BUSCO.

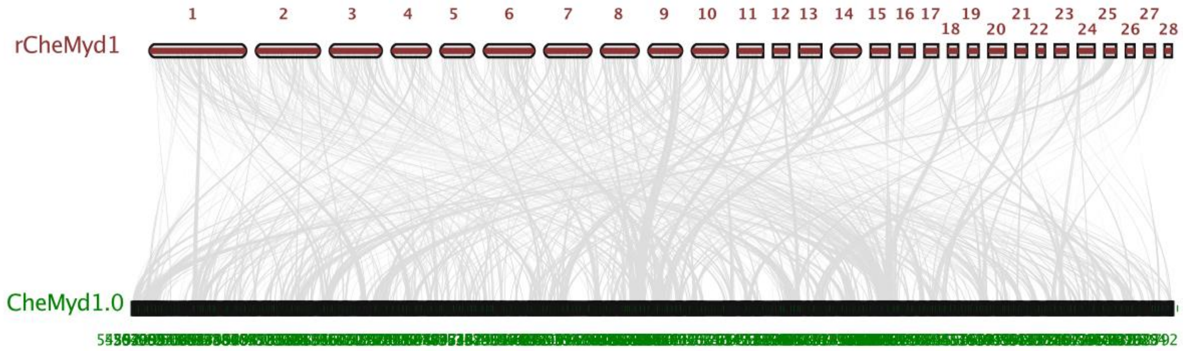


**Fig. S3 |** Gene synteny and collinearity per chromosome between the high-quality genome assembly rCheMyd1 (GCF_015237465.1 - red) and the draft genome assembly CheMyd1.0 (GCA_000344595.1 - green). Each bar represents chromosomes with respective numbers and gray lines represent homolog gene connections among species. It's not possible to visualize the 1520 bars from the CheMyd1.0 assembly.

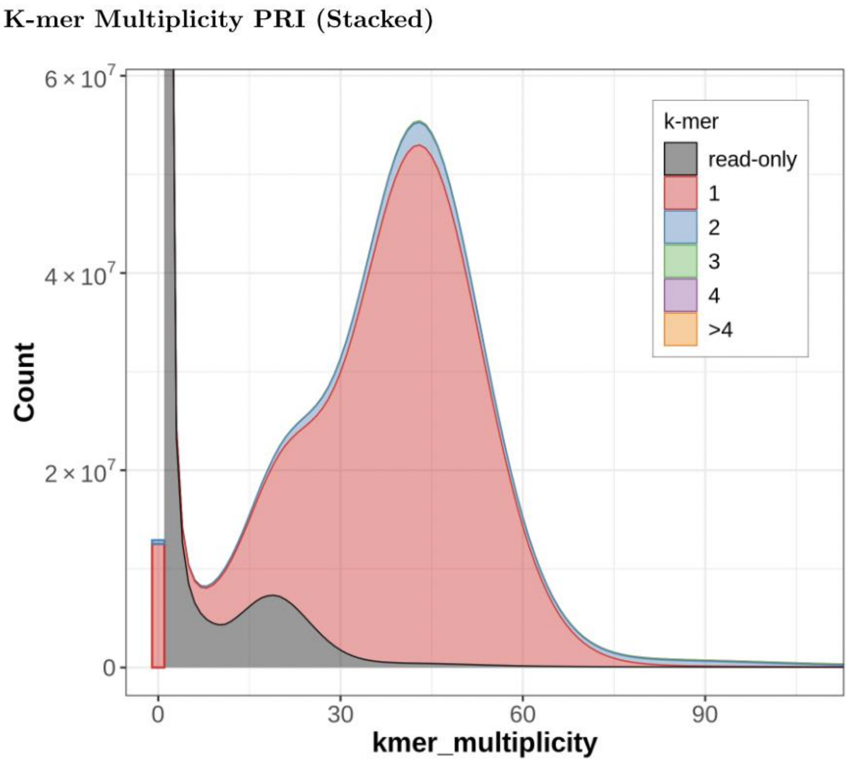


**Fig. S4** | K-mer multiplicity analysis (performed with KAT) for the draft genome of *Chelonia mydas* (CheMyd1.0; Wang et al. 2013).

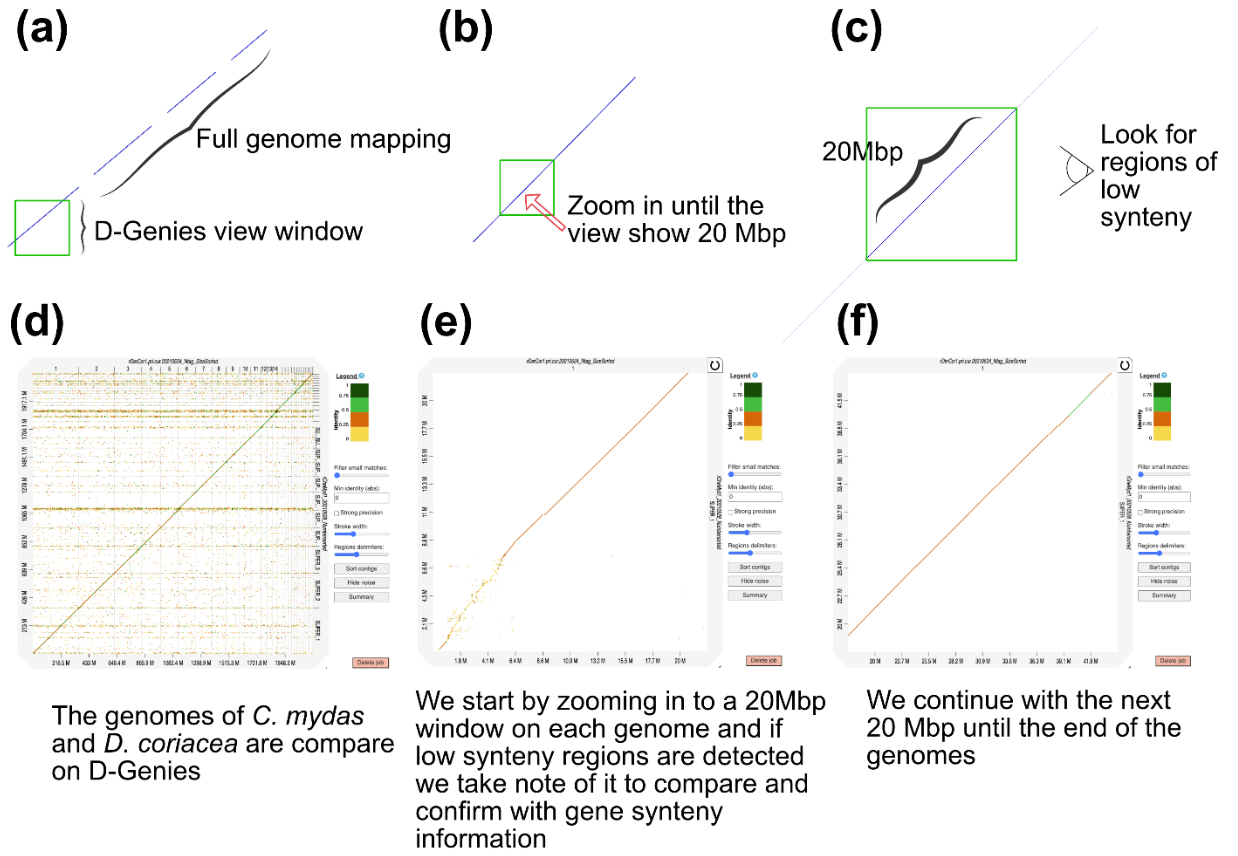


**Fig. S5 |** Schematic of the manual inspection of the RRCs using D-genies. The top panels (a-c) represent an ideogram of the process used, while panels d-f show the actual process using the D-genies software.

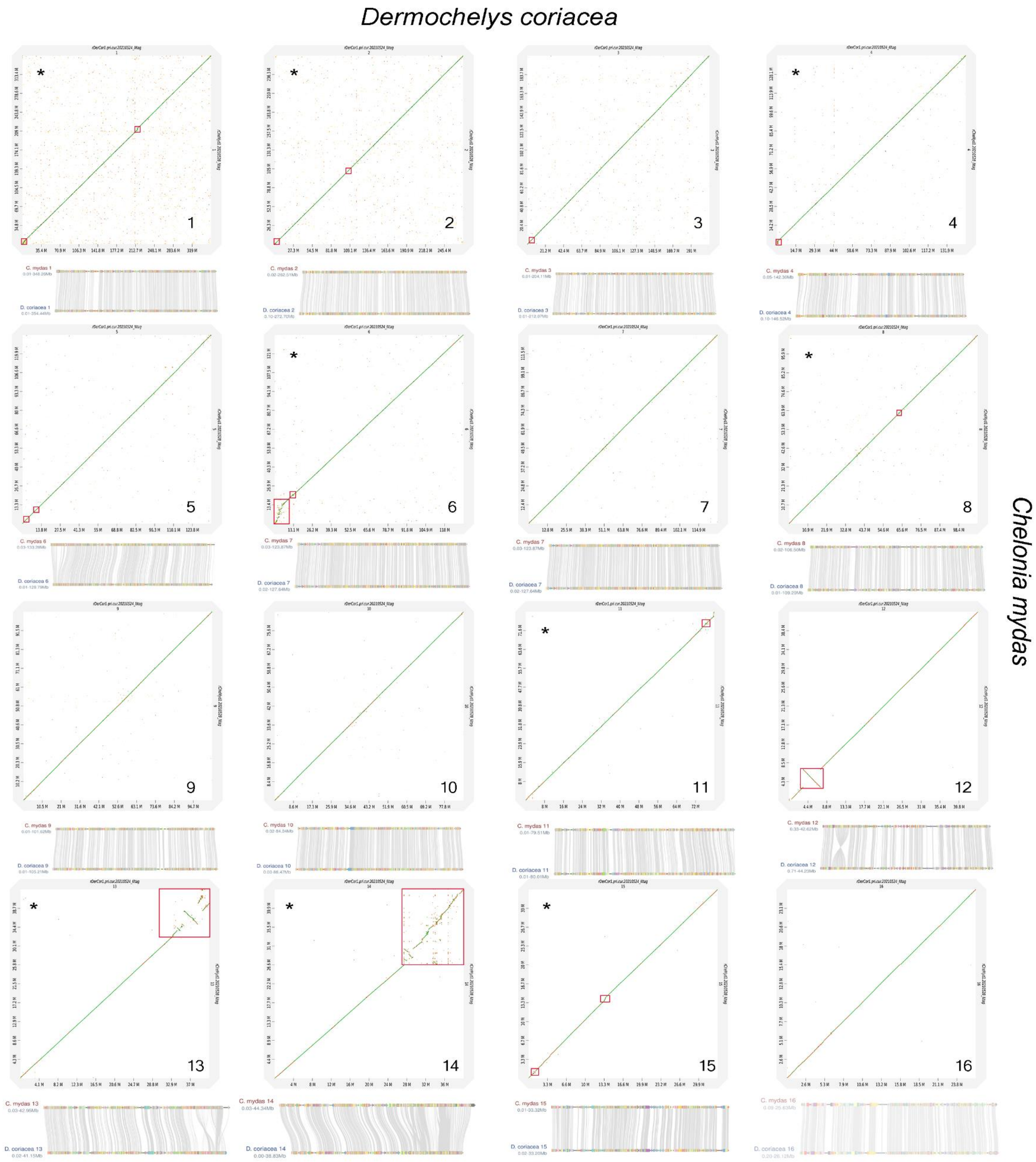
Dermochelys coriacea
Chelonia mydas

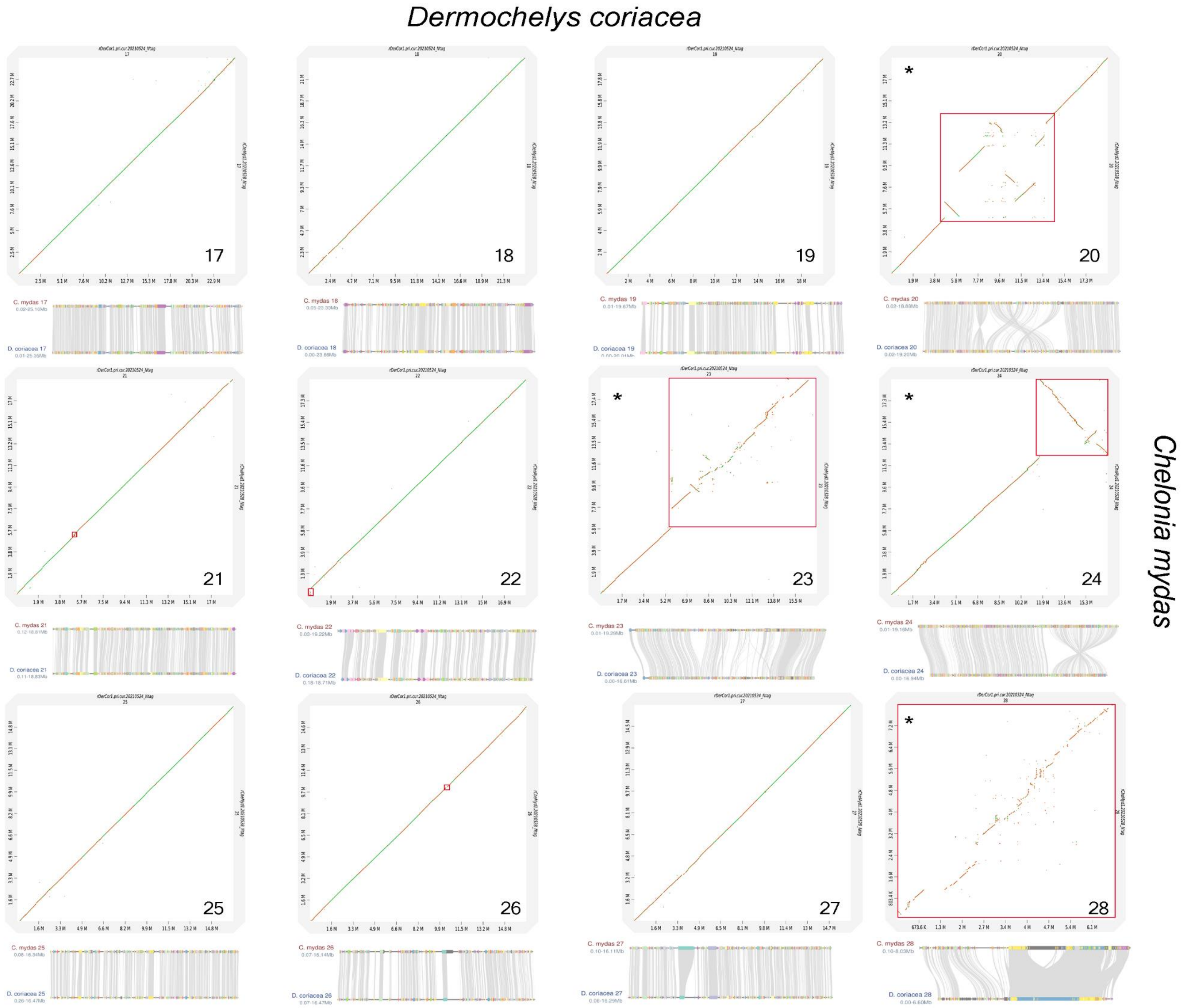


**Fig. S6 |** Dot plot analysis for all individual chromosomes in the leatherback turtle (*Dermochelys coriacea*) and the green turtle (*Chelonia mydas*) genomes, with identified regions of low synteny denoted by red boxes (top panel, each chromosome), and gene synteny analysis (bottom panel, each chromosome). The colored blocks with the same color in gene synteny graphs represent orthologous genes and the grey lines represent the links between them in the two species. At the genomic level, near end-to-end synteny was observed in 9 chromosomes (chromosomes: 7, 9, 10, 16, 17, 18, 19, 25, and 27), while from the remaining 19, 8 exhibited lower synteny restricted to specific sub-regions (>0.1Mbp - 3Mbp; chromosomes:  2, 3, 5, 8, 15, 21, 22, and 26), and 11 present low synteny regions larger than 3Mbp (chromosomes: 1, 4, 6, 11, 12, 13, 14, 20, 23, 24 and 28). Of the 19 chromosomes with regions of low synteny, the 13 that exhibited putative gene duplications within these regions are denoted by (*) in the upper left graph corner. The low synteny regions found on chromosomes 1, 4, 6, 8, 13, 14, 15, 20, 23, and 24 present multiple copies of genes related to immune system and/or olfactory reception in *C. mydas*. See details of region locations and compositions in Table S3.

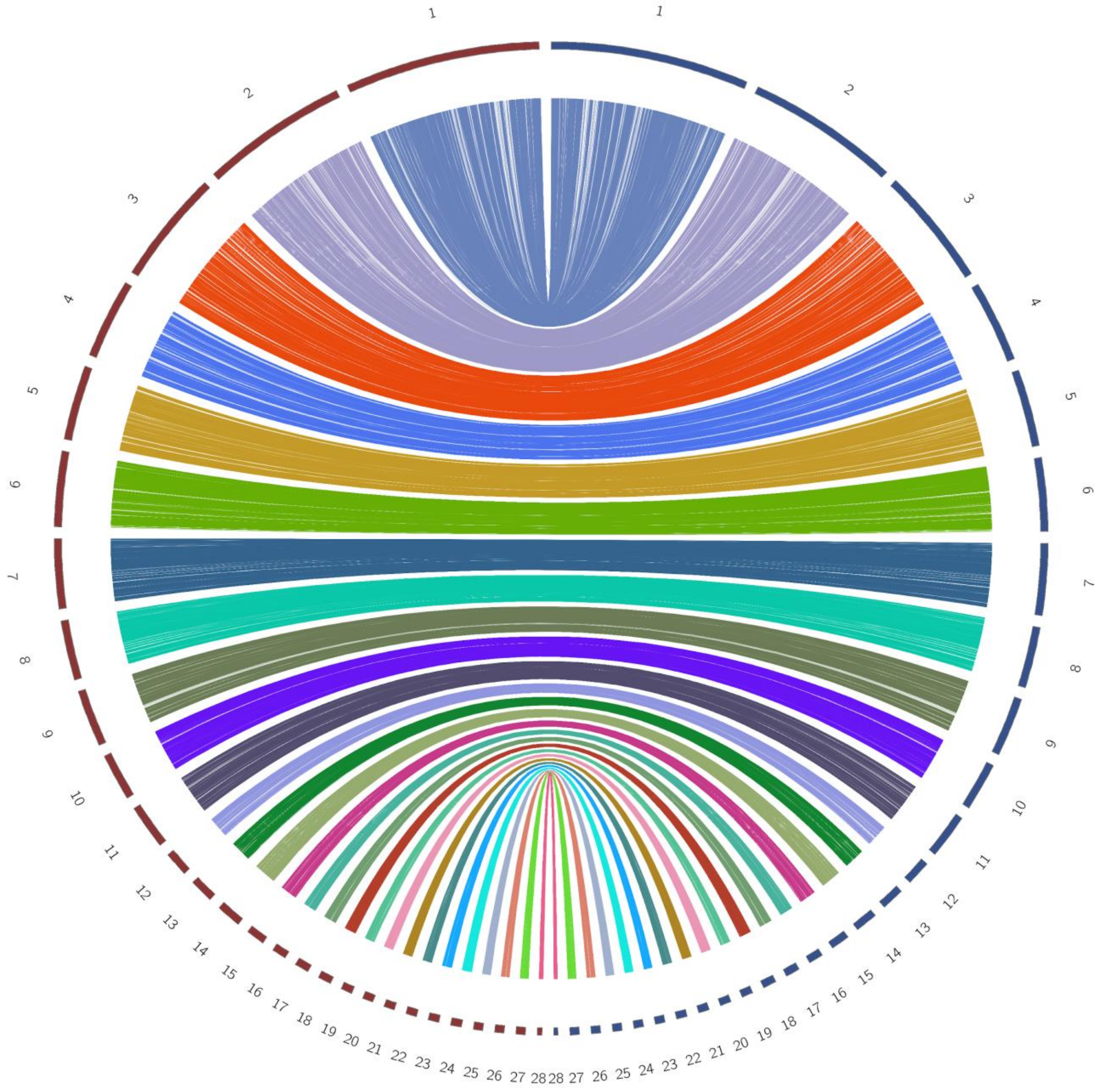


**Fig. S7 |** Circos plot for the genomes of the leatherback (*Dermochelys coriacea*) and the green turtles (*Chelonia mydas*) showing high gene synteny between species, with the outer rings showing respective chromosome numbers for *C. mydas* (red) and *D. coriacea* (blue).

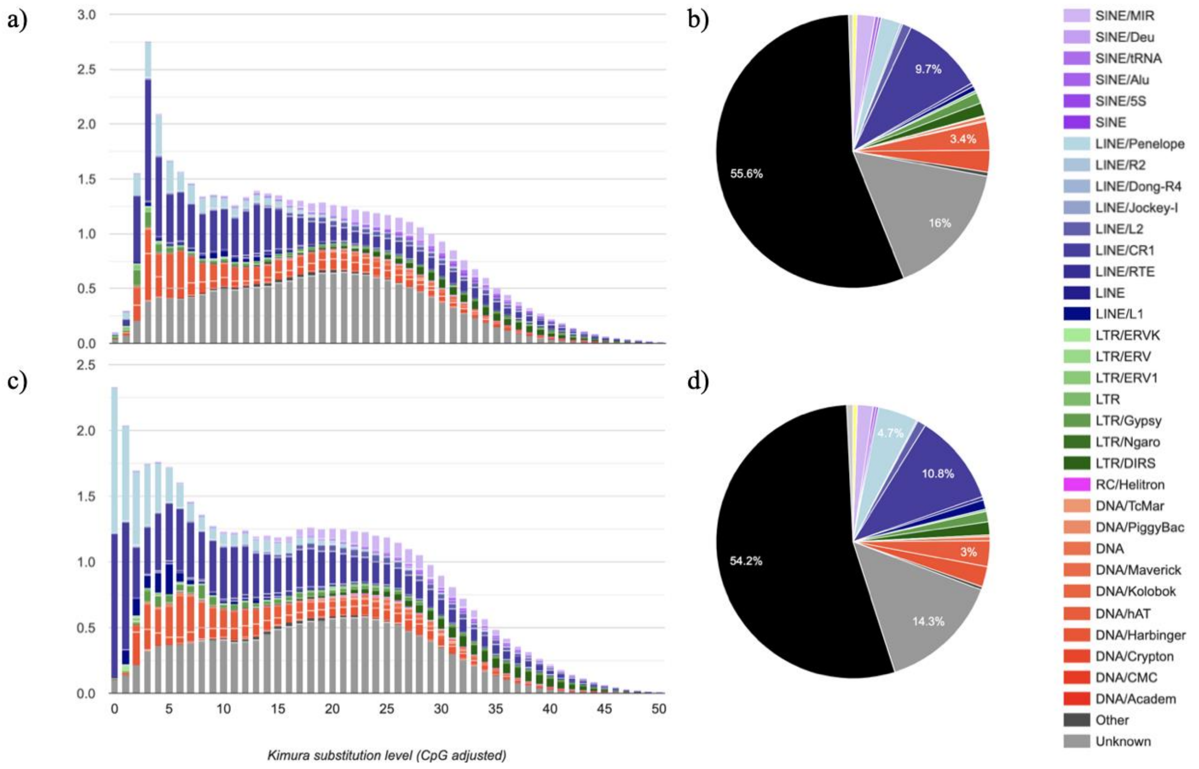


**Fig. S8** | Repeat element (RE) landscape for *Chelonia mydas* (a,b) and *Dermochelys coriacea* (c,d). Colors in the stacked bar charts and pie charts correspond to the transposable elements subfamilies and Unknown REs as indicated in the key, with the proportion of the unmasked genome depicted in black in b and d. See Table S4 for details.

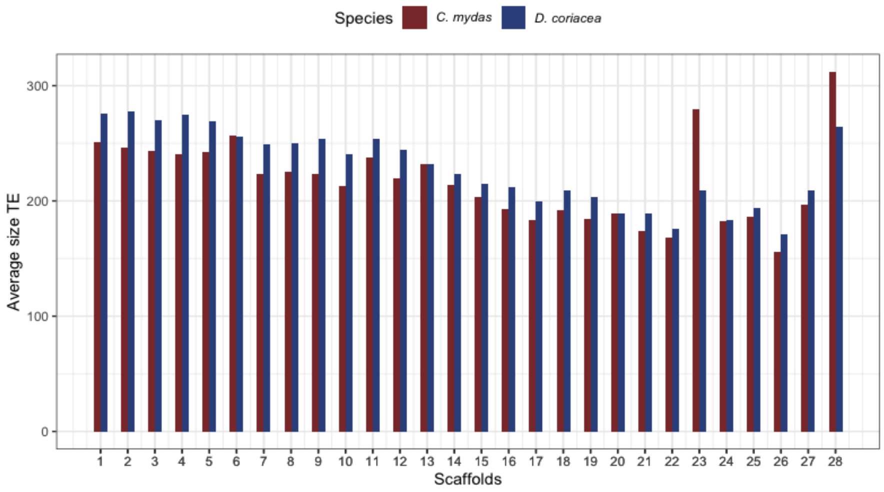
Species
C. mydas
D. coriacea
Average size TE
Scaffolds

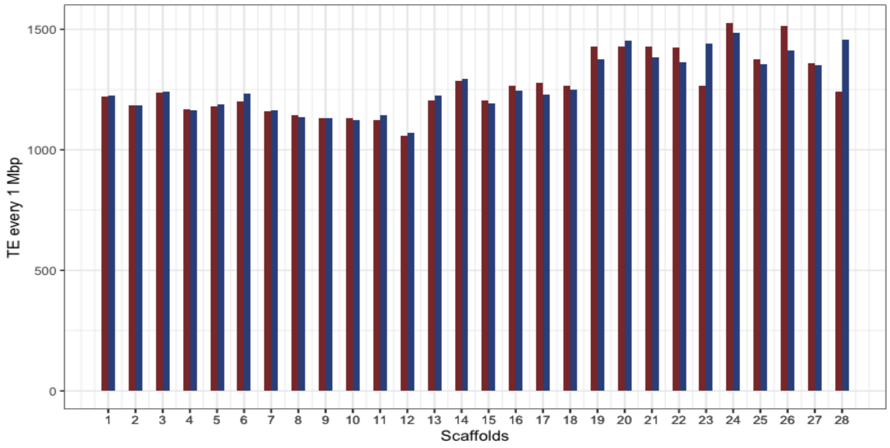
TE every 1 Mbp
Scaffolds

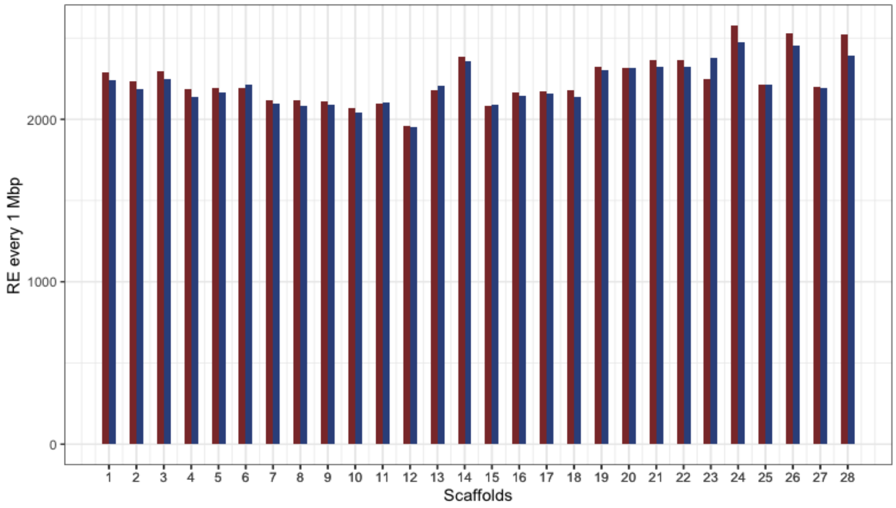
RE every 1 Mbp
Scaffolds

**Fig. S9 |** Distribution of **(a)** average size in bp of classified transposable elements (TEs), **(b)** number of TEs per 1 million bp and **(c)** number of all Repeat Elements per 1 million bp for each chromosome in *Chelonia mydas* (red) and *Dermochelys coriacea* (blue).

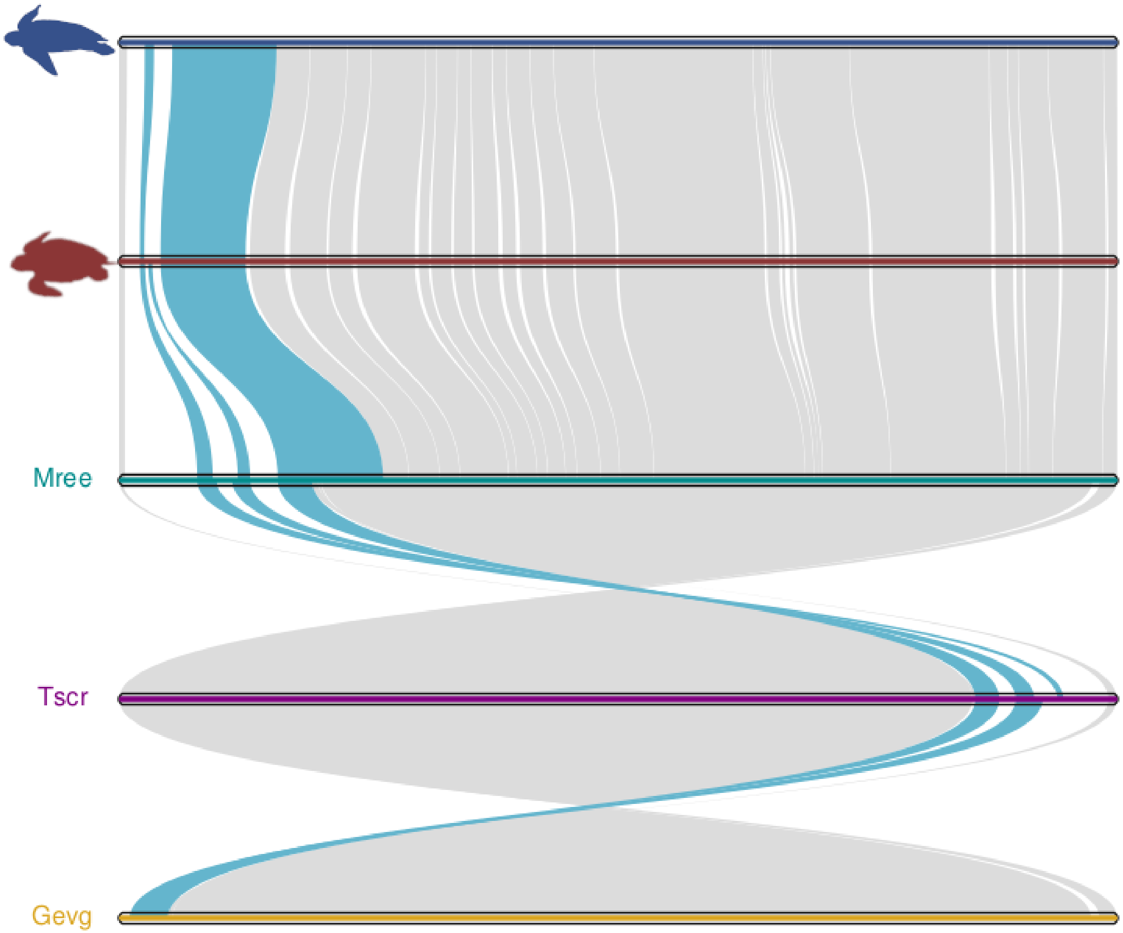


**Fig. S10 |** Comparison of Chromosome 1 homology across five turtle species depicting (cyan) the region with a cluster of Olfactory receptors class I. *Chelonia mydas* (red), *Dermochelys coriacea* (blue), *Mauremys reevesii* (Mree), *Trachemys scripta* (Tscr) and *Gopherus evgoodei* (Gevg).

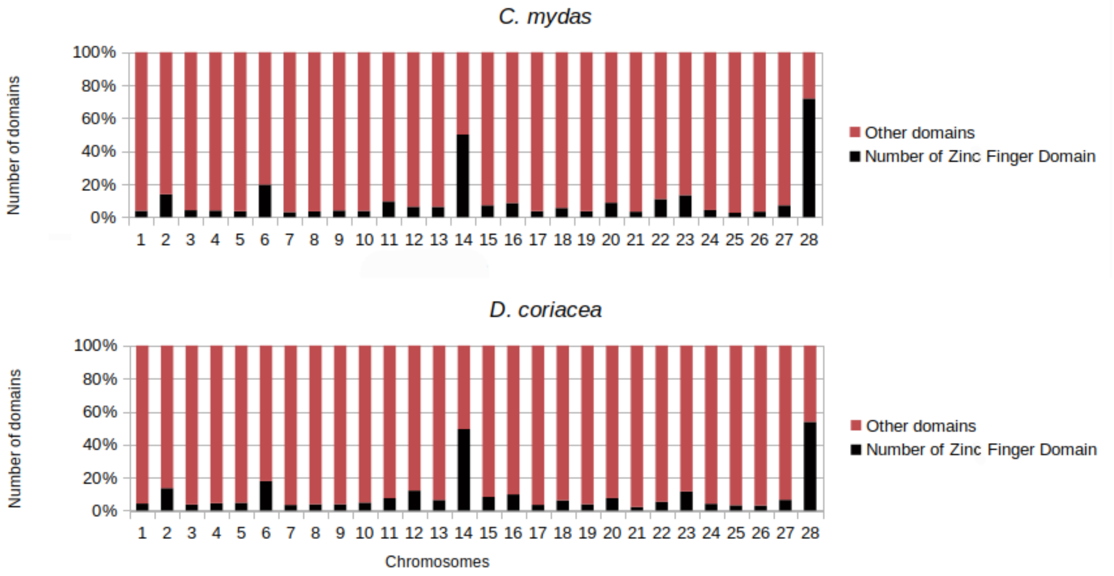


**Figure S11** | Proportion of Zinc finger domains per chromosome for the green turtle (*Chelonia mydas*) and the leatherback turtle (*Dermochelys coriacea*). A concentration of Zinc finger domains can be observed in chromosomes 6, 14 and 28 for both species.

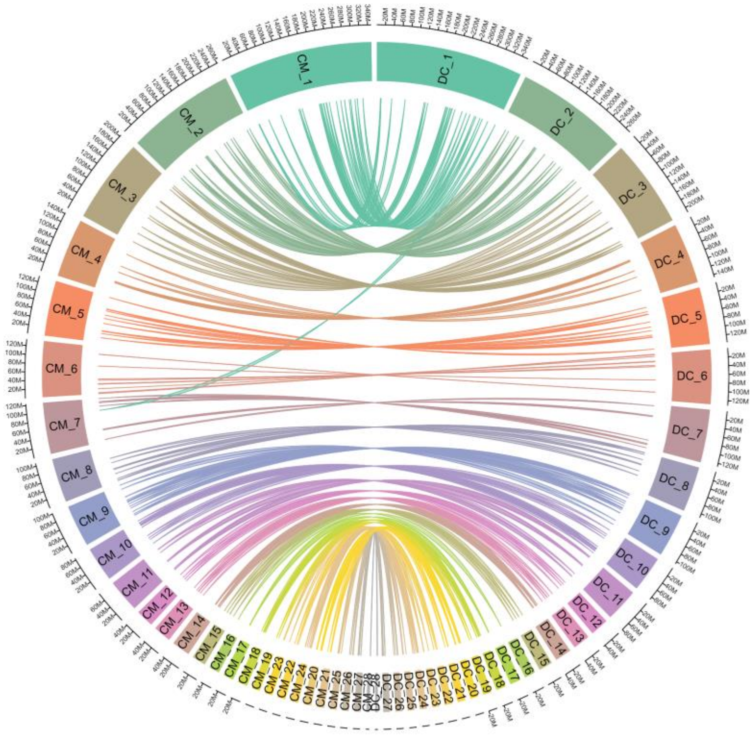


**Fig S12** | Locations of 213 genes that have been implicated in temperature-dependent sex determination and that were located in the genomes of both species of sea turtle (green turtle (*Chelonia mydas*): left; leatherback turtle (*Dermochelys coriacea*): right).

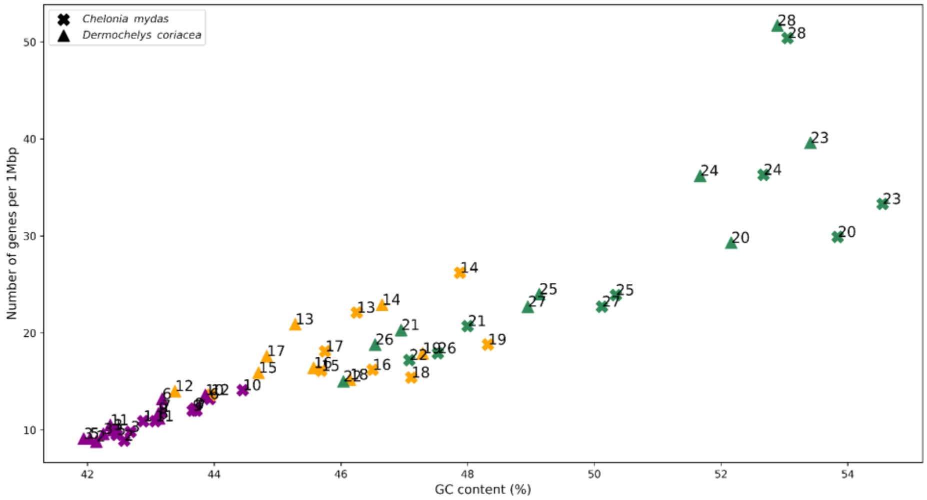


**Fig S13** | Relation between number of genes per 1 Mb and GC content for *Chelonia mydas* and *Dermochelys coriacea*. Macro-chromosomes are grouped in purple, micro-chromosomes with >20 Mb in orange and micro-chromosomes with <20 Mb in *C. mydas*.

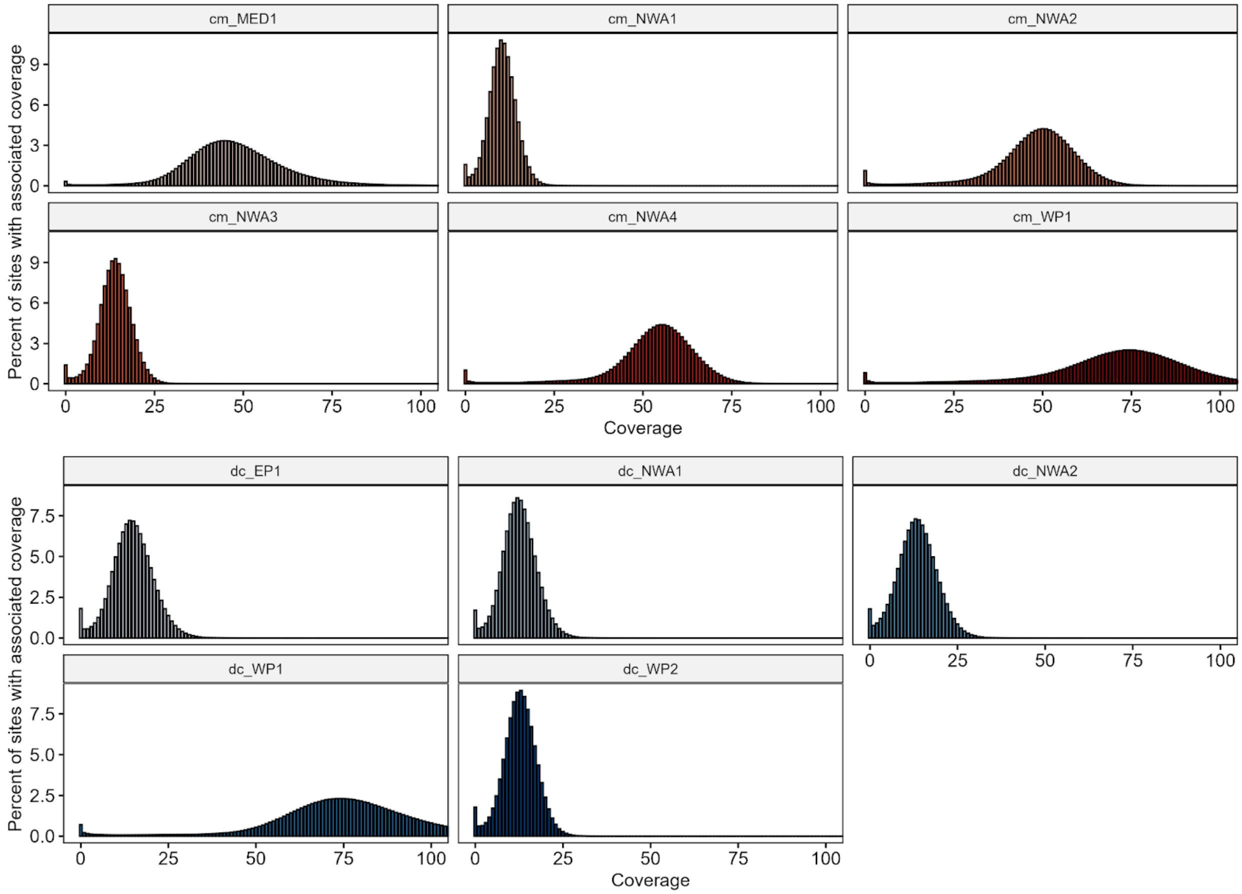


**Fig. S14 |** Depth coverage distribution for the two reference individuals (10X linked-reads; cm_MED1 and dc_WP1), as well as the additional resequenced individuals (Illumina short reads).

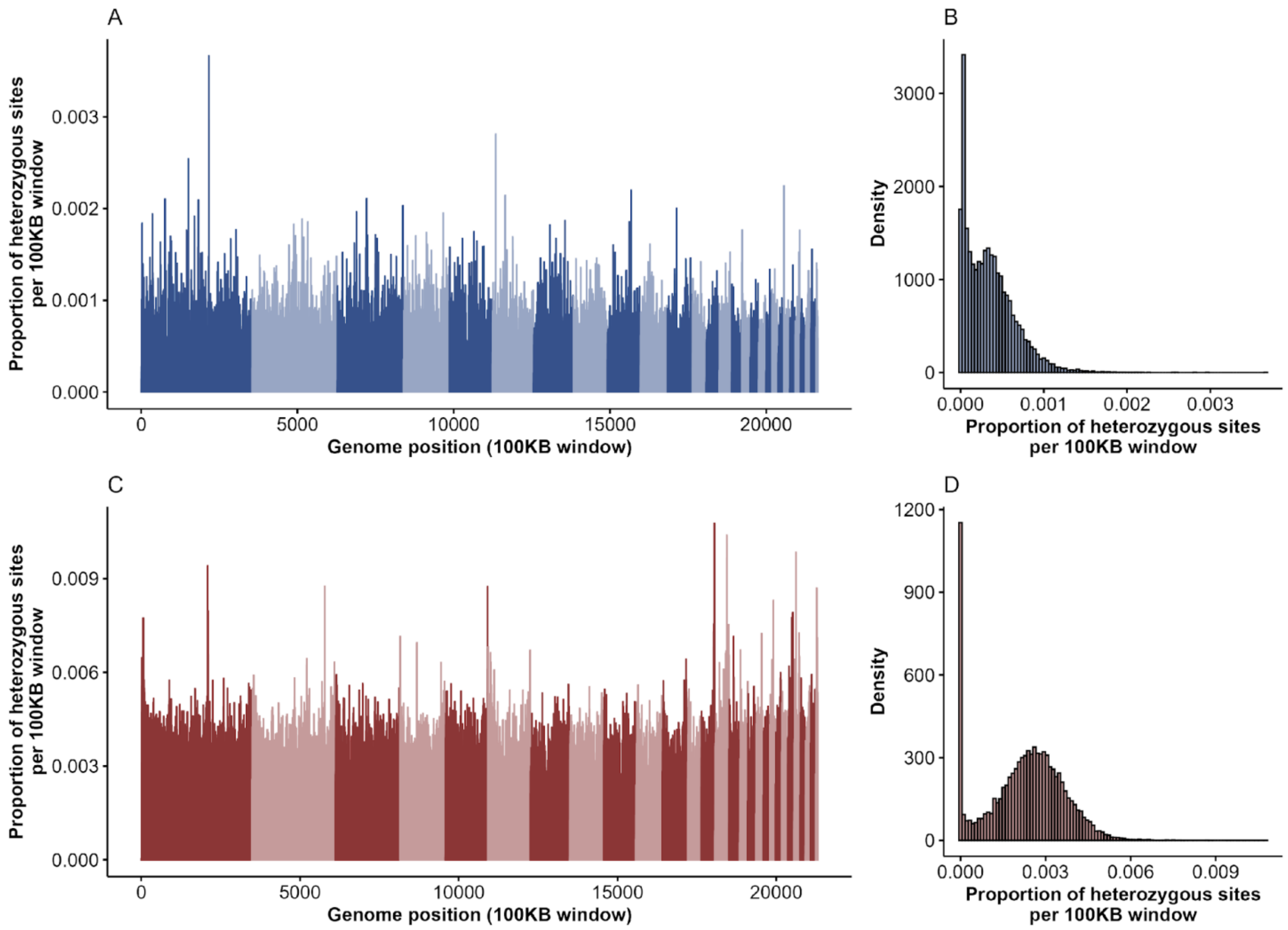


**Fig. S15 |** Genome-wide heterozygosity plots generated through GATK for both *Dermochelys coriacea* (A, B) and *Chelonia mydas* (C, D) reference individuals for the known 28 chromosomes. Both (A) and (C) show the proportion of heterozygous sites in 100 Kb windows where at least 90% of the sites were callable. Alternating colors show breaks between chromosomes. Plots (B) and (D) are histograms displaying the relative density of windows with associated heterozygous proportions. Note that the mean genome-wide heterozygosity estimates are approximately 6.5-times higher for *C. mydas*.

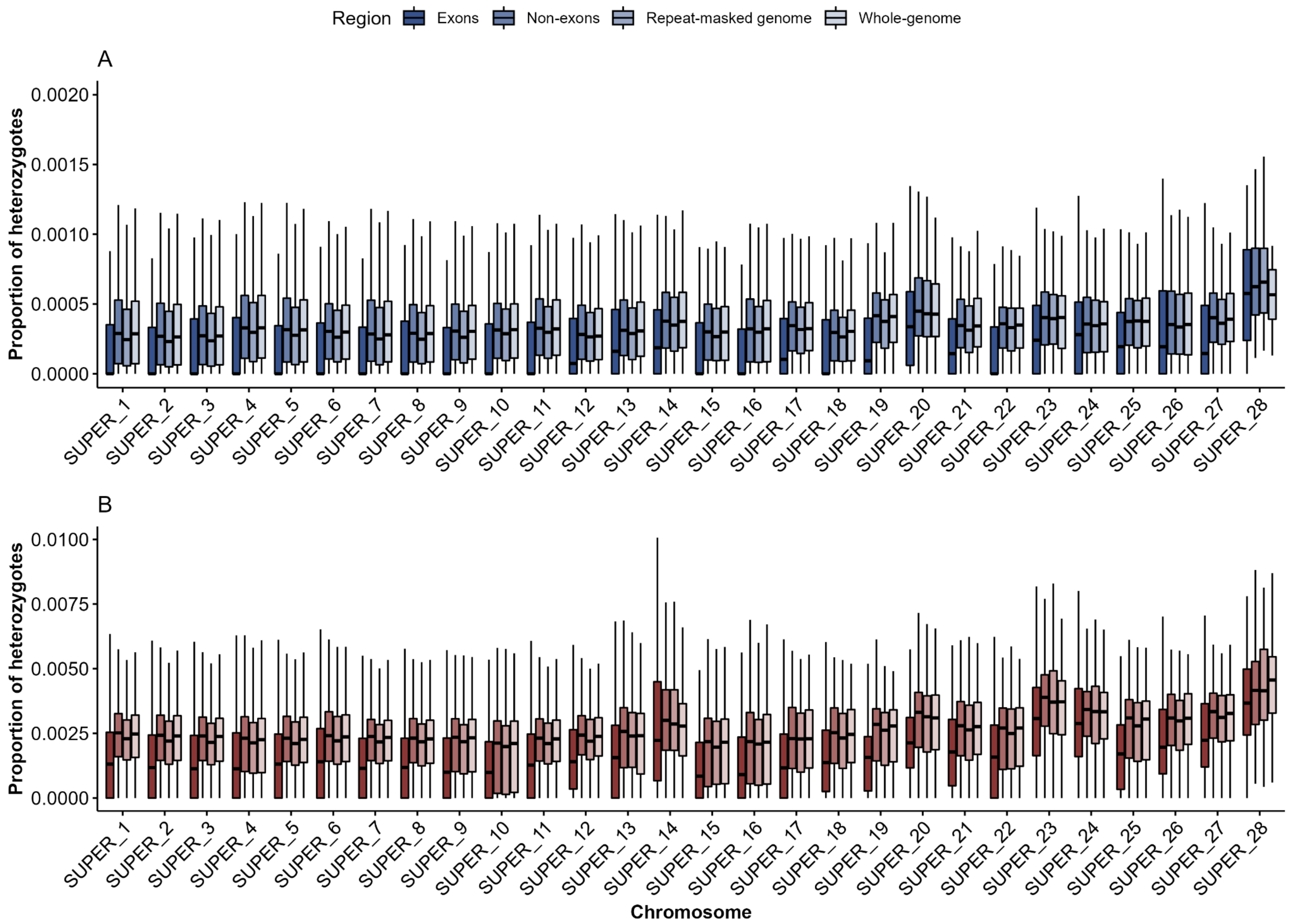


**Fig. S16 |** Chromosome-specific estimations of diversity for whole-genome, repeat-masked, exon, and non-exon regions for the reference individuals of both species.

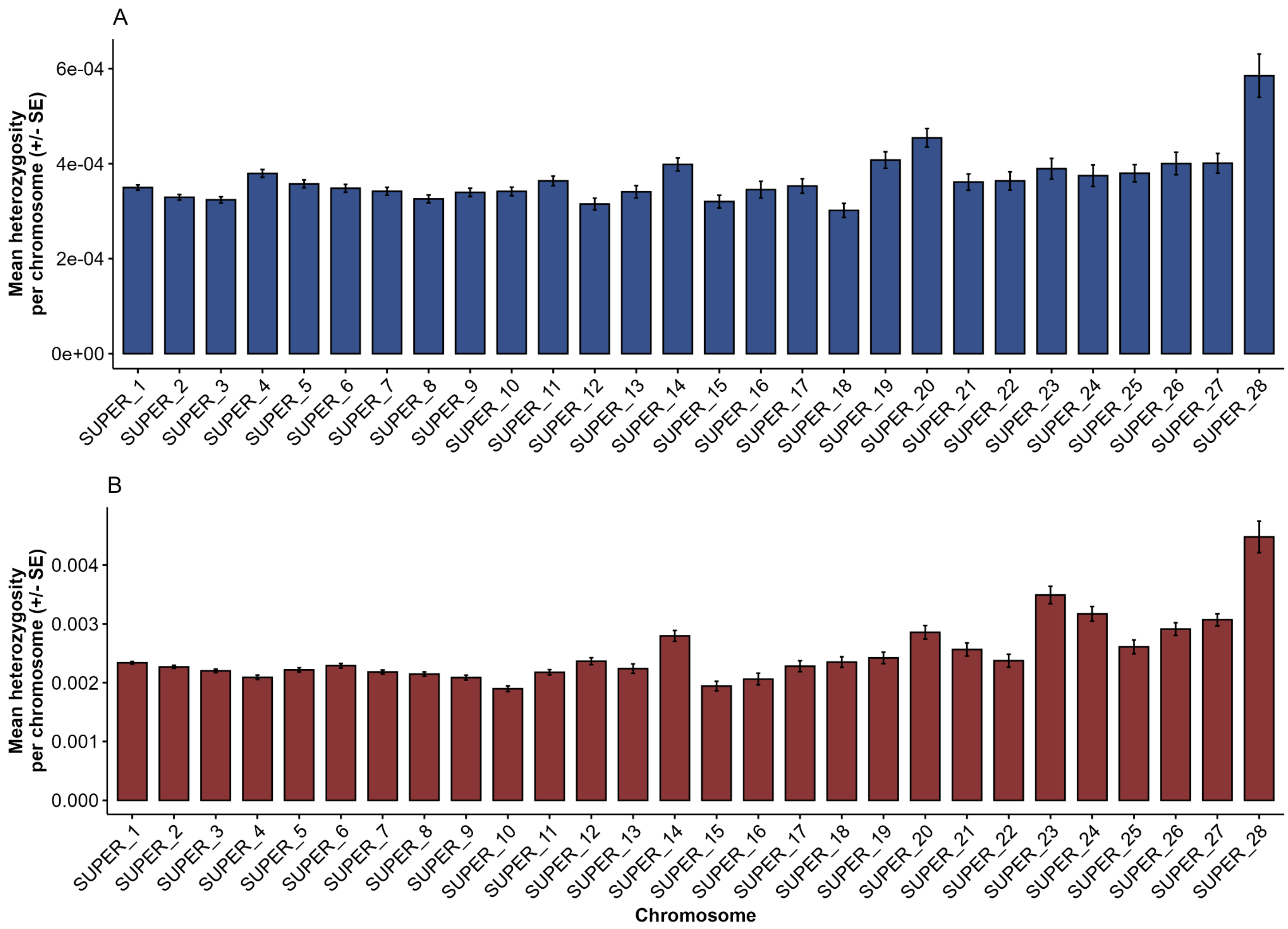


**Fig. S17** | Mean heterozygosity per chromosome (+/- SE) for the reference individuals of *Dermochelys coriacea* (A) and *Chelonia mydas* (B).

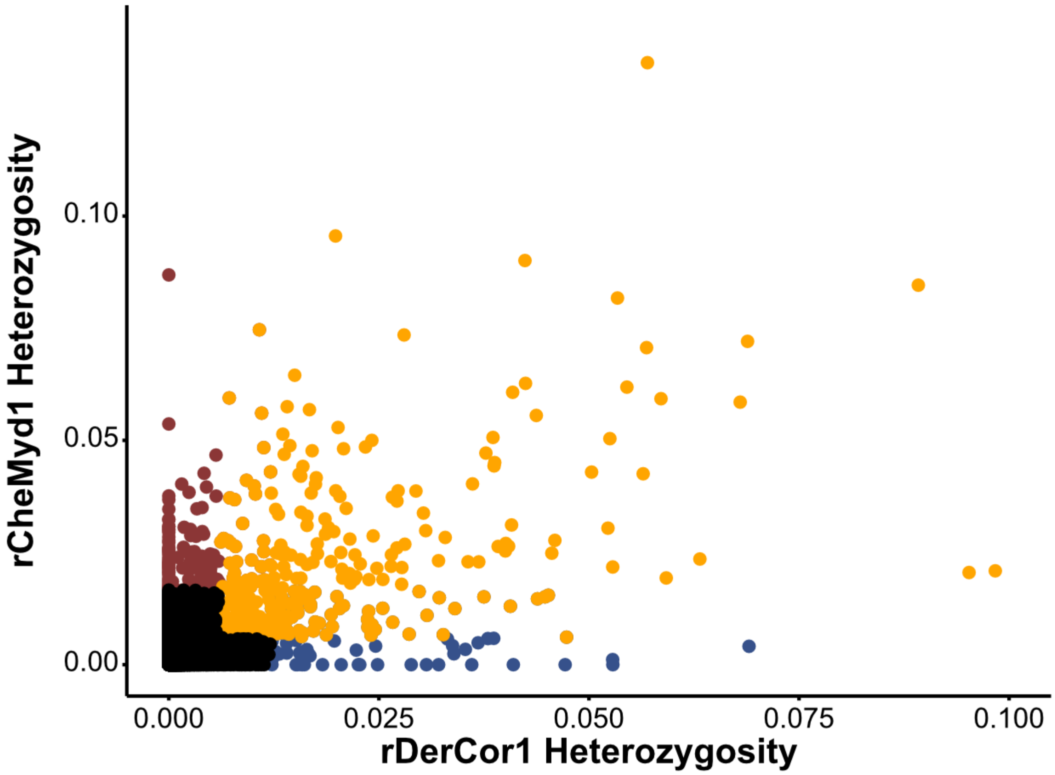


**Fig. S18 |** Correlation between heterozygosity in 100 Kb windows containing only exons, generated through alignment to a common reference genome. Windows with higher than mean diversity in leatherbacks (blue), higher in greens (red), and generally high diversity (orange) are highlighted.

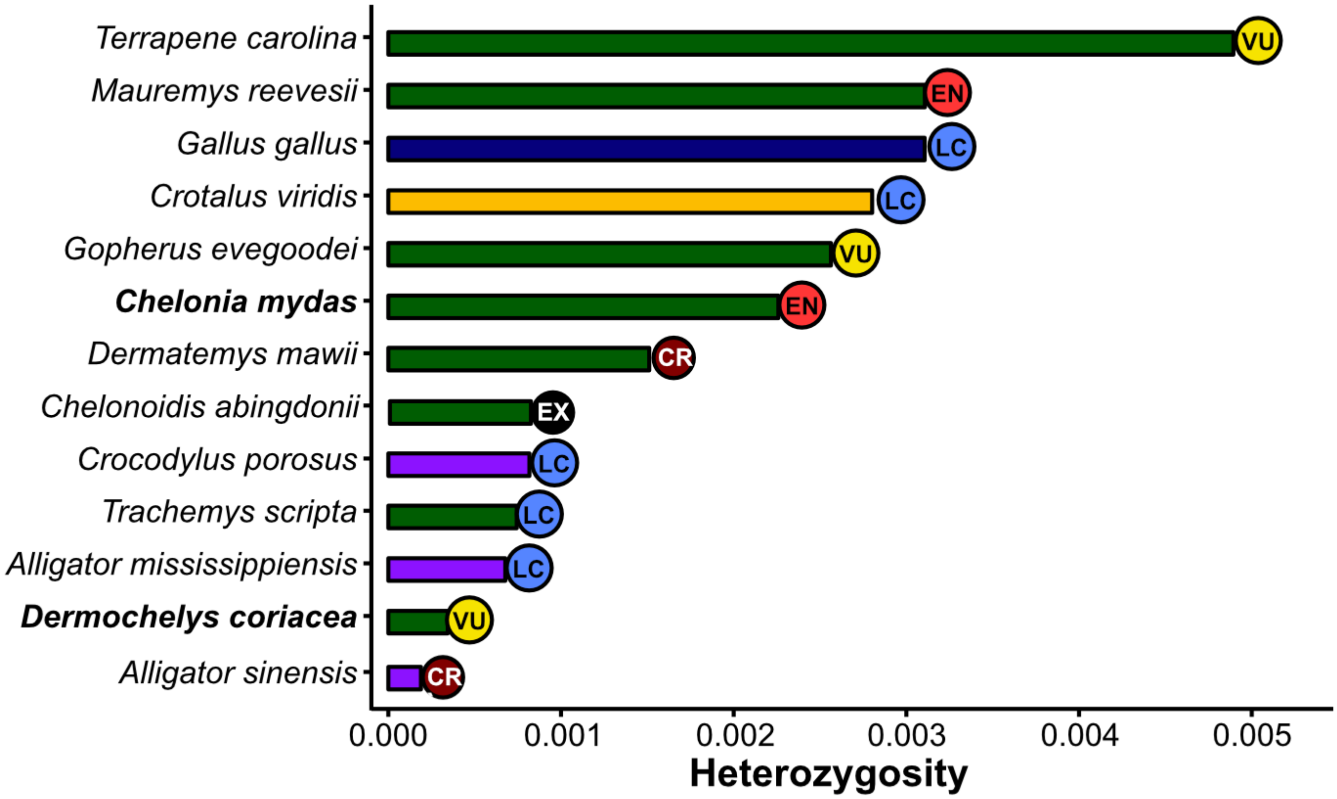


**Fig. S19** | Comparison of genome-wide heterozygosity for the reference individuals of leatherback (*Dermochelys coriacea*) and green (*Chelonia mydas*) turtles in relation to other reptile species where assembled genomes are available, and their associated IUCN conservation status. Green bars depict turtle species, with purple bars representing crocodilians, yellow bars showing squamates, and avians shown by navy blue bars.

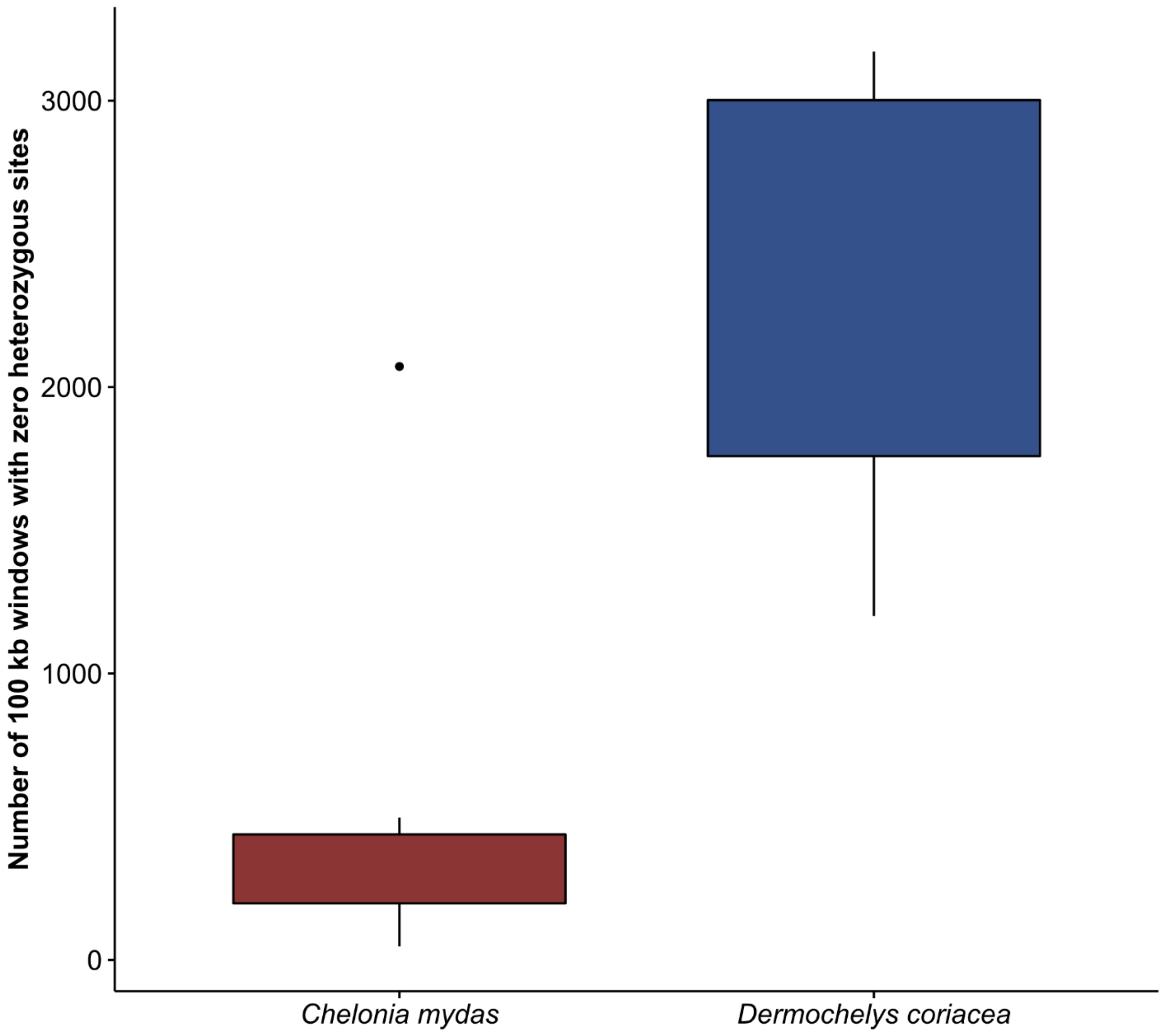


**Fig. S20 |** Total number of 100 Kb windows containing zero heterozygous sites for both green (*Chelonia mydas*) and leatherback (*Dermochelys coricea*) turtles.

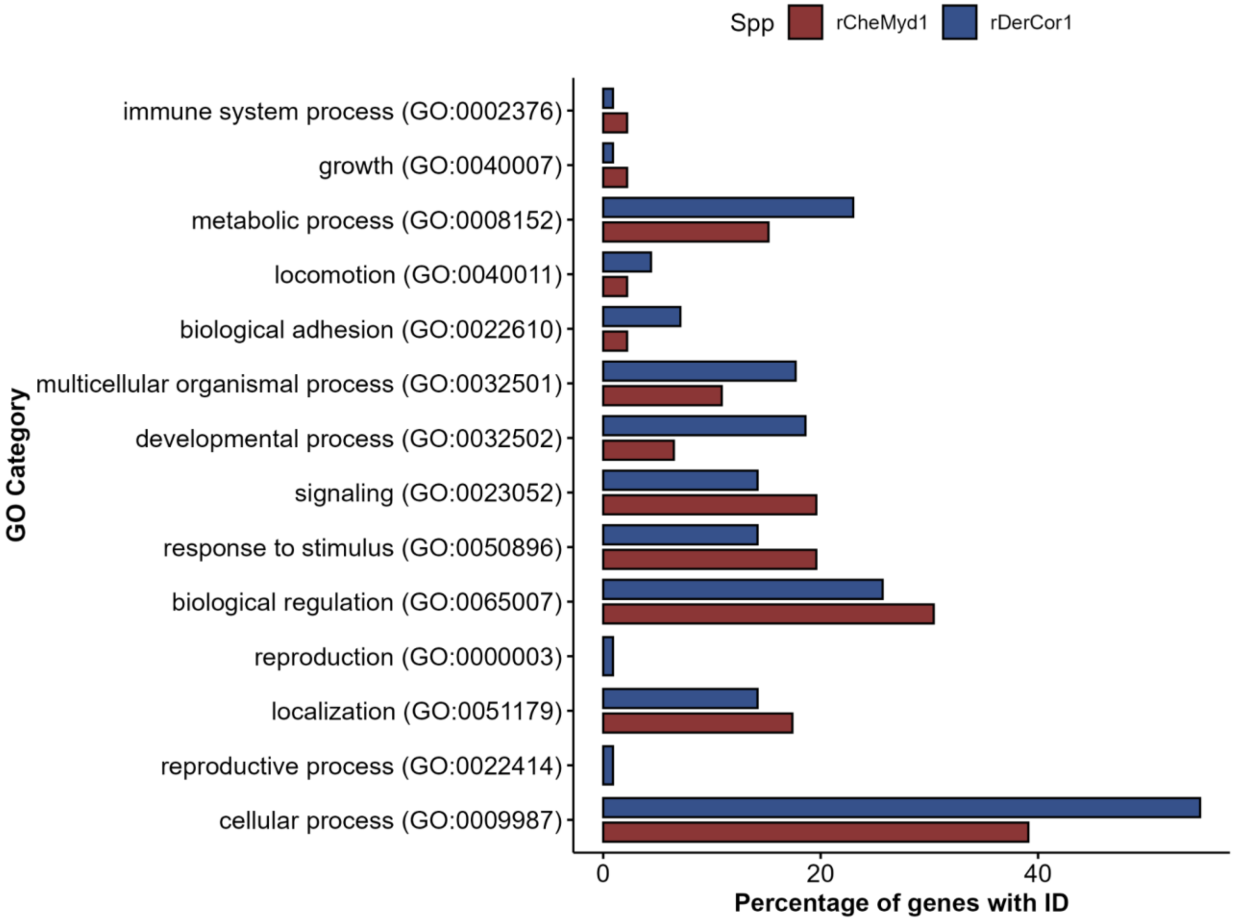


**Fig. S21 |** GO Biological Process Categories for genes identified with higher than average (mean + 3*SD) diversity in the leatherback (*Dermochelys coriacea*) and the green turtle (*Chelonia mydas*) reference individuals as predicted by PANTHER.

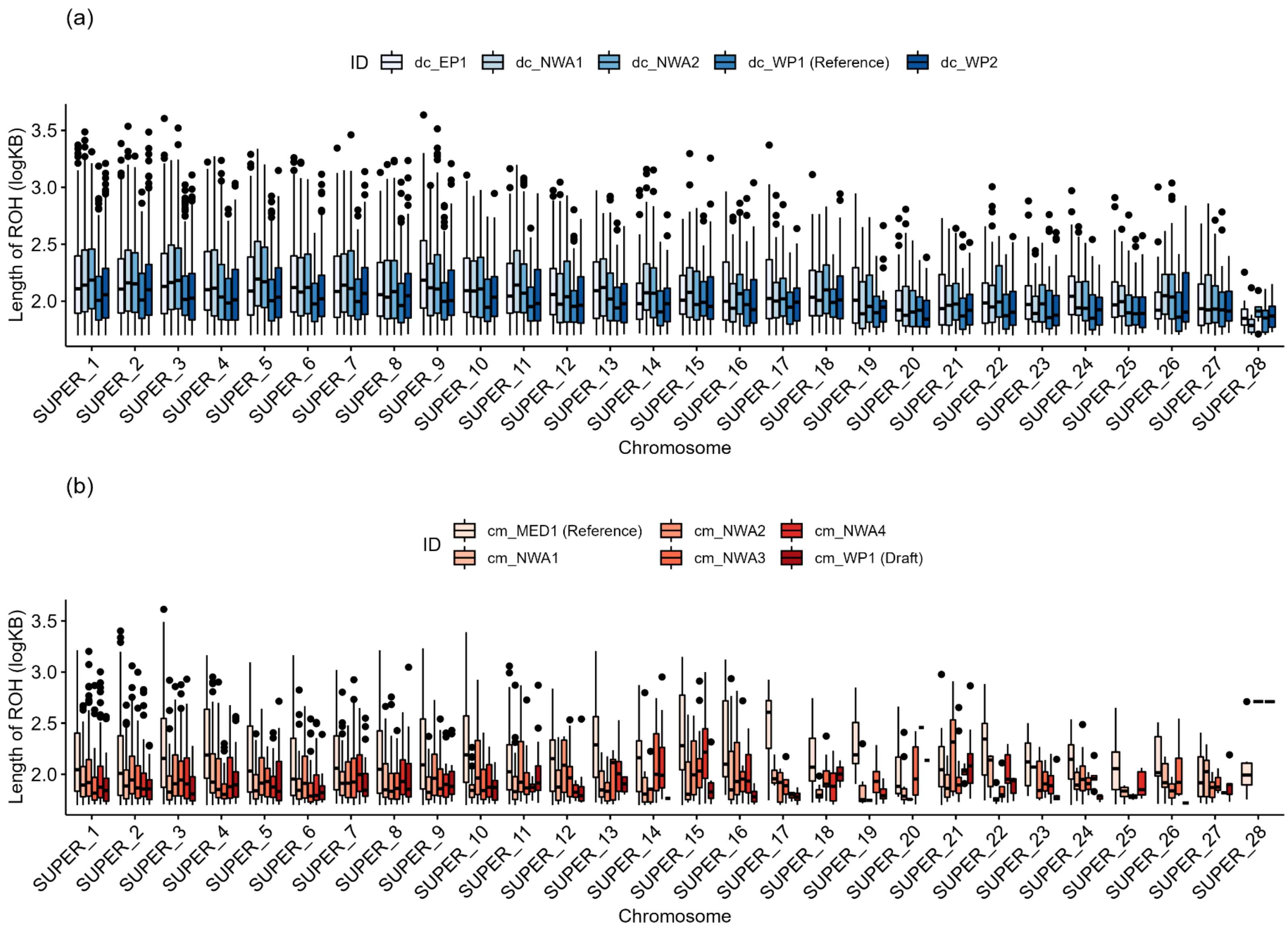


**Fig. S22** | Lengths (logKB) of runs of homozygosity (ROH) per chromosome for *Dermochelys coriacea* (a) and *Chelonia mydas* (b).

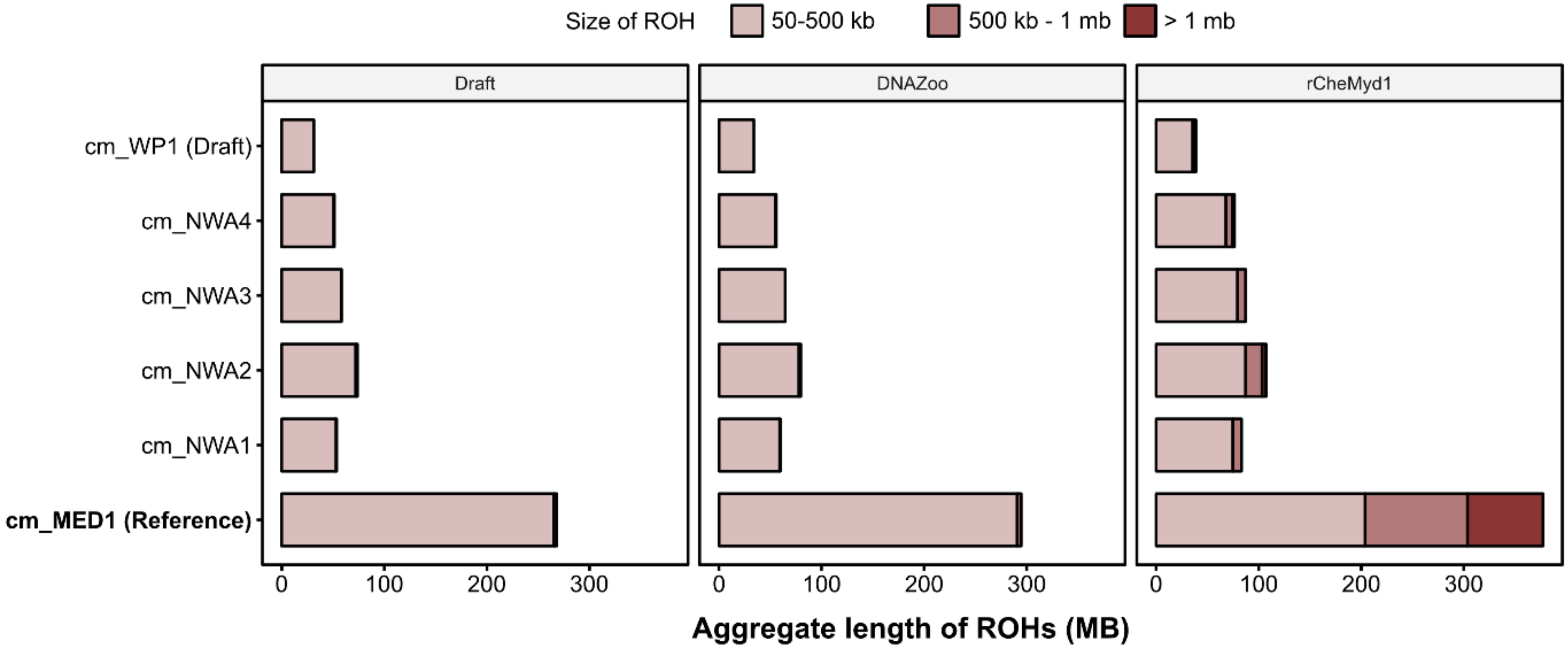


**Fig. S23** | Comparison of ROH distributions for all green turtle individuals when aligned to the draft genome, DNAZoo re-scaffolded draft genome, and our newly assembled reference genome.

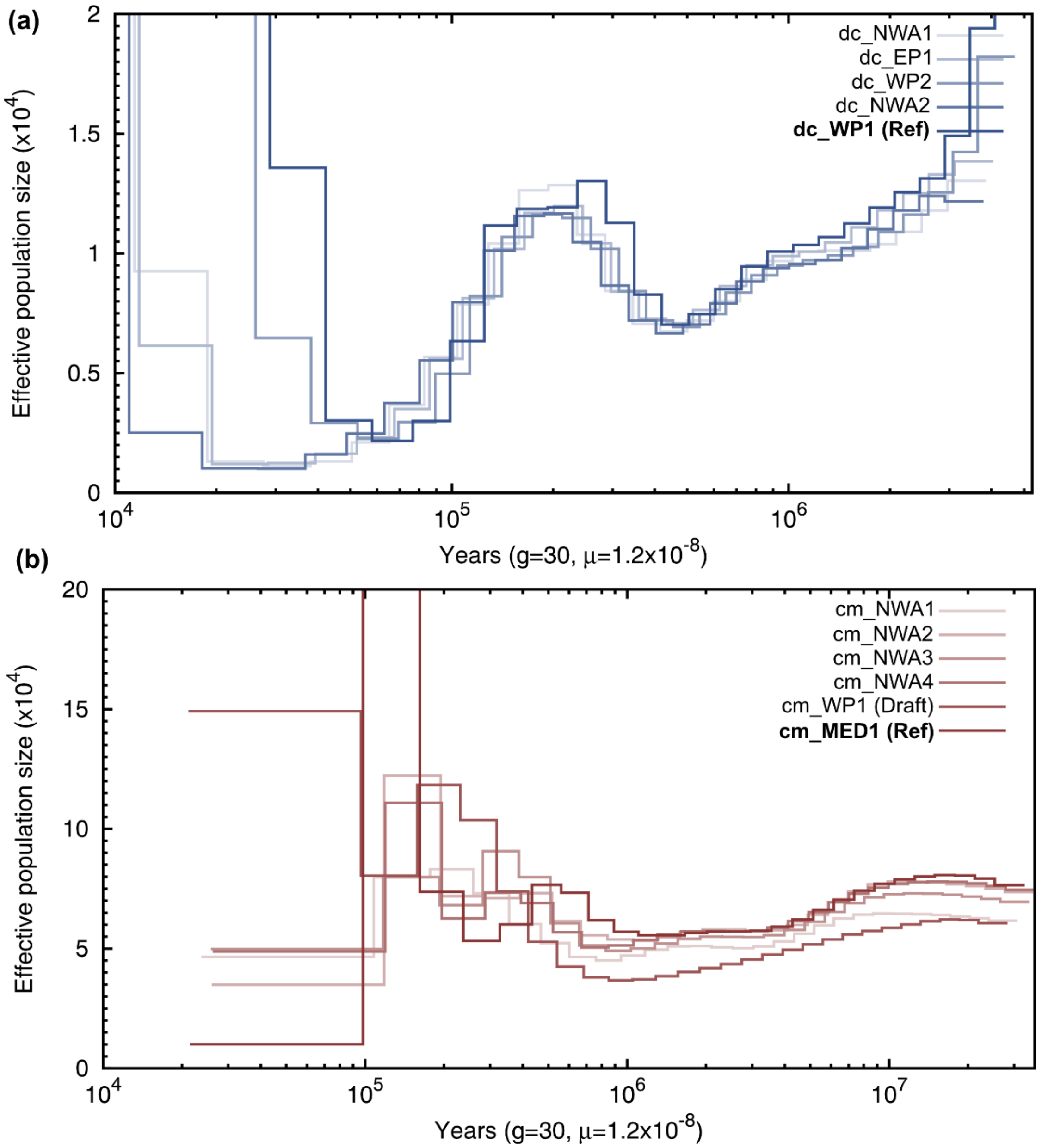


**Fig. S24** | Additional PSMC plots for *Dermochelys coriacea* (a) and *Chelonia mydas* (b) showing outputs from the additional resequenced individuals. Outputs were generated with a mutation rate of $1.2x10^{-8}$ and a generation time of 30 years for both species. Y-axes were constrained for clear visualization, with large artefactual peaks exceeding the upper limit of the Y-axes. Sample information is available in Table S13.

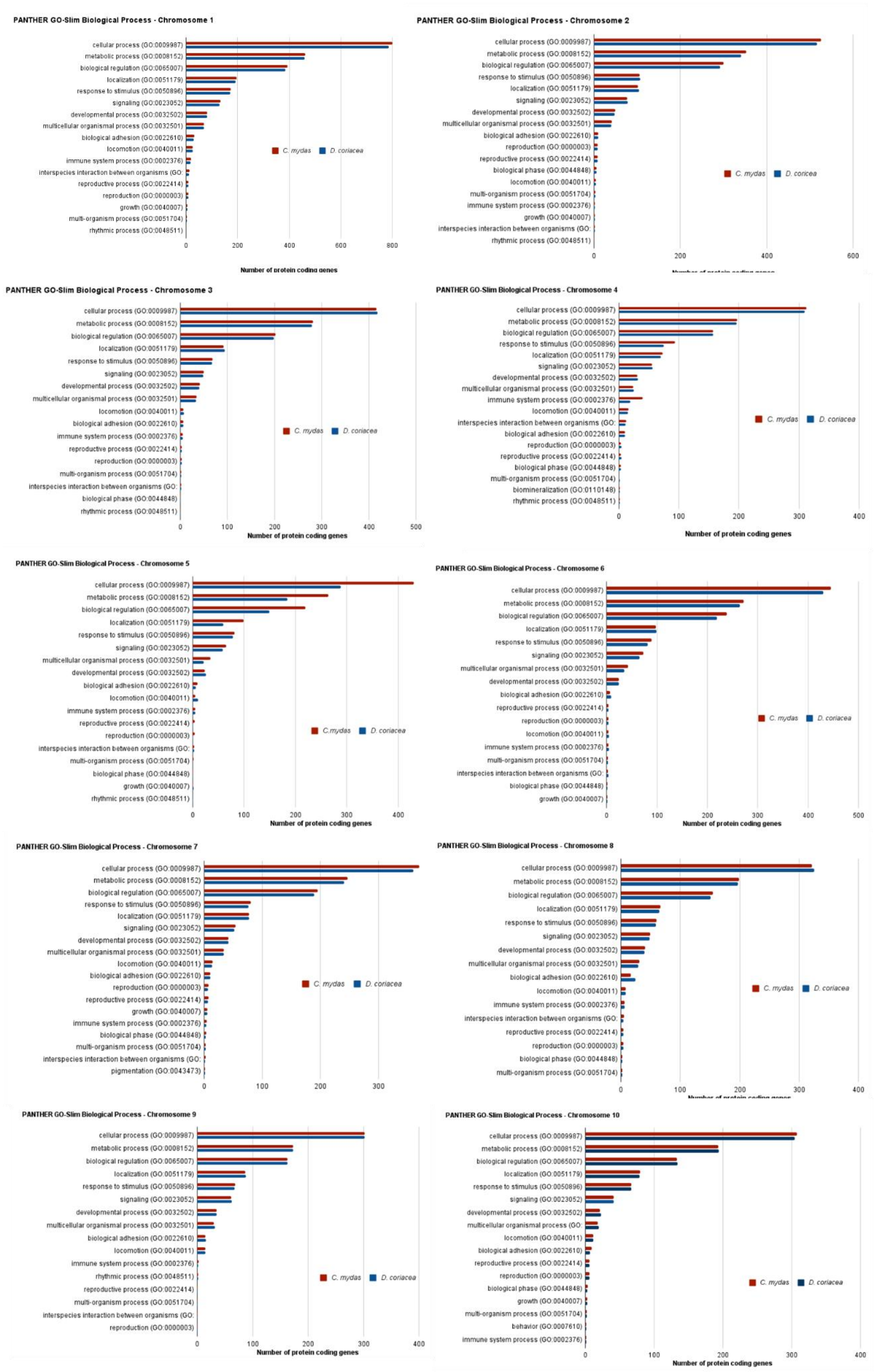

PANTHER GO-Slim Biological Process - Chromosome 1
cellular process (GO:0009987)
metabolic process (GO:0008152)
biological regulation (GO:0065007)
localization (GO:0051179)
response to stimulus (GO:0050896)
signaling (GO:0023052)
developmental process (GO:0032502)
multicellular organismal process (GO:0032501)
biological adhesion (GO:0022610)
locomotion (GO:0040011)
immune system process (GO:0002376)
interspecies interaction between organisms (GO:
reproductive process (GO:0022414)
reproduction (GO:0000003)
growth (GO:0040007)
multi-organism process (GO:0051704)
rhythmic process (GO:0048511)
C. mydas
D. coriacea
0
200
400
600
800
Number of protein coding genes
PANTHER GO-Slim Biological Process - Chromosome 2
cellular process (GO:0009987)
metabolic process (GO:0008152)
biological regulation (GO:0065007)
response to stimulus (GO:0050896)
localization (GO:0051179)
signaling (GO:0023052)
developmental process (GO:0032502)
multicellular organismal process (GO:0032501)
biological adhesion (GO:0022610)
reproduction (GO:0000003)
reproductive process (GO:0022414)
biological phase (GO:0044848)
locomotion (GO:0040011)
multi-organism process (GO:0051704)
immune system process (GO:0002376)
growth (GO:0040007)
interspecies interaction between organisms (GO:
rhythmic process (GO:0048511)
C. mydas
D. coricea
0
200
400
600
PANTHER GO-Slim Biological Process - Chromosome 3
cellular process (GO:0009987)
metabolic process (GO:0008152)
biological regulation (GO:0065007)
localization (GO:0051179)
response to stimulus (GO:0050896)
signaling (GO:0023052)
developmental process (GO:0032502)
multicellular organismal process (GO:0032501)
locomotion (GO:0040011)
biological adhesion (GO:0022610)
immune system process (GO:0002376)
reproductive process (GO:0022414)
reproduction (GO:0000003)
multi-organism process (GO:0051704)
interspecies interaction between organisms (GO:
biological phase (GO:0044848)
rhythmic process (GO:0048511)
C. mydas
D. coriacea
0
100
200
300
400
500
Number of protein coding genes
PANTHER GO-Slim Biological Process - Chromosome 4
cellular process (GO:0009987)
metabolic process (GO:0008152)
biological regulation (GO:0065007)
response to stimulus (GO:0050896)
localization (GO:0051179)
signaling (GO:0023052)
developmental process (GO:0032502)
multicellular organismal process (GO:0032501)
immune system process (GO:0002376)
locomotion (GO:0040011)
interspecies interaction between organisms (GO:
biological adhesion (GO:0022610)
reproduction (GO:0000003)
reproductive process (GO:0022414)
biological phase (GO:0044848)
multi-organism process (GO:0051704)
biomineralization (GO:0110148)
rhythmic process (GO:0048511)
C. mydas
D. coriacea
0
100
200
300
400
Number of protein coding genes
PANTHER GO-Slim Biological Process - Chromosome 5
cellular process (GO:0009987)
metabolic process (GO:0008152)
biological regulation (GO:0065007)
localization (GO:0051179)
response to stimulus (GO:0050896)
signaling (GO:0023052)
multicellular organismal process (GO:0032501)
developmental process (GO:0032502)
biological adhesion (GO:0022610)
locomotion (GO:0040011)
immune system process (GO:0002376)
reproductive process (GO:0022414)
reproduction (GO:0000003)
interspecies interaction between organisms (GO:
multi-organism process (GO:0051704)
biological phase (GO:0044848)
growth (GO:0040007)
rhythmic process (GO:0048511)
C.mydas
D. coriacea
0
100
200
300
400
Number of protein coding genes
PANTHER GO-Slim Biological Process - Chromosome 6
cellular process (GO:0009987)
metabolic process (GO:0008152)
biological regulation (GO:0065007)
localization (GO:0051179)
response to stimulus (GO:0050896)
signaling (GO:0023052)
multicellular organismal process (GO:0032501)
developmental process (GO:0032502)
biological adhesion (GO:0022610)
reproductive process (GO:0022414)
reproduction (GO:0000003)
locomotion (GO:0040011)
immune system process (GO:0002376)
multi-organism process (GO:0051704)
interspecies interaction between organisms (GO:
biological phase (GO:0044848)
growth (GO:0040007)
C. mydas
D. coriacea
0
100
200
300
400
500
Number of protein coding genes
PANTHER GO-Slim Biological Process - Chromosome 7
cellular process (GO:0009987)
metabolic process (GO:0008152)
biological regulation (GO:0065007)
response to stimulus (GO:0050896)
localization (GO:0051179)
signaling (GO:0023052)
developmental process (GO:0032502)
multicellular organismal process (GO:0032501)
locomotion (GO:0040011)
biological adhesion (GO:0022610)
reproduction (GO:0000003)
reproductive process (GO:0022414)
growth (GO:0040007)
immune system process (GO:0002376)
biological phase (GO:0044848)
multi-organism process (GO:0051704)
interspecies interaction between organisms (GO:
pigmentation (GO:0043473)
C. mydas
D. coriacea
0
100
200
300
PANTHER GO-Slim Biological Process - Chromosome 8
cellular process (GO:0009987)
metabolic process (GO:0008152)
biological regulation (GO:0065007)
localization (GO:0051179)
response to stimulus (GO:0050896)
signaling (GO:0023052)
developmental process (GO:0032502)
multicellular organismal process (GO:0032501)
biological adhesion (GO:0022610)
locomotion (GO:0040011)
immune system process (GO:0002376)
interspecies interaction between organisms (GO:
reproductive process (GO:0022414)
reproduction (GO:0000003)
biological phase (GO:0044848)
multi-organism process (GO:0051704)
C. mydas
D. coriacea
0
100
200
300
400
PANTHER GO-Slim Biological Process - Chromosome 9
cellular process (GO:0009987)
metabolic process (GO:0008152)
biological regulation (GO:0065007)
localization (GO:0051179)
response to stimulus (GO:0050896)
signaling (GO:0023052)
developmental process (GO:0032502)
multicellular organismal process (GO:0032501)
biological adhesion (GO:0022610)
locomotion (GO:0040011)
immune system process (GO:0002376)
rhythmic process (GO:0048511)
reproductive process (GO:0022414)
multi-organism process (GO:0051704)
interspecies interaction between organisms (GO:
reproduction (GO:0000003)
C. mydas
D. coriacea
0
100
200
300
400
Number of protein coding genes
PANTHER GO-Slim Biological Process - Chromosome 10
cellular process (GO:0009987)
metabolic process (GO:0008152)
biological regulation (GO:0065007)
localization (GO:0051179)
response to stimulus (GO:0050896)
signaling (GO:0023052)
developmental process (GO:0032502)
multicellular organismal process (GO:
locomotion (GO:0040011)
biological adhesion (GO:0022610)
reproductive process (GO:0022414)
reproduction (GO:0000003)
biological phase (GO:0044848)
growth (GO:0040007)
multi-organism process (GO:0051704)
behavior (GO:0007610)
immune system process (GO:0002376)
C. mydas
D. coriacea
0
100
200
300
400
Number of protein coding genes

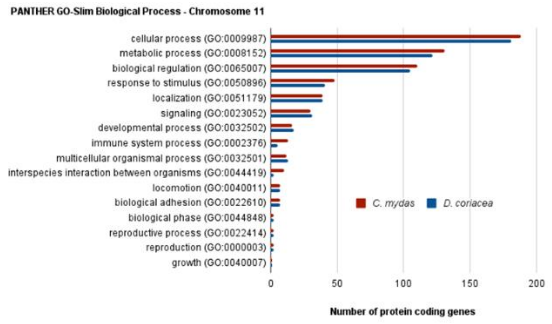
PANTHER GO-Slim Biological Process - Chromosome 11
cellular process (GO:0009987)
metabolic process (GO:0008152)
biological regulation (GO:0065007)
response to stimulus (GO:0050896)
localization (GO:0051179)
signaling (GO:0023052)
developmental process (GO:0032502)
immune system process (GO:0002376)
multicellular organismal process (GO:0032501)
interspecies interaction between organisms (GO:0044419)
locomotion (GO:0040011)
biological adhesion (GO:0022610)
biological phase (GO:0044848)
reproductive process (GO:0022414)
reproduction (GO:0000003)
growth (GO:0040007)
C. mydas
D. coriacea
0 50 100 150 200
Number of protein coding genes

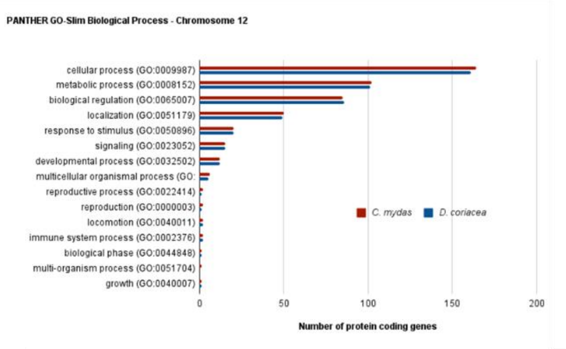
PANTHER GO-Slim Biological Process - Chromosome 12
cellular process (GO:0009987)
metabolic process (GO:0008152)
biological regulation (GO:0065007)
localization (GO:0051179)
response to stimulus (GO:0050896)
signaling (GO:0023052)
developmental process (GO:0032502)
multicellular organismal process (GO:
reproductive process (GO:0022414)
reproduction (GO:0000003)
locomotion (GO:0040011)
immune system process (GO:0002376)
biological phase (GO:0044848)
multi-organism process (GO:0051704)
growth (GO:0040007)
C. mydas
D. coriacea
0 50 100 150 200
Number of protein coding genes

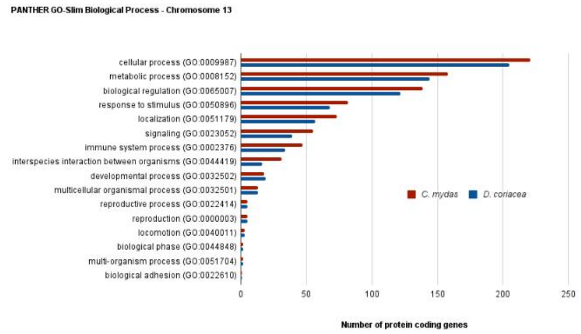
PANTHER GO-Slim Biological Process - Chromosome 13
cellular process (GO:0009987)
metabolic process (GO:0008152)
biological regulation (GO:0065007)
response to stimulus (GO:0050896)
localization (GO:0051179)
signaling (GO:0023052)
immune system process (GO:0002376)
interspecies interaction between organisms (GO:0044419)
developmental process (GO:0032502)
multicellular organismal process (GO:0032501)
reproductive process (GO:0022414)
reproduction (GO:0000003)
locomotion (GO:0040011)
biological phase (GO:0044848)
multi-organism process (GO:0051704)
biological adhesion (GO:0022610)
C. mydas
D. coriacea
0 50 100 150 200 250
Number of protein coding genes

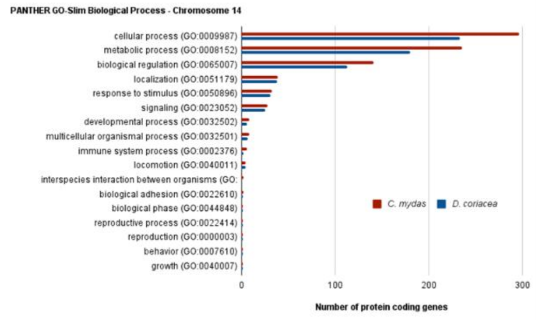
PANTHER GO-Slim Biological Process - Chromosome 14
cellular process (GO:0009987)
metabolic process (GO:0008152)
biological regulation (GO:0065007)
localization (GO:0051179)
response to stimulus (GO:0050896)
signaling (GO:0023052)
developmental process (GO:0032502)
multicellular organismal process (GO:0032501)
immune system process (GO:0002376)
locomotion (GO:0040011)
interspecies interaction between organisms (GO:
biological adhesion (GO:0022610)
biological phase (GO:0044848)
reproductive process (GO:0022414)
reproduction (GO:0000003)
behavior (GO:0007610)
growth (GO:0040007)
C. mydas
D. coriacea
0 100 200 300
Number of protein coding genes

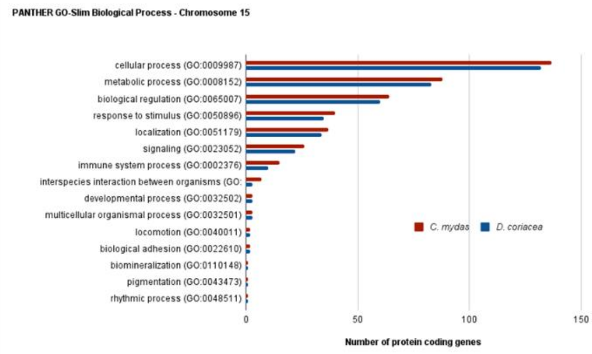
PANTHER GO-Slim Biological Process - Chromosome 15
cellular process (GO:0009987)
metabolic process (GO:0008152)
biological regulation (GO:0065007)
response to stimulus (GO:0050896)
localization (GO:0051179)
signaling (GO:0023052)
immune system process (GO:0002376)
interspecies interaction between organisms (GO:
developmental process (GO:0032502)
multicellular organismal process (GO:0032501)
locomotion (GO:0040011)
biological adhesion (GO:0022610)
biomineralization (GO:0110148)
pigmentation (GO:0043473)
rhythmic process (GO:0048511)
C. mydas
D. coriacea
0 50 100 150
Number of protein coding genes

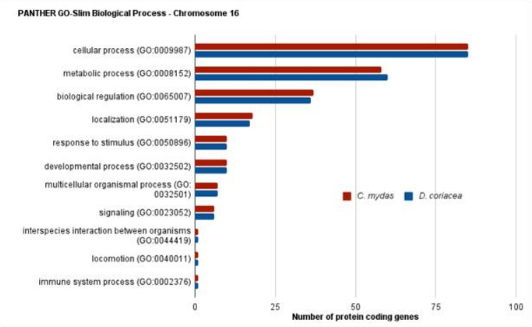
PANTHER GO-Slim Biological Process - Chromosome 16
cellular process (GO:0009987)
metabolic process (GO:0008152)
biological regulation (GO:0065007)
localization (GO:0051179)
response to stimulus (GO:0050896)
developmental process (GO:0032502)
multicellular organismal process (GO:
0032501)
signaling (GO:0023052)
interspecies interaction between organisms
(GO:0044419)
locomotion (GO:0040011)
immune system process (GO:0002376)
C. mydas
D. coriacea
0 25 50 75 100
Number of protein coding genes

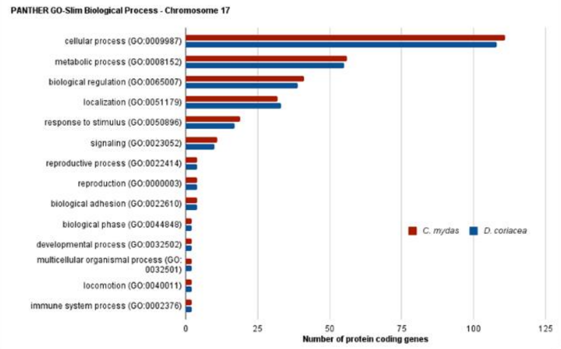
PANTHER GO-Slim Biological Process - Chromosome 17
cellular process (GO:0009987)
metabolic process (GO:0008152)
biological regulation (GO:0065007)
localization (GO:0051179)
response to stimulus (GO:0050896)
signaling (GO:0023052)
reproductive process (GO:0022414)
reproduction (GO:0000003)
biological adhesion (GO:0022610)
biological phase (GO:0044848)
developmental process (GO:0032502)
multicellular organismal process (GO:
0032501)
locomotion (GO:0040011)
immune system process (GO:0002376)
C. mydas
D. coriacea
0 25 50 75 100 125
Number of protein coding genes

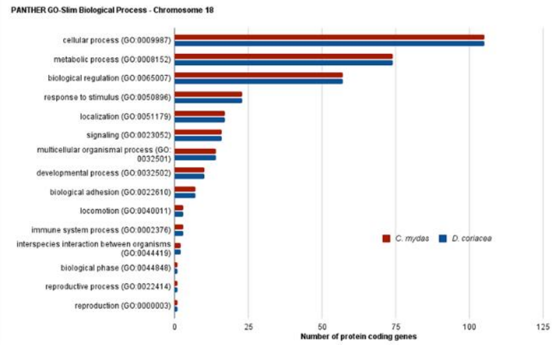
PANTHER GO-Slim Biological Process - Chromosome 18
cellular process (GO:0009987)
metabolic process (GO:0008152)
biological regulation (GO:0065007)
response to stimulus (GO:0050896)
localization (GO:0051179)
signaling (GO:0023052)
multicellular organismal process (GO:
0032501)
developmental process (GO:0032502)
biological adhesion (GO:0022610)
locomotion (GO:0040011)
immune system process (GO:0002376)
interspecies interaction between organisms
(GO:0044419)
biological phase (GO:0044848)
reproductive process (GO:0022414)
reproduction (GO:0000003)
C. mydas
D. coriacea
0 25 50 75 100 125
Number of protein coding genes

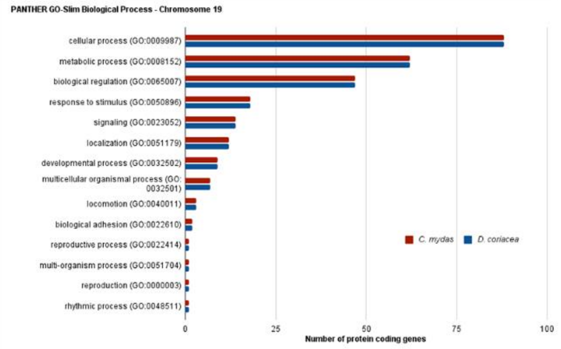
PANTHER GO-Slim Biological Process - Chromosome 19
cellular process (GO:0009987)
metabolic process (GO:0008152)
biological regulation (GO:0065007)
response to stimulus (GO:0050896)
signaling (GO:0023052)
localization (GO:0051179)
developmental process (GO:0032502)
multicellular organismal process (GO:
0032501)
locomotion (GO:0040011)
biological adhesion (GO:0022610)
reproductive process (GO:0022414)
multi-organism process (GO:0051704)
reproduction (GO:0000003)
rhythmic process (GO:0048511)
C. mydas
D. coriacea
0 25 50 75 100
Number of protein coding genes

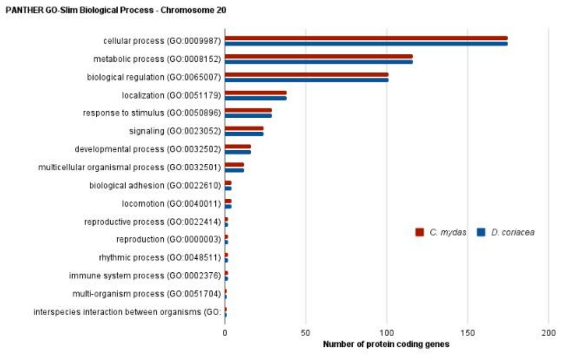
PANTHER GO-Slim Biological Process - Chromosome 20
cellular process (GO:0009987)
metabolic process (GO:0008152)
biological regulation (GO:0065007)
localization (GO:0051179)
response to stimulus (GO:0050896)
signaling (GO:0023052)
developmental process (GO:0032502)
multicellular organismal process (GO:0032501)
biological adhesion (GO:0022610)
locomotion (GO:0040011)
reproductive process (GO:0022414)
reproduction (GO:0000003)
rhythmic process (GO:0048511)
immune system process (GO:0002376)
multi-organism process (GO:0051704)
interspecies interaction between organisms (GO:
C. mydas
D. coriacea
0 50 100 150 200
Number of protein coding genes

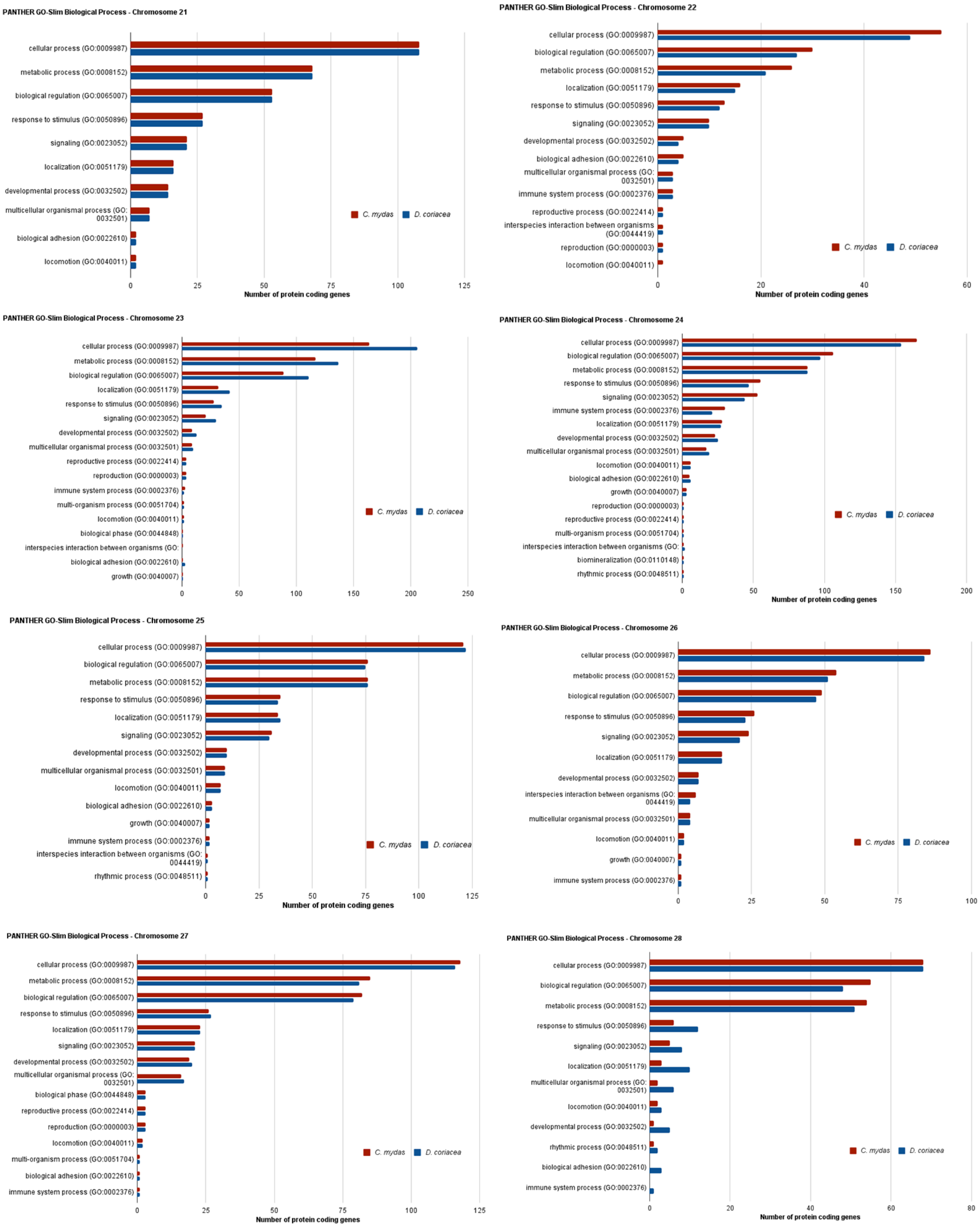


**Fig. S25 |** PANTHER GO-slim classification by biological process of the coding sequences present in each chromosome for *Chelonia mydas* and *Dermochelys coriacea*.

54. R Core Team, *R: A language and environment for statistical computing* (R Foundation for Statistical

Computing, 2020).
